# Quantification of single molecule glycan-mannose receptor binding kinetics on myeloid cells reveals high subcellular binding heterogeneity

**DOI:** 10.1101/2025.01.24.634682

**Authors:** Kas Steuten, Johannes J.A. Bakker, Ward Doelman, Diana Torres-García, Roger Riera, Lorenzo Albertazzi, Sander I. van Kasteren

## Abstract

Extracting single-molecule lectin binding kinetics from primary cells has not been possible to date. Here, we present **Glyco-PAINT-APP** (Automated Processing Pipeline), an automated method that enables the extraction of subcellular glycan interaction kinetics using a Points Accumulation for Imaging in Nano-Topography (PAINT)-based approach. This approach leverages an algorithm for precise, high-throughput subcellular analysis of glycan binding dynamics, facilitating the exploration of functional correlations between glycoform binding patterns and immune cell polarization. Using synthetic glycans and glycosylated antigens, we demonstrate the ability of the technique to automatically correlate glycan binding parameters in subregions of dendritic cell membranes with increased uptake and cross-presentation efficiency of these antigens. Additionally, we show how the method can uncover subtle differences in glycan binding preferences between resting and polarized macrophages. Taken together, Glyco-PAINT-APP has the potential to enable new insights into the cell-intrinsic heterogeneity of glycan-structure-activity relationships in immune cells.

## Introduction

Myeloid immune cells such as dendritic cells and macrophages are key sentinels of the immune system. They sense the presence of microbial species or altered self through pattern recognition receptors (PRRs), which in turn can trigger innate and adaptive immune responses. One important family of PRRs are the carbohydrate-binding proteins of the C-type lectin receptor family (CLRs).^1,2^ CLR ligation can induce various immune functions, including endocytosis, polarization, cytokine secretion, cell adhesion, and motility, which can subsequently trigger downstream immune responses.^1,3^

The signalling pathways leading to these biological outcomes are intricate and finely tuned, which complicates the study of how the emit signals upon ligand recognition, particularly in their native environment on the cell surface. This complexity arises from a variety of factors: binding of the receptors to ligands is weak and often multivalent, multiple receptors have overlapping specificities, and the surface expression patterns of CLRs can be highly dynamic. Myeloid cells express a diverse array of CLRs for sensing aberrant glycosylation. These can exists as monomers, multimers, and even form heterodimeric complexes^4,5^. Various factors, such as the polarization state of the cell modulates the expression levels of these lectins, as well as the cycling of these lectins between the surface and intracellular storage pools.^6,7^ The resulting dynamic and heterogeneous surface expression leads to significant variability in receptor-ligand interactions.

To complicate matters further, myeloid cells also present a repertoire of glycans that serve as ligands for their lectins that can modulate signalling pathways through ligand competition, and receptor clustering. Polarization also influences the expression of these cis-ligands, adding yet another layer of complexity.^8,9^ The final complicating factor in CLR-binding and signalling is that the affinities of CTLs for their ligands are weak (K_D_ in µM to mM range)^10^, which complicates the detection and precise study of these interactions. Together with the intrinsic chemical complexity of glycan ligands, these factors have made it exceedingly difficult to establish clear relationships between glycan binding properties and lectin function, particularly as studying these lectins out of their native context also removes these modulating factors.

We have recently reported a technique for quantifying carbohydrate-lectin binding kinetics on cell lines called Glyco-PAINT.^11^ Glyco-PAINT exploits the transient interaction between fluorescently-labelled sugars and their receptors to generate single molecule binding events^11^, analogous to those induced by the mismatches of DNA-strands in conventional Points Accumulation for Imaging in Nanoscale Topography (DNA-PAINT)-imaging.^12^ Accumulation of these binding events over time allows for a complete mapping of receptor-ligand interactions on the cell surface and subsequent derivatization of kinetic parameters such as on-rate, off-rate and diffusion coefficients on a per-cell basis derived from single-molecule binding information.

We envisaged that Glyco-PAINT could be a powerful technique to unravel the complexity of myeloid lectin binding, as the single molecule as well as the live cell aspects of the technique are ideal for studying these capricious receptors within their native environment on the myeloid cell surface. However, three major hurdles needed to be overcome to achieve this. The first is that Glyco-PAINT had only been demonstrated in an over-expression system. Second, the Glyco-PAINT processing pipeline could only be used to obtain the binding parameters averaged over the surface of a whole cell, which we hypothesized to be insufficient, due to the microheterogeneity of receptor expression on primary cells. The final problem relates to the probes: the Glyco-PAINT reagents consist of a carbohydrate cluster directly attached to a fluorophore, which minimises the downstream immunological evaluation of receptor biology, therefore precluding the correlation of binding to biological function.

One receptor which is archetypal in its and biological complexity is the mannose receptor (CD206, MR). This lectin is expressed on cells of the myeloid lineage^13–15^, where it carries out multiple functions: as an endocytic receptor to clear glycoproteins, as a phagocytic receptor binding to, and internalizing, glycosylated particles for clearance and antigen presentation^16–18^, and as a mediator of inflammatory signalling by inducing M2 polarization of macrophages and inducing T cell tolerance.^19,20^ How the MR translates ligand recognition to all these functions remains unknown. Structurally, the MR contains multiple functional domains, including an N-terminal cysteine-rich domain capable of binding sulphated glycans (that can inhibit receptor function^21^), a fibronectin-like domain capable of binding collagen peptides^22,23^, and eight C-type lectin domains (CTLs), of which only CTL4 and CTL5 exhibit functional calcium-dependent binding of neutral sugars.^24^ These domains are further modulated by glycosylation at seven N-linked sites, influencing receptor binding properties.^25^ Moreover, the intracellular tail lacks classical signalling motifs ^6,16^, though recent evidence suggests phosphorylation and diaromatic motifs may facilitate endosomal sorting and signalling.^26–28^

Despite its biological complexity, MR has been under heavy investigation as vaccine receptor. Ligation of the MR was shown to lead to the enhanced cross-presentation (and thus cytotoxic CD8 T-cell activation) of glycoprotein antigens, which can be employed to enhance the anti-cancer and anti-viral properties of therapeutic vaccines.^29^ However, further pursuit of this phenomenon has led to conflicting results.^30–38^ No studies have yet managed to correlate the binding of ligands to the MR to downstream biology (i.e., uptake and/or cross-presentation). Studies using bulk methods, such as surface plasmon resonance (SPR), failed to correlate binding affinities with uptake or cross-presentation, highlighting the need for methodologies capable of quantifying receptor-ligand kinetics in the native cellular context.^11,31^

Here, we present the redevelopment of Glyco-PAINT, such that we can obtain binding parameters of glycans to the MR on myeloid cells. We have developed a new analysis algorithm to independently process large volumes of Glyco-PAINT recordings of glycan binding events to primary immune cells in such a way that binding event concentration, dwell times, diffusion coefficients and displacement can be determined for the highly heterogeneous and disperse binding observed across the cell basal membrane of dendritic cells and macrophages. We have dubbed this method “Glyco-PAINT-Automated Processing Pipeline” (Glyco-PAINT-APP). In addition, we developed a new probe library containing a pendant cross-presentable epitope that allowed us to quantify lectin binding kinetics, uptake, and cross-presentation of the same construct. Correlating the binding kinetics to uptake and cross-presentation biology yielded surprising findings. The first of these was, as described above, that binding events are not homogeneously distributed across the DC or macrophage membrane.

In addition – using the novel segmentation approach – we found a strong negative correlation (Pearson’s R: 0.90) between the cell surface mobility of ligands to subsequent endocytosis by the DC. In contrast, cross-presentation of mannosylated antigens showed a strong positive correlation to the residence time on the DC (Pearson’s R: 0.94) independent of the amount of surface mobility. In a second application of the approach, we could also quantify changes in mannoside binding during the macrophage polarisation trajectory from M0 to M2, but not M1.

## Results

### Quantifying glycan-lectin binding on primary DC

Our first aim was to determine whether the Glyco-PAINT methodology could be used to study binding events on the surface of primary bone-marrow derived mouse dendritic cells (BMDCs).^39^ Unlike the overexpression system used previously, BMDCs suffer from heterogeneous and dynamic receptor expression, broad lectin repertoires, and structural features such as tight focal adhesions and podosomes that may exclude binding events from the TIRF plane.^40,41^

Preliminary attempts to quantify binding kinetics of a labelled mannose glycan library on BMDCs yielded a surprising result: whereas binding events could clearly be seen in the Glyco-PAINT recordings (**Figure 1B** and **Figure S1A-D**) no significant differences were observed between the probes for either the events µm-^2^sec^-1^ (a parameter that correlates to k_on*_[receptor concentration]), nor in the average ligand dwell time (**τ**, which is the inverse of the off rate, or k_off_^-1^) (**Figure S1E-F**). Control experiments in CHO-MR cells did show the binding behaviour as originally reported (**Figure 1A** and **Figure S2A-D** and **Figure S2E, F** for quantification).

**Figure 1.**
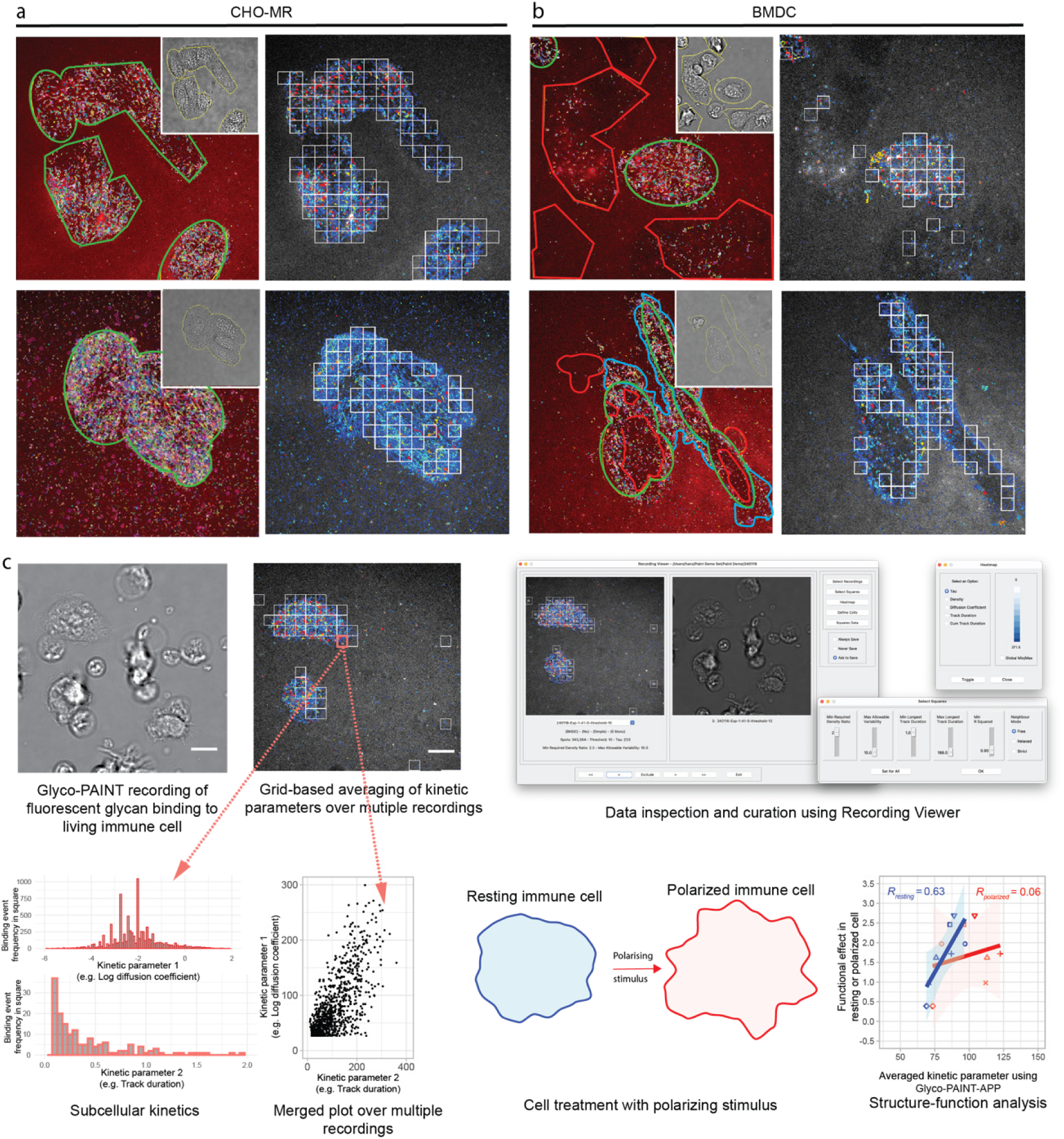
Glyco-PAINT-APP allows subcellular analysis of immune cell lectin binding. **A)** Left images: original Glyco-PAINT analysis of fluorescent mannose glycan binding to CHO-MR cells. ROIs are drawn using brightfield-defined cell contours (inset, yellow outlines) and areas where binding events are overlapping with cell areas are outlined in green. Binding trajectories are randomly coloured. Right: Grid-based identification of binding events using Glyco-PAINT-APP. White-outlined squares show an independent selection of cell-related binding events and define the area over which events are integrated. **B)** Same as **A** but for BMDC. In the left images, non-overlapping areas devoid of binding events arising from primary cell intrinsic heterogeneity are outlined in red and areas engaging in binding events but outside of brightfield-defined cell outlines are coloured in blue. **C)** Overview of the Glyco-PAINT-APP workflow: primary immune cells are cultured in a live-cell imaging chamber and a brightfield image is recorded. After addition of a fluorescent probe, spots are recorded using PAINT imaging and reconstructed into tracks reflecting glycan binding events which are then subdivided into squares of variable size depicting subcellular regions of live immune cells. Kinetic parameters such as on-, off-rates and diffusion coefficients of glycan ligands averaged per individual square can then be analyzed and filtered using the *Recording Viewe*r tool and displayed as heatmaps, histograms or scatterplots. Kinetic parameters derived using the algorithm can then be correlated to functional effects of resting or polarized immune cells. Scalebars represent 10 µm, data figures indicate representative illustrative data

Interestingly, density maps did reveal a stark heterogeneity of binding events, both between cells – with some cells devoid of binding events, and other showing large numbers of events – but also between different regions of the basal membrane of a single BMDC. Areas of high binding events were observed overlapping with brightfield-defined cell outlines (**Figure 1B** green areas and **Figure S1A-D**). However, large areas of the cell basal membrane were also devoid of any binding event density (**Figure 1B** red areas and **Figure S1A-D**). Areas outside the brightfield-defined cell surface appeared to be also engaging in binding events (**Figure 1B**, blue areas), suggesting the presence of receptors on ultrathin dendritic protrusions. This heterogeneity was not observed for CHO-MR cells (**Figure 1A**, green areas and **Figure S2**).

These observations meant that the fundamental assumption underpinning the Glyco-PAINT method – that the receptor concentration [R] is homogeneous over the whole basal membrane – was incorrect. This in turn implicates that the per-cell relative on-rate was an irrelevant parameter for the study of primary cells. The off-rates too are averaged on a per-cell basis as they are integrated over the complete area only a subsection contains binding-enabled lectins. Taken together, the observed highly heterogeneous binding event distribution, therefore renders the assumption of a constant and homogeneous receptor concentration incorrect and calls for a more dynamic surface area definition as to obtain meaningful quantification of kinetic parameters.

### Glyco-PAINT-APP

To address the heterogeneity observed on BMDC surfaces, we developed the Glyco-PAINT-Automated Processing Pipeline (Glyco-PAINT-APP). Instead of using the whole cell as the averaging unit, subcellular regions are selected. This allows precise quantification of glycan binding kinetics within localized areas. This new workflow has fully automated every step of the Glyco-PAINT analysis method, accelerating analysis speed. The Glyco-PAINT-APP is schematically outlined in **Figure 1C** (and described in full detail in the **Supplementary Manual**).

Glyco-PAINT-APP starts with subjecting sets of microscope recordings representing individual Glyco-PAINT experiments to single particle tracking analysis (see also **Supplementary Video 1**). This yields output files containing spatial information belonging to each spot and the resulting tracks formed by connecting closely linked spots through multiple consecutive frames. We then apply a new processing algorithm to these data where, instead of manual cell definition by ROI drawing, a raster of squares (typically 20×20 squares per field of view) is laid over the image. Kinetic information is then averaged per square, rather than per cell. Quality control parameters such as the *Density Ratio*, *Maximum Variability* and *Connectivity* can be retrieved per square to accurately discriminate signal from background. *Density Ratio* is a measure for signal-to-background (by setting background levels as the 10% squares with lowest event density but containing at least one event), *Maximum Variability* is a heterogeneity score for event distribution within a square and *Connectivity* is a measure for squares being surrounded by adjacent squares or not. Further selection takes place via statistical parameters such as the *R Squared* value that relates to the quality of the one-phase exponential decay fit or minimum number of binding events (see **Supplementary Manual** for a detailed description of quality control parameters). The interactive tuning of these parameters using the *Select Squares* window together with visual feedback from the *Recording Viewer* (**Figure 1C** and **Supplementary Video 2**) where the tracking overlay with grids and brightfield image are displayed side by side allows for user-informed data curation. Kinetic parameters including glycan on- and off-rates, diffusion coefficients, speed, displacement and maximum, average, or total binding event duration can be displayed for each square per recording using the heatmap toggle function in the *Recording Viewer* (**Figure 1C**). The *Compile Project* built-in modality can integrate kinetic data from multiple recordings obtained from multiple biological replicates within a single project and can generate a merged output file to which statistical analysis can be applied with scientific graphing software of choice (R Studio^43^, GraphPad Prism, etc.), yielding a suitable method for the study of biological systems with the required biological replicates.

### Subcellular analysis of immune cell lectin binding

We then applied Glyco-PAINT-APP to the profiling of glycan binding events on DC. This required us to first optimize the grid size. For this we used the complete dataset of mannose ligands binding to BMDC, which consists of approximately 300 different recordings (each of standard 81 x 81 µm², 512 x 512 pixels, and 2000 frames at 20 fps). Our analysis demonstrated that 20 squares per axis provided an optimal balance between statistical quality, the number of binding events, and the minimization of averaging artifacts (**Figure S5**). *R Squared* distribution remained in the 0.7-1 range for the chosen grid size. Off-rates (**τ)** and density (rel. k_on_) values diverged, reflecting enhanced sensitivity for measurement of inherently heterogeneous systems.

With this new analysis pipeline in hand, we set out to reprocess the recordings of the mannose probe library on primary DCs for which the original Glyco-PAINT procedure had not yielded any significant differences in binding between the different glycoforms. The square-based analysis for DCs is displayed in **Figure 2** (for the same recordings as in **Figure S1**) and extended in **Figure S3** for mono- and trimannosides **8**, **10**, **11** and **13**. From **Figure 2B and F**, it can be seen that the new algorithm was able to independently distinguish areas with binding event density from background using filtering based on uniform parameters for the entire experiment. The added advantage of this approach is that binding events that fell outside brightfield-invisible features of the DC, such as on the dendrites that lend the cells their name, are now also included in the analysis. The thus observed binding rates in different areas of the cell basal membrane can be visualised as heatmaps in **Figure 2C, D** and **G, H**. The grid-based analysis by Glyco-PAINT-APP even improves the effective selection of cell areas engaged in binding events for mono- and trimannoside-based probe binding to CHO-MR (**Figure S4**), highlighting the omnipresence of microheterogeneity even in overexpression systems. Taken together, this method allows for precise analysis of glycan-lectin interactions on live immune cells with intrinsically heterogenic expression avoiding the emergence of averaging artefacts as in the original Glyco-PAINT analysis. In addition, some interesting parameters began to emerge. For example, a discrepancy between areas of high number of binding events (i.e., high [R]*k_on_; **Figure 2C, G**) and areas with long average duration of binding (**τ**, k_off_^-1^, **Figure 2D, H**) within the same cell.

**Figure 2.**
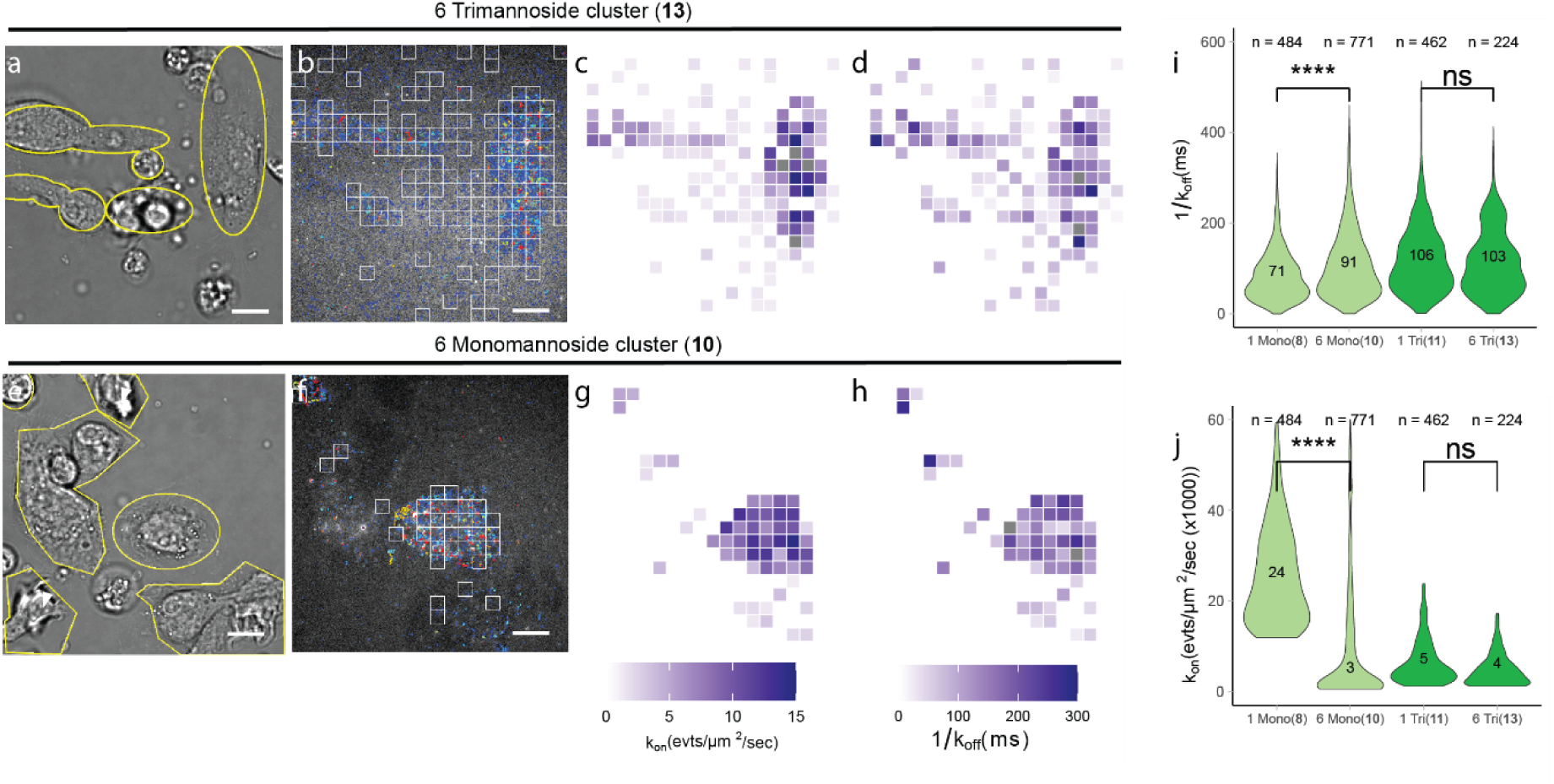
Subcellular analysis of glycan binding on dendritic cells resolves averaging artefacts. **A)** brightfield image of live BMDCs. **B)** Reconstruction of binding events recorded with 5 nM trimannoside glycan **13** processed using a 20×20 grid with all squares that have passed selection criteria (*Density_Ratio* > 2, *R_Squared* > 0.9, *Nr_Tracks/square* > 20) displayed by white outlines (**Figure S3A, B** show the same set of images for glycan clusters **11** and **13**). **C)** Heatmap projection of the number of binding events/square showing the high heterogeneity of binding events even between the selected squares on the same cell. **D)** Heatmap display of τ per square. **E-H)** The same as **A-D** but for glycan cluster **10**. **I-J)** Violin distribution plots and median value of k_on_ and k_off_^-1^ for all glycans **10-13**. n = the number of squares that were analyzed per violin. Significance was assessed using two-way ANOVA followed by a Tukey post-hoc test. Scalebars represent 10 µm. Tracks in **B** and **F** are colored from short to long track length (light blue to red).

### Uptake and antigen presentation of SLP glycoforms

We next wanted to determine whether any of the on-cell kinetic parameters could prove predictive for some of the previously reported roles (see introduction) of the MR and related CLRs on BMDCs. We therefore designed Synthetic Long Peptide (SLP) versions of the fluorophore-labelled mannose clusters that also contained the model cross-presentable epitope Ovalbumin_247-264_ (OVA SLP)^44^, as it has been hypothesized that different glycoforms of these SLPs show different cross-presentation behaviour.^34,45^ This additional feature would allow us to not only monitor the various binding parameters (by Glyco-PAINT-APP), but also uptake (by flow cytometry) and antigen cross-presentation (using the cognate OT-I T-cell), thereby having readouts for these three aspects of MR biology from a single probe.

The constructs were synthesized using in-line solid-phase peptide synthesis of the peptide antigen and an oligo-azidolysine (6-azidonorleucine) cluster, followed by on-resin modification with fluorophore. Copper-catalysed azide-alkyne cycloaddition (CuAAC)-mediated^46,47^ coupling of the mannose glycans to the azidolysine clusters in solution yielded the target glycosylated SLPs. This resulted in mono-, bi- and hexavalent versions of mono-(**1-3**) or trimannosylated (**4-6**) SLPs and a non-glycosylated control molecule (**7**) that are displayed in **Figure 3A** (synthetic procedures and characterization can be found in the supplementary information and **Figure S6**). These glycan motifs span a K_D_ range for MR-binding from 3 µM to more than 100 µM (based on SPR studies^11,31^), thereby enabling investigation of glycan-structure-binding-activity relationships over a large affinity range.

**Figure 3.**
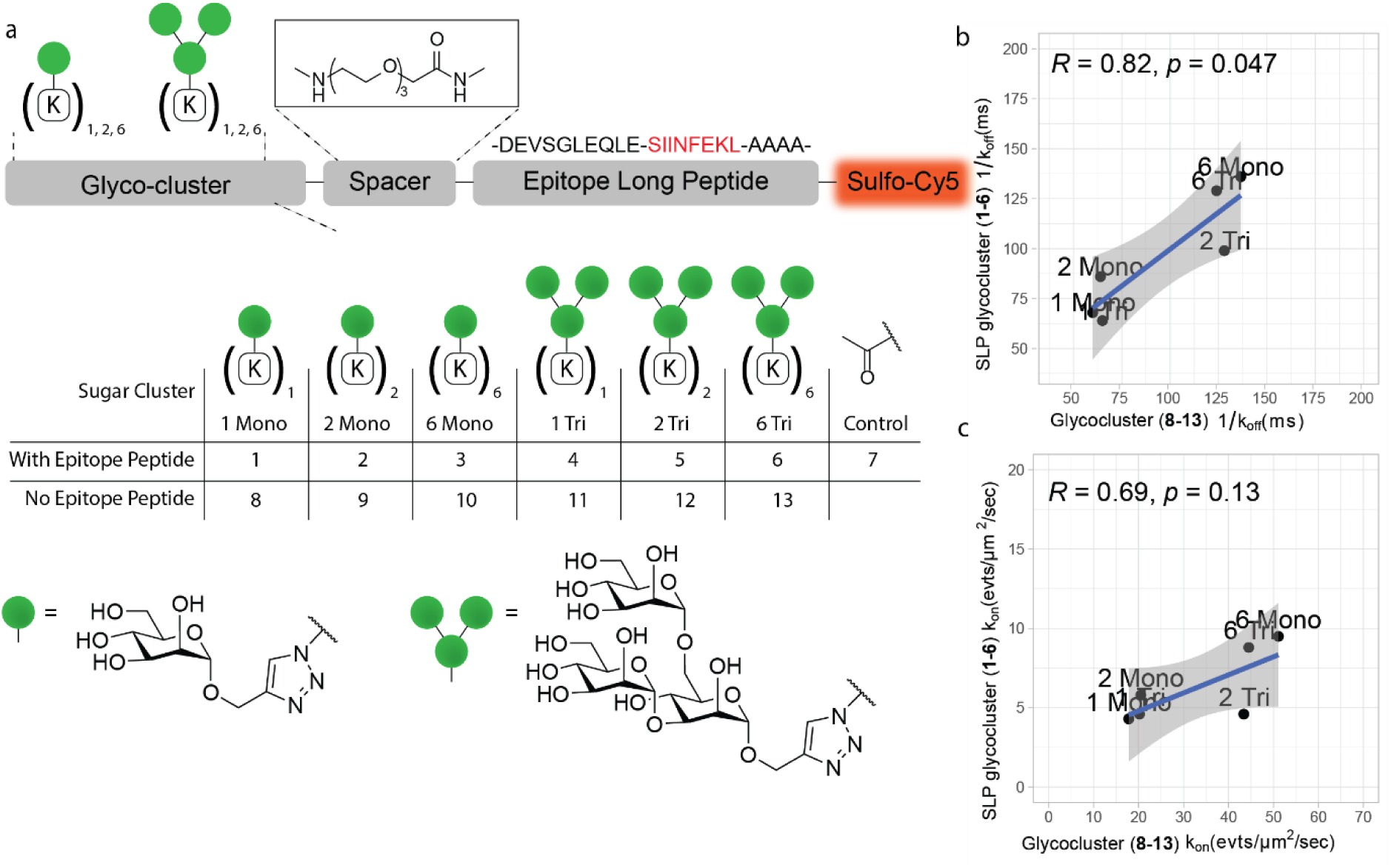
Library of fluorescently labeled mannose ligands with SLP antigenic peptide. **A)** Design of synthetic glycoclusters. **1-7** carry an N-terminal long peptide from Ovalbumin_263-275_ and a sulfo-Cy5 fluorophore on a C-terminal lysine side chain. Glycocluster series **8-13** do not contain this SLP, but are directly attached to an ATTO655 fluorophore. **B-C)** Correlation analysis of the median rel. k_on_ and k_off_^-1^ between probes with identical glycan but bearing long peptide antigens binding to CHO-MR cells. Data points are medians for all squares analyzed with a 20×20 grid and filtered with *Density_Ratio* > 2, *R_Squared* > 0.9 and *Nr_Tracks*/*Square* > 20. Pearson’s R and two-tailed t test P value are displayed in the correlation plots.

To verify the absence of artefacts resulting from the SLP antigenic cargo on kinetics we compared the median on- and off-rates of OVA SLP **1-6** to glycoclusters **8-13** on CHO-MR cells and plotted the correlation between the two in **Figure 3B, C**. Neither extension of the peptide core, nor the change of the fluorophore from ATTO655 to sulfo-Cy5 significantly affected k_off_. A decrease in the median number of binding events per square was detected for the glycan SLPs with respect to the glycans **8-13**, potentially originating from modestly altered physical-chemical properties of the peptide cargo.

The binding parameters of these glycosylated SLPs to live DC were then determined using the Glyco-PAINT-APP. To prevent the termination of binding events by endocytosis – which is one of the biological functions resulting from MR-ligation – we also performed kinetic measurements after 30 minutes treatment of the cells with Cytochalasin D (CytD), an inhibitor of actin polymerization.^48,49^ This inhibitor is known to inhibit many endocytic pathways that depend on F-actin filament polymerisation. We obtained off-rates and diffusion coefficients per square for **1-7** after CytD addition (**Figure 4A, B**, and **Table S1**) and compared glycoform-specific trends to those of non-CytD-treated cells. On-rates, off-rates and diffusion coefficients were significantly increased for all glycosylated probes **1-6** after DC and CHO-MR treatment with Cytochalasin D (**Figure 4A, B** and **Table S1**, and **Figure S7)** but not for the non-glycosylated control antigen **7**. Confirming the interactions for these probes being glycan-lectin-mediated, particularly those that depend on actin polymerization and rearrangement of the cytoskeleton for endocytosis.

**Figure 4.**
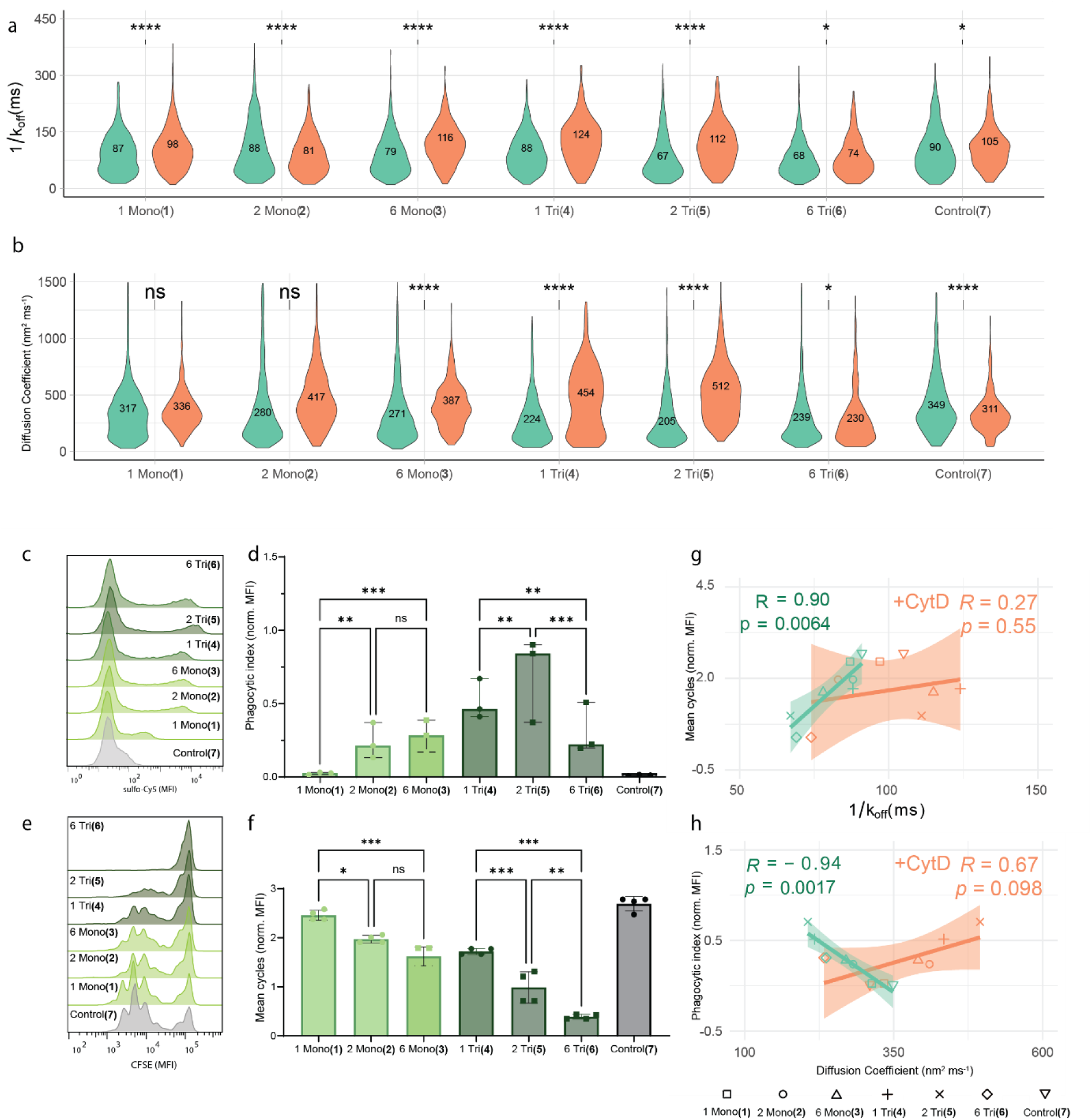
Kinetic parameters correlate with ligand functionality. **A)** Distribution of k_off_^-1^ for binding events between probes **1-7** and live untreated (green violins) or Cytochalasin D treated (red violins) BMDCs. Numerical median values are displayed in the violins. **B)** Same as in **A)** for the average diffusion coefficient per square. **C)** Histograms of BMDCs incubated with probe **1-7** for 1h and subsequent measurement of endocytosis in CD11c^+^ cells using flow cytometry. **D)** Bar chart of **C)** as the phagocytic index. **E)** Histograms of CFSE dilution in dividing OT-I T cells that were cocultured for 3 days with mature BMDC that were pulsed for 2h with antigens **1-7** and measured using flow cytometry. **F)** Bar chart of the average number of cell divisions per T cell obtained from **E)**. **G)** Correlation between median k_off_^-1^ for untreated or CytD treated cells and the mean number of T-cell divisions. **H)** Correlation between probe median diffusion coefficients for untreated or CytD treated cells and phagocytic index. Glyco-PAINT-APP analysis was done using a 20×20 grid and filtered with *Density_Ratio* > 2, *R_Squared* > 0.9 and *Nr_Tracks* > 20. Significance was assessed using two-way ANOVA followed by a Tukey post-hoc test. Pearson’s R and two-tailed t test P value are displayed in the correlation plots.

Next, the uptake of these SLPs was measured by incubating BMDCs with SLP **1-7** for 1h, followed by quantification of the sCy5 signal by flow cytometry. The histograms in **Figure 4C** show that a specific subpopulation of MR^+^CD11c^+^ DCs (**Figure S8B**, for MR co-staining) are mostly engaged in this MR-mediated uptake (**Figure S8A**, for competition with MR ligand mannan), highlighting the intrinsic heterogeneity of these cells. Quantification of this effect using phagocytic index (PI)^50^ revealed a distinct glycoform-specific profile with divalent trisaccharide-modified SLP **5** being most effectively taken up in contrast to its hexavalent counterpart **6**. Uptake of the non-glycosylated control **7** was below the limit of detection, emphasizing the glycan-mediated engagement of endocytic lectins.

The cross-presentation efficiency of these glycosylated SLPs was tested next by pulsing differentiated DCs, which were matured for 2 hours with TLR4 ligand MPLA, with 40 nM SLP (which is in a similar concentration range as Glyco-PAINT binding studies, see **Figure S8C, D** for different antigen dosages and quantification) for 2h followed by a 3-day coculture with cognate CD8 T cells that were freshly isolated from OT-I mice. To our surprise, non-mannosylated control **7** was most efficiently cross-presented whereas SLPs decorated with mannosides of increasing complexity and valency **1-6** showed a surprising decrease in T-cell activation/proliferation induction. Being aware of the controversy around the precise role of the MR in antigen cross-presentation^33–36^, we verified next to antigen dose and DC origin (see **Figure S8F** for steady-state cross-presentation of antigens in splenic DCs^34^), the effect of the timing of DC maturation and found an identical pattern for all timepoints in agreement with the observed diminishing effect of antigen mannosylation on T cell proliferation (**Figure S8E** and **G**).

Having observed this non-linear (**Figure S9**) and, to us at least, counter-intuitive relationship between binding, the amounts of endocytosed antigen, and MHC-I-restricted cross-presentation, we next determined whether any of the subcellular kinetic parameters of SLP binding to DCs was predictive of either of these biological functions. A correlation study was performed using the median values of the following 10 kinetic parameters as derived by Glyco-PAINT-APP per analysed square: **τ** (ms), rel. k_on_ (events·sec^-1^µm^-2^), total track duration (s), long track duration (10% longest, s), short track duration (90% shortest, s), diffusion coefficient (nm^2^sec^-1^), speed (µm·s^-1^) and max. speed (µm·s^-1^). In **Table S2** Pearson’s R for the correlation between all these parameters and the phagocytic index or cross-presentation efficiency of SLP **1-7** is listed. Notably, glycan-SLP residence time (**τ**) on the DC surface was the best predictor for cross-presentation, whereas high diffusion coefficients (and accordingly larger displacement of the track) correlated with low uptake of the SLP (**Figure 4E, H**). These observations imply a mechanism where binding events of long duration and high on-membrane mobility are less likely to result in receptor-mediated endocytosis but more likely to enter a cytosolic cross-presentation pathway. Taken together, the strong correlations between on-cell kinetics and cellular functionality demonstrate the potential of the Glyco-PAINT-APP to uncover functional glycan-structure-activity relationships.

### CLR dynamics in polarizing macrophages

To further explore the utility of Glyco-PAINT-APP we studied the changes in carbohydrate binding upon macrophage polarization. Certain pro- and anti-inflammatory stimuli are known to affect absolute levels of the MR on the cell surface upon polarisation of the macrophage from its ‘naïve’ M0-like state to either the inflamed M1-like or wound-healing M2-like phenotype.^51^ It is, however, not known whether this change in MR (and related CLR)-levels also alters binding preferences of the macrophage. We attempted to investigate this by quantifying the binding parameters of MR ligand hexatrimannoside **6** to murine bone marrow-derived macrophages (BMDM) treated with either M1 inducing (LPS and IFNy) or M2-inducing (IL-4) stimuli (**Figure 5A-C**). Quantification of the number of binding events (rel. [R]*k_on_) indicated that the number of **6**-binding receptors was markedly increased upon M2-polarization (**Figure 5F**). When looking at the total binding as quantified by the total track duration (number of events multiplied with their duration) within a square, there appeared to be a vast increase upon M2 stimulation and slight decrease for the M1 phenotype (**Figure 5G**). This suggests the emergence of a new population of long duration binding events that are potentially the result of CLR upregulation by M2 macrophages. Additionally, binding events in the M2 state exhibited longer diffusion coefficients compared to the M1 or naïve states, indicating the presence of a distinct subpopulation (**Figure 5D and H**). Taken together, these kinetic data suggest a mechanism where CLR shuttling is enhanced in inflammatory M1 macrophages, while cell surface dwell times are prolonged in the M2-like state.

**Figure 5.**
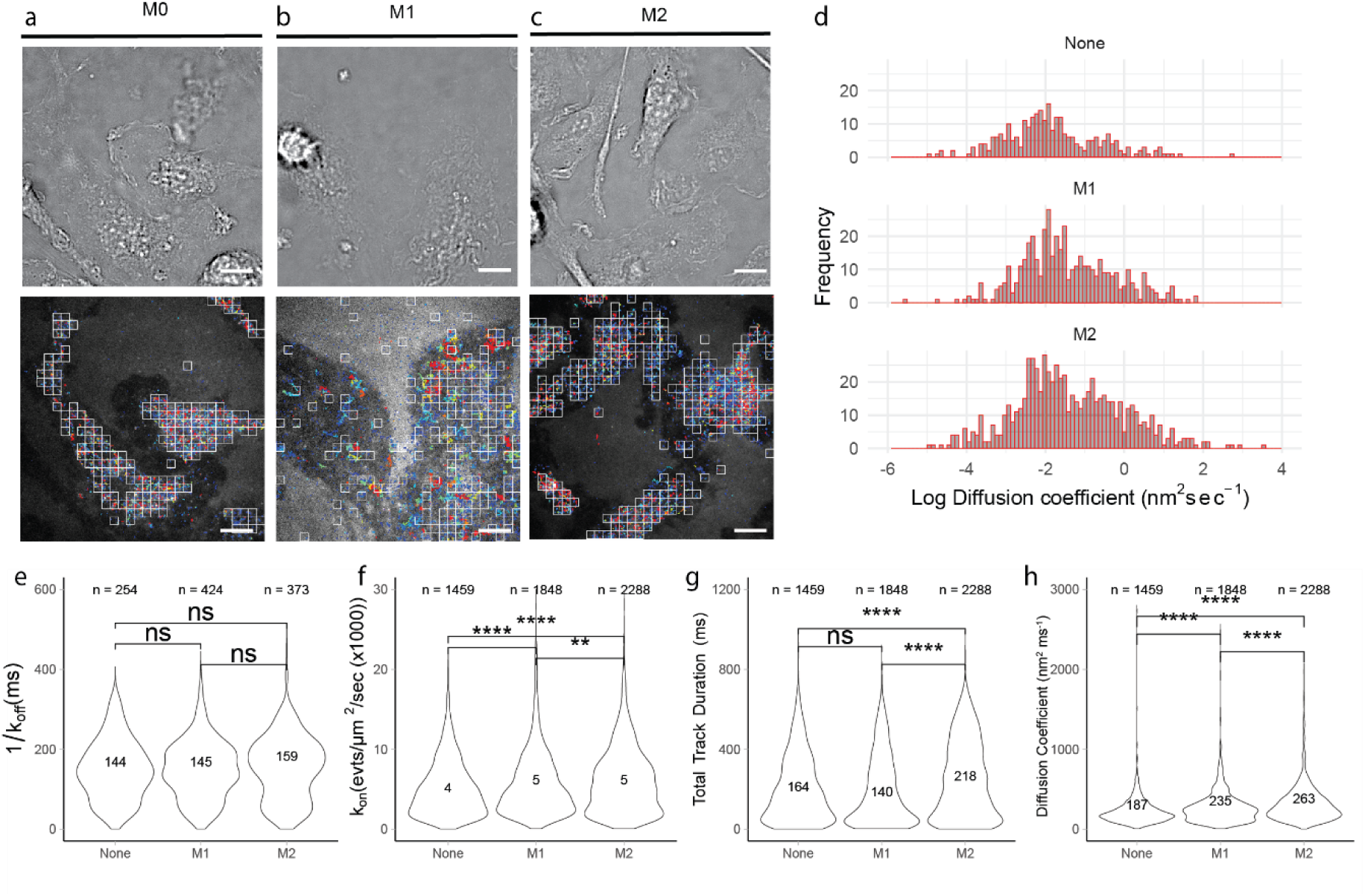
Macrophage polarization alters glycan binding profile. **A-C)** Top: brightfield image live BMDM. Bottom: reconstruction of binding trajectories of glycan **13** to BMDM treated for 16h with no stimulus, an M1-inducing stimulus, or an M2-inducing stimulus. Recordings were processed using a grid sized 20×20 and squares with *Density_Ratio* > 10 and *Nr_Tracks* > 20 are displayed using white borders in the image. **D)** Distribution of diffusion coefficients obtained from the single square with highest event density per recording for the M0-M2 macrophage phenotypes showing the appearance of longer and faster moving tracks upon M2-polarisation. **E-H)** Violin distribution and median value of k_off_^-1^, k_on_, total track durations and diffusion coefficients for glycan **13** binding to BMDM treated with different stimuli showing the appearance of longer binding tracks upon M2-polarization without a change in off rate. n = the number of squares from which the data were obtained. For **E** squares were additionally filtered using *R_Squared > 0.8.* Significance was assessed using two-way ANOVA followed by a Tukey post-hoc test. Tracks are in colored from short to long track length (light blue to red).

## Discussion

This paper describes the single-molecule quantification of glycan-lectin binding on primary immune cells. Whilst quantification of glycan-lectin binding using glycan arrays^52^, SPR^53^ or ELISA-based^54^ technologies with immobilised ligands has yielded a wealth of information related to binding preferences of individual lectins to glycans and vice versa, it has never done so in the context of the living cell surface, where the glycocalyx, other lectins and the constant movement of the membrane can affect glycan binding. Glyco-PAINT-APP offers a significant technological advance that enables the study of glycan-lectin binding in its native context. It is particularly potent, as it now allows the quantification of binding of non-homogeneous distributions of binding events on the cell surface. The method starts with PAINT recordings of glycan binding to heterogeneous, semi-adherent, cell types and is able to extract kinetic information in the form of off-rates, relative on-rates, diffusion coefficients, displacement and speed of the receptor-ligand interaction in an unbiased and high-throughput manner whilst respecting the subcellular variations typically associated with live cells of primary origin. We believe that the Glyco-PAINT-APP can be of use to expand the information that can be obtained from all PAINT-like technologies where increasingly attention is shifting to the use of physiological ligands such as peptides, glycans and proteins as imaging probes beyond the original DNA-PAINT method.^55–57^ Furthermore, our approach could aid in fundamental investigations into the dynamics of immune cell lectins of other families such as the Siglecs. It could also be of use for structure-function or target-engagement studies of other ligands that have a typical low affinity interaction with their receptors such as low-affinity antibodies, peptides or TCR-pMHC interactions^58,59^, or the other interactions where (weak) binding affinities do not appear to correlate with function, such as the recently reported anti-CD40 antibody library, where affinity reduction led to increased receptor engagement.^60^

The second advance is that in this work, the live-cell kinetic parameters that were obtained with the Glyco-PAINT-APP-approach could be correlated to receptor functions, such as uptake and cross-presentation, which had been the subject of much controversy. The wealth of binding and receptor-ligand information obtained by this approach – which had previously thwarted correlation of bulk binding parameters to function - led us to identify key parameters of the interaction that could be predictive of uptake and cross-presentation behaviour: the presence of regions with very long ligand dwell times correlated with cross-presentation ability, and the lack of ligand movement of the receptor with internalisation. This – combined with the fact that uptake efficiency and cross presentation showed no correlation – suggests that multiple processes are likely taking place on the cell surface, either mediated by other lectins, or perhaps simply the micropinocytosis of which dendritic cells display a phenomenal rate: they can internalise their entire cell membrane in about half an hour, or the equivalent of a cell volume in an hour.^61,62^ Watts and co-workers have shown macropinocytosis to be a key mechanism for inducing antigen cross-presentation.^63,64^ We propose the hypothesis that the increased dwell time on the surface of a dendritic cell – either through receptor mediated interaction or through a-specific interactions – increases the chance of an antigen being internalised in a macropinosome. At the same time, the binding of a ligand to a clathrin-anchored MR increases its chance of internalisation into a non-cross presentation enabled vesicle. To precisely delineate the cell biology of these mechanisms, further studies using for example genetic knockouts of DC internalizing receptors are needed. This model, would however, explain some of the other reports of non-linear uptake-cross-presentation relationships for mannosylated SLPs^30–32^ and potentially even for antibody-antigen conjugates.^65^

## Supporting information

supplemnetary manual

## Supplementary Figures

**Figure S1.**
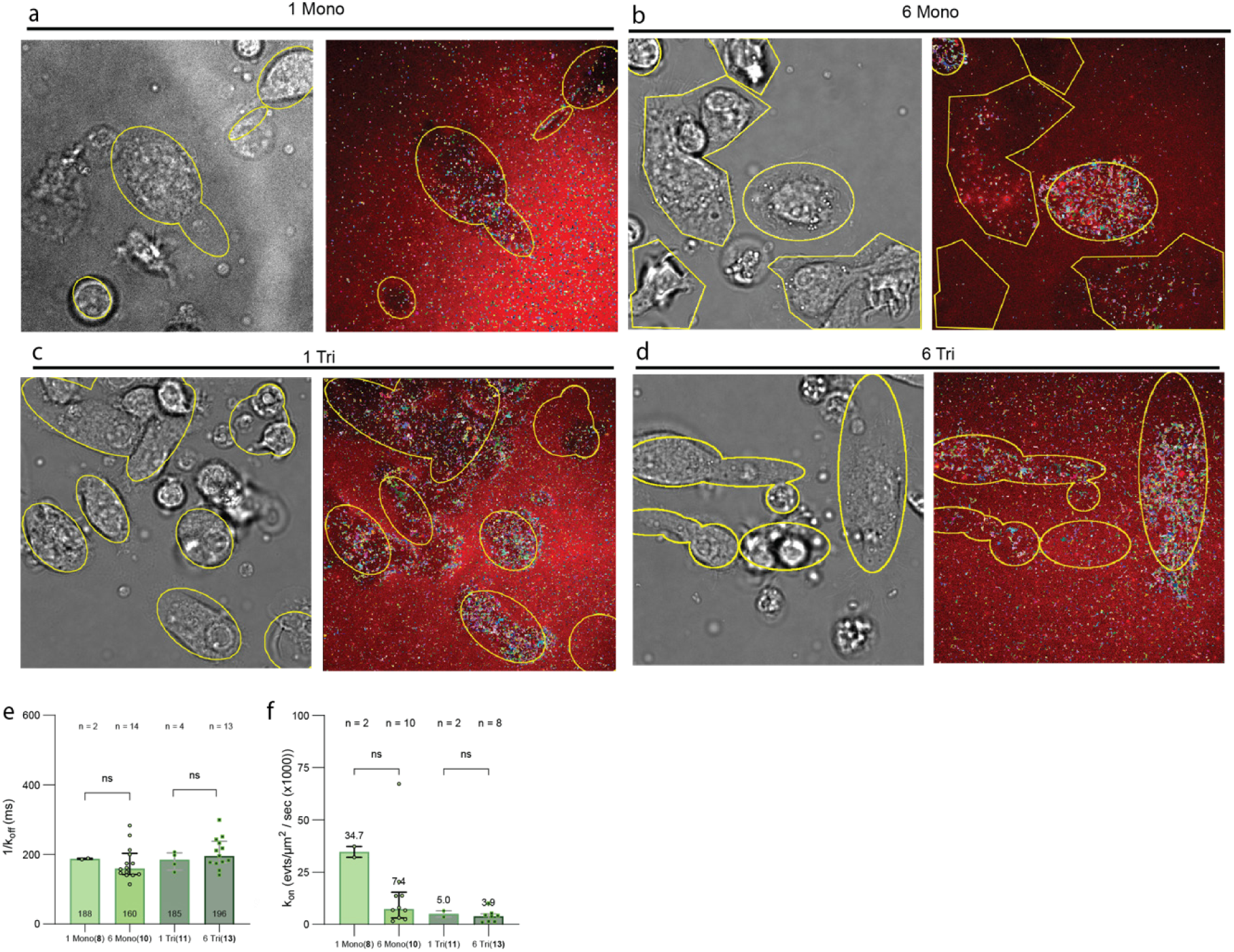
Manual analysis of glycan binding on dendritic cells is complicated by intrinsic cell heterogeneity and results in averaging artefacts. **A)** BMDC were incubated with 5 nM of fluorescent mannoside **8** in a live cell imaging chamber and a brightfield image was captured followed by a 2000 frame Glyco-PAINT recording. Closely linked spots were reconstructed into tracks representing binding events using ImageJ TrackMate plugin and background filtering was performed by manual cell definition using ROI drawing (yellow areas). **B-D)** identical analysis for glycans **10**, **11** and **13,** respectively. **E)** Quantification of the number of events per area which is a relative measure of k_on_. **F)** The average k_off_^-1^ or τ for these events determined by fitting a one-phase exponential decay function over the binding event duration distribution histograms. n, depicts the number of FOVs that were analyzed in total and were acquired over at least 3 biological replicates. Significance was assessed using two-way ANOVA followed by a Tukey post-hoc test. Tracks are colored randomly.

**Figure S2.**
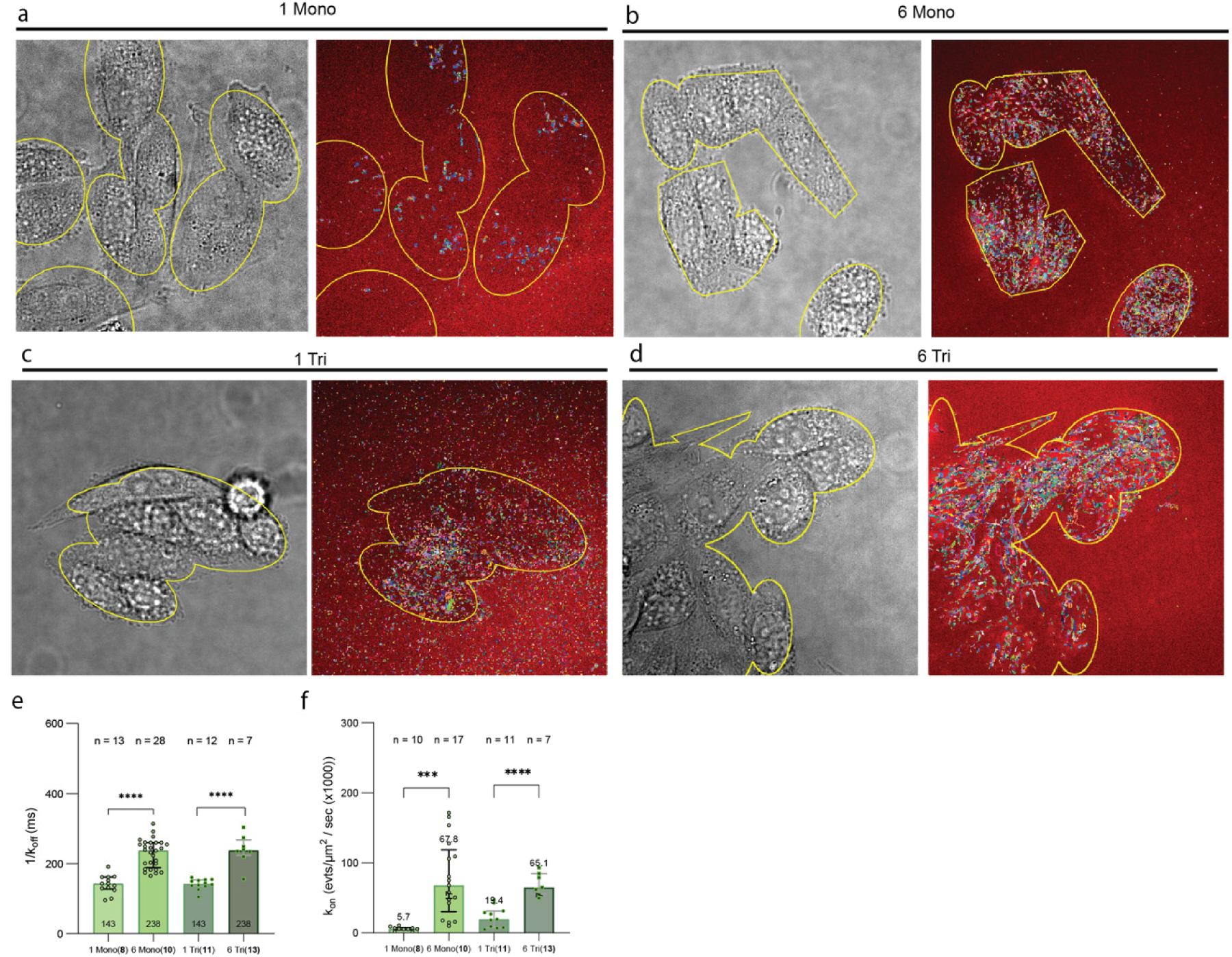
Manual analysis of glycan binding to CHO-MR using original Glyco-PAINT method. **A)** CHO-MR cells were incubated with 1 nM of fluorescent mannoside **8** in a live cell imaging chamber and a brightfield image was captured followed by a 2000 frame Glyco-PAINT recording. Closely linked spots were reconstructed into tracks representing binding events using ImageJ TrackMate plugin and background filtering was performed by manual cell definition using ROI drawing (yellow areas). **B-D)** identical analysis for glycans **10**, **11** and **13,** respectively. **E)** Displays the number of events per area which is a relative measure of k_on_. **F)** Displays the average k_off_^-1^ for these events determined by fitting a one-phase exponential decay function over the binding event duration distribution histograms. n, depicts the number of FOVs that were analyzed in total and were acquired over at least 3 biological replicates. Significance was assessed using two-way ANOVA followed by a Tukey post-hoc test. Tracks are colored randomly.

**Figure S3.**
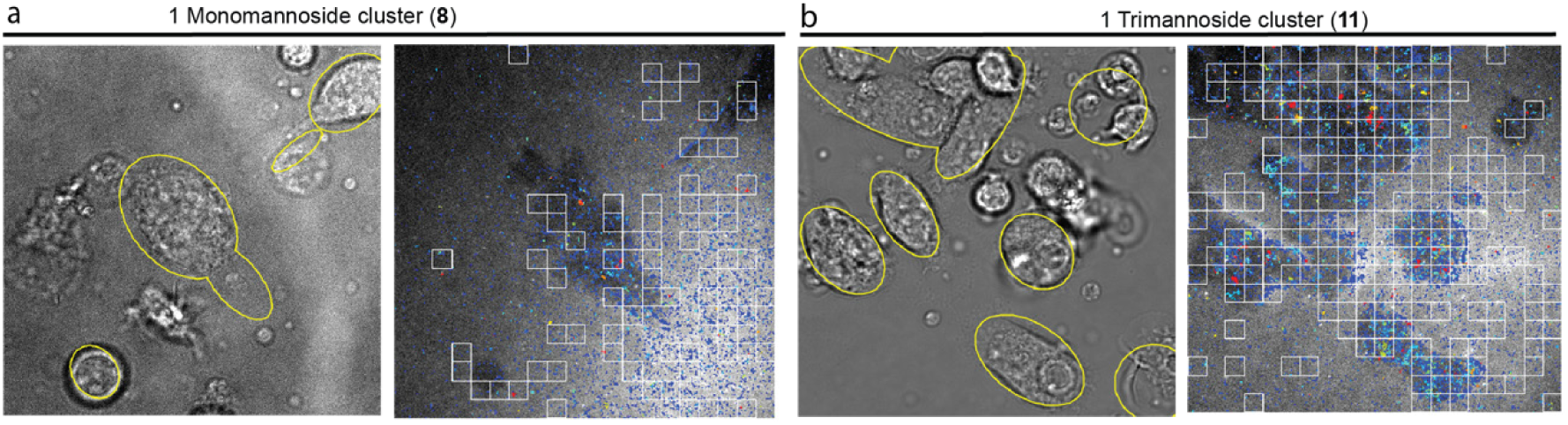
Glyco-PAINT-APP analysis of glycan binding to dendritic cells. **A-B)** Additional to main Figure 2. Brightfield and square-based detection of binding events for representative recordings of glycans **8** and **11**. Analysis and selection parameters are identical as in Figure 2.

**Figure S4.**
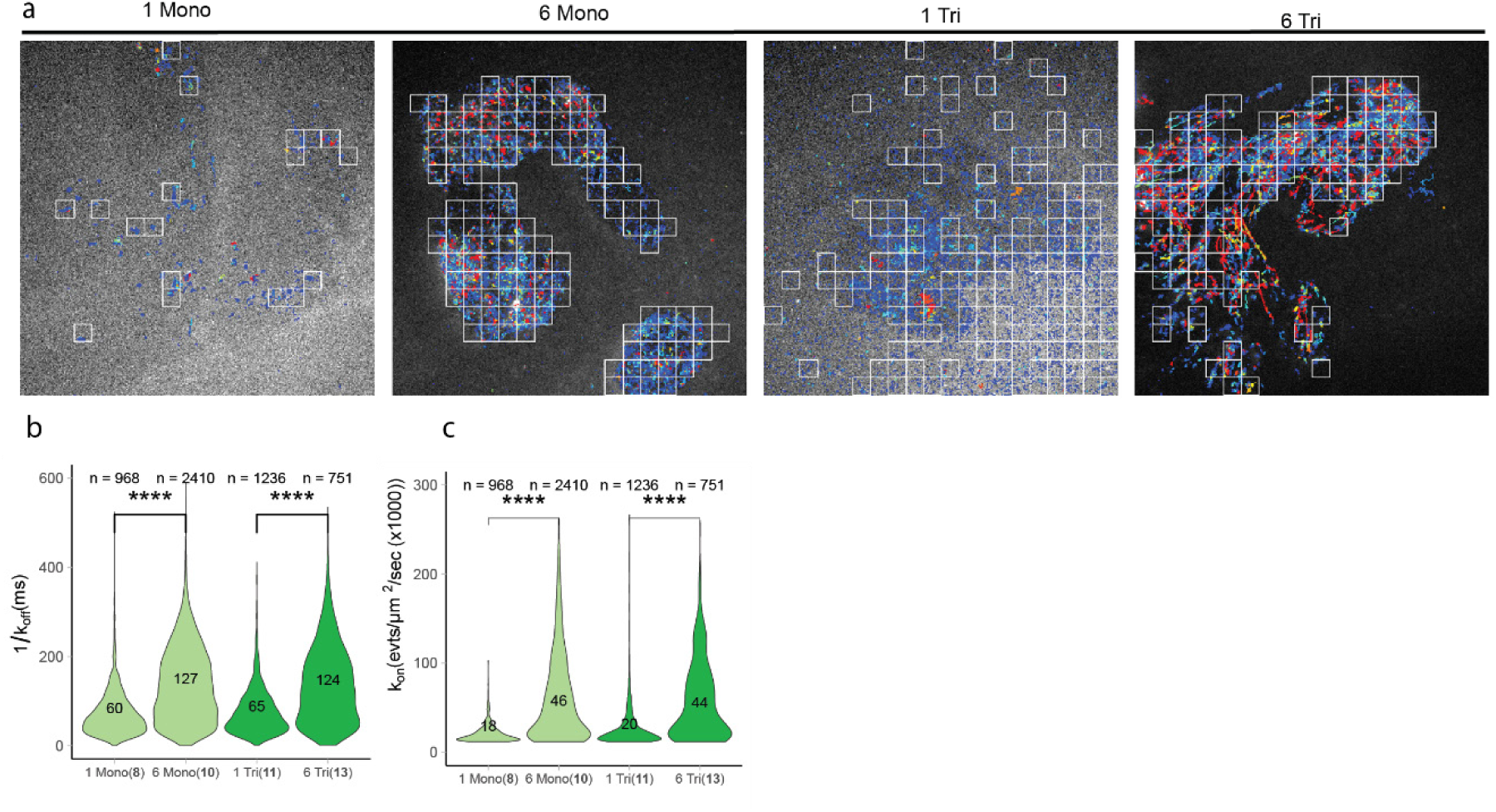
Glyco-PAINT-APP analysis of glycan binding to CHO-MR. **A)** In addition to **Figure S2.** Square-based detection of binding events for representative recordings of glycans **8** and **11. A)** Square-based detection of **8, 10, 11** and **13** binding to CHO-MR, identical images as in shown in **Figure S2** using ROI analysis. **B)** The number of events per area which is a relative measure of k_on_. **C)** average k_off_^-1^ or τ for these events determined by fitting a one-phase exponential decay function over the binding event duration distribution histograms. n, depicts the number of FOVs that were analyzed in total and were acquired over at least 3 biological replicates. Recordings were analyzed using a 20×20 grid and filtered with *Density_Ratio* > 2, *R_Squared* > 0.9 and *Nr_Tracks* > 20. Significance was assessed using two-way ANOVA followed by a Tukey post-hoc test.

**Figure S5.**
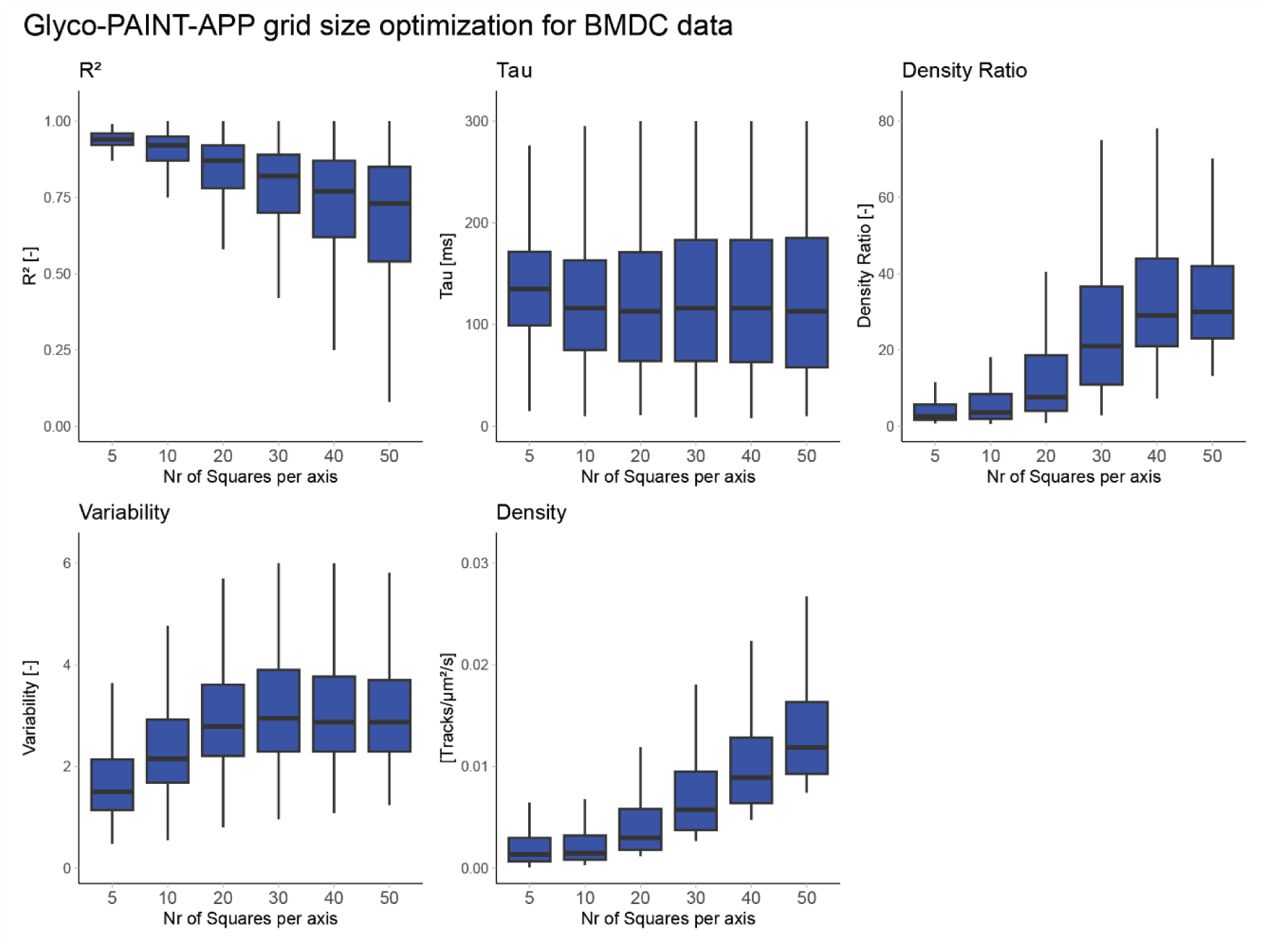
Glyco-PAINT-APP grid size optimization for dendritic cells. Analysis of the effect of mesh size on kinetic and statistical parameters for glycan binding to DC.

**Figure S6.**
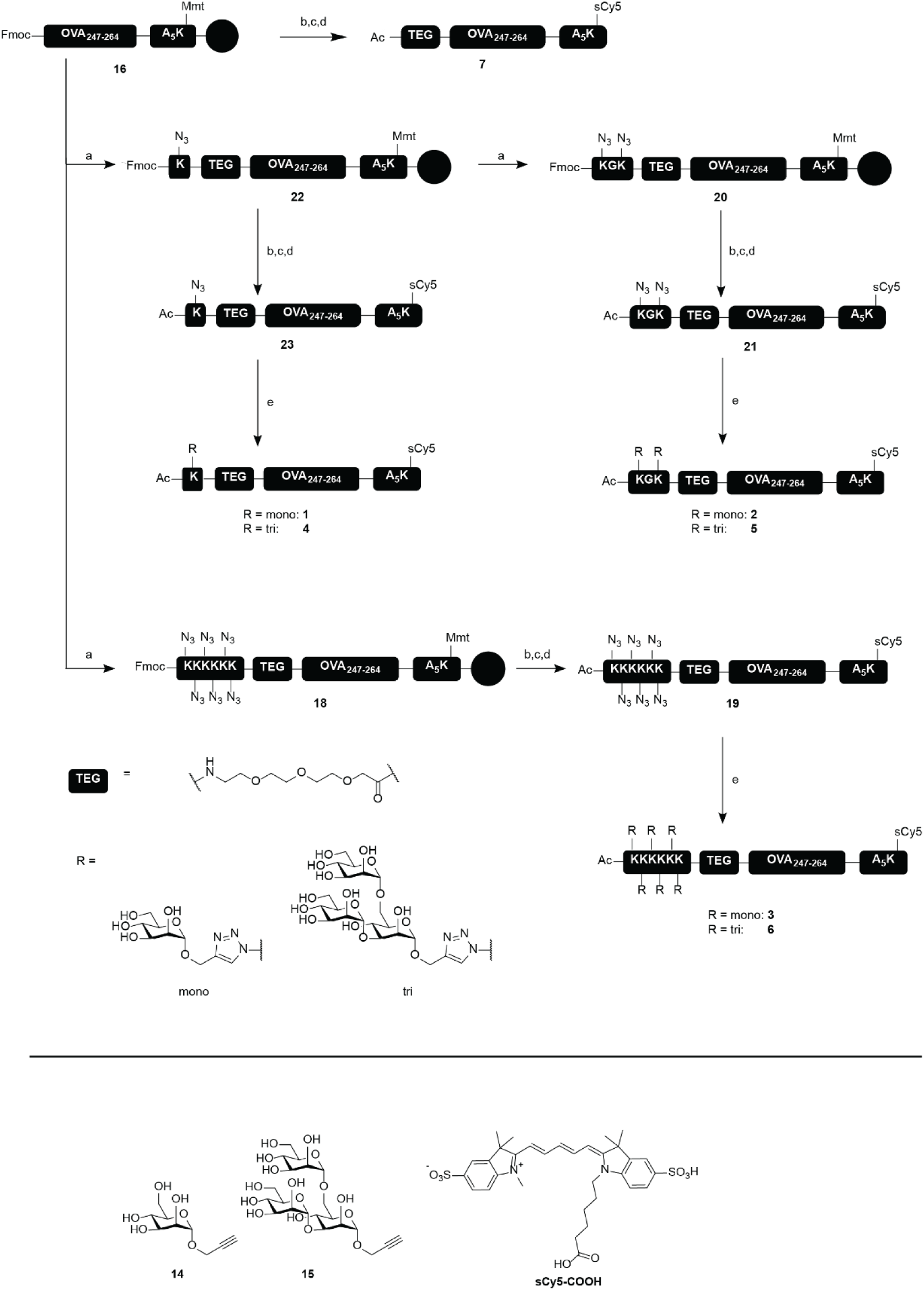
Synthetic scheme for glycopeptides 1-7. Reagents and conditions: a) SPPS (HCTU, Fmoc-TEG-OH, Fmoc-Lys(N_3_)-OH) b) i) 20 % (v/v) piperidine/DMF ii) Ac_2_O, DiPEA, DMF c) i) AcOH, TFE, DCM ii) Et_3_N, DMF iii) **sCy5-COOH**, HCTU, DiPEA, DMF d) i) TFA, TIS, H_2_O ii) RP-HPLC e) i) **14** or **15**, CuSO_4_, NaAsc, THPTA, DMSO, 40°C ii) SEC. Synthetic procedures have been described for propargyl mannosides **14** and **15** by Riera et al.^1^ and Hogervorst and Li et al.^2^ respectively.

**Figure S7.**
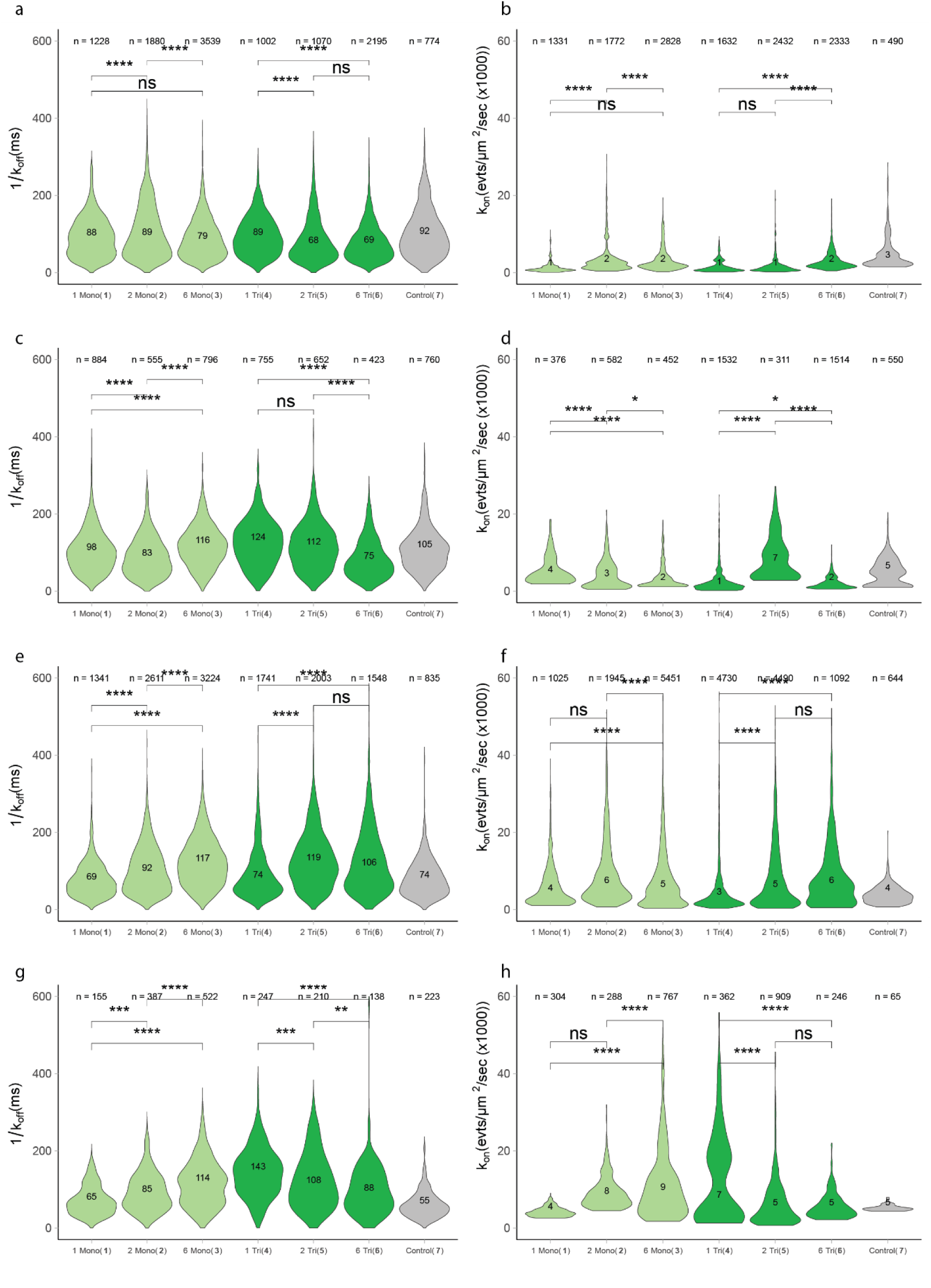
Violin distribution of Glyco-PAINT-APP derived k_on_ and k_off_^-1^ for probe 1-7 binding to BMDC and CHO-MR. Square selection criteria as in main Figure 4**. A,B** BMDC, **C,D** BMDC treated with cytochalasin D, **E,F** CHO-MR and **G,H** CHO-MR treated with cytochalasin D. n, indicates the number of squares that were selected. Significance was assessed using two-way ANOVA followed by a Tukey post-hoc test.

**Figure S8.**
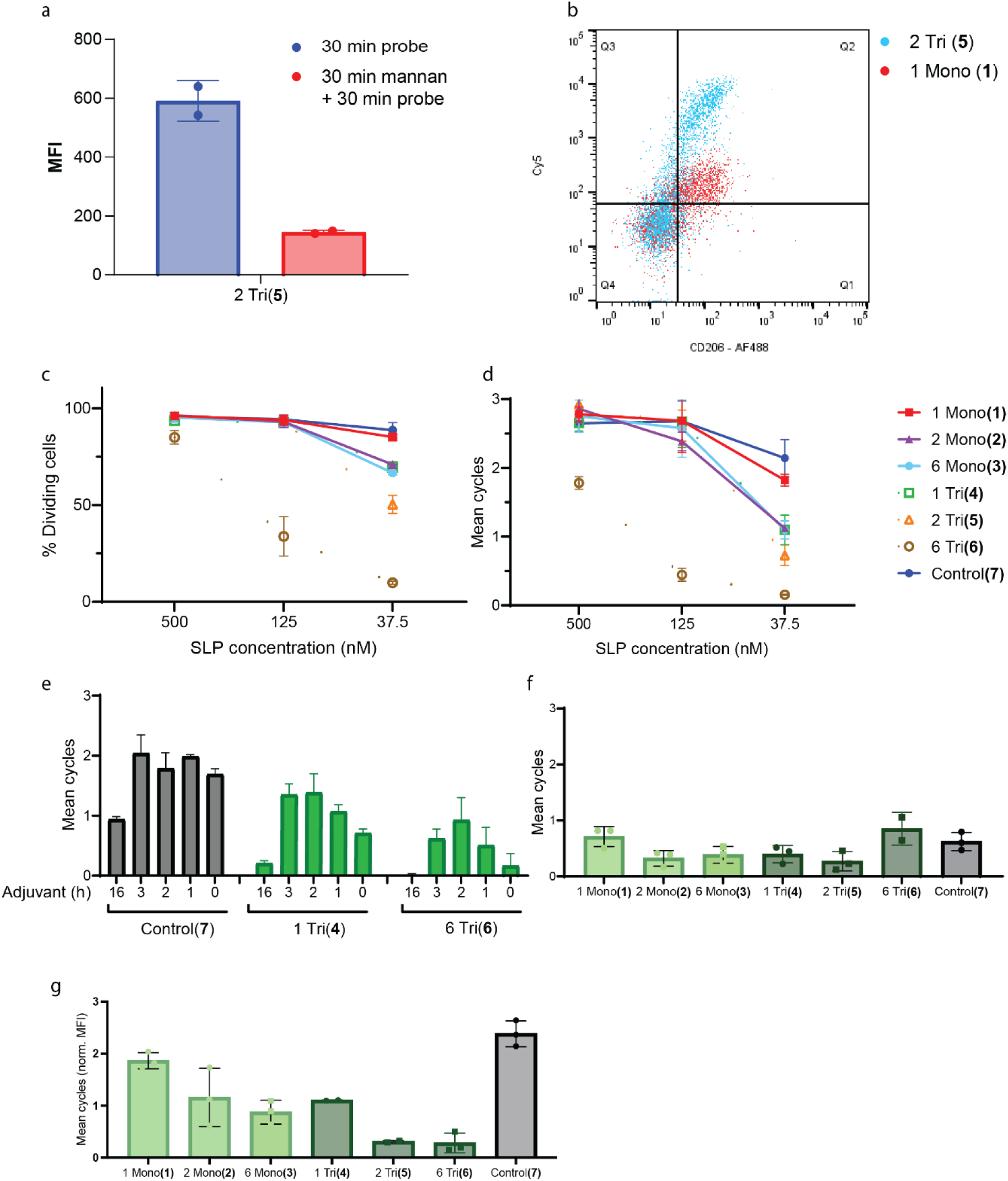
Uptake and cross-presentation of 1-7. **A)** Endocytosis of **4** is blocked by pre-incubation of BMDC with 2 mg/mL mannan for 30 min. **B)** Mannosylated antigens **1** and **4** are taken up by CD11c^+^MR^+^ BMDC. **C-D)** Cross-presentation by BMDC pulsed with different concentrations of antigen **1-7**. Amounts of cross-presentation were determined by flow cytometric measurement of CFSE-dilution by proliferating cognate T cells and quantified using mean cycle or percentage dividing cells of DMSO. **E)** Pretreatment of BMDC with 1 µg/mL MPLA for indicated time before 2h pulse with antigen **1-7** and 3d coculture with OT-I cells. **F**, Cross-presentation of antigens **1-7** by magnetically purified splenic DC. **G**, Cross-presentation of immature BMDC pulsed with SLPs **1-7**

**Figure S9.**
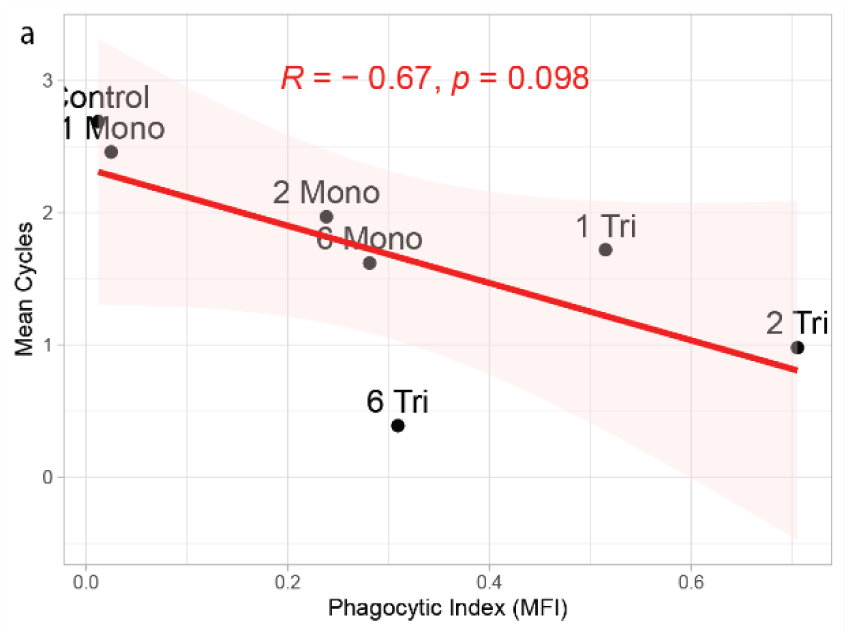
Correlation between phagocytic index and cross-presentation of SLP 1-7 by BMDC. Pearson’s R and two-tailed t test P value are displayed in the correlation plots.

**Table S1.** Median values of Glyco-PAINT-APP derived kinetics for probe 1-7 binding to BMDC and CHO-MR. Square selection criteria as in **Figure 4A-B**. Cells were treated with 10 µM Cytochalasin D for 30 min in indicated tables. TTD, total track duration LTD, long track duration. STD, short track duration. DC, diffusion coefficient.

| Cell Type | Adjuvant | Probe | Tau | Density | TTD | Avg LTD | Avg STD | DC | Avg Speed | Max Speed |
| --- | --- | --- | --- | --- | --- | --- | --- | --- | --- | --- |
| CHOMR | None | 1 Mono | 68 | 4.2 | 16.2 | 1.52 | 0.1 | 200 | 2.1 | 3.2 |
| CHOMR | None | 2 Mono | 91 | 6.3 | 25.8 | 1.50 | 0.1 | 354 | 2.3 | 4.5 |
| CHOMR | None | 6 Mono | 117 | 10.3 | 46.2 | 1.40 | 0.1 | 371 | 2.1 | 4.5 |
| CHOMR | None | 1 Tri | 79 | 4.9 | 17.9 | 1.52 | 0.1 | 211 | 1.7 | 3.2 |
| CHOMR | None | 2 Tri | 117 | 3.6 | 14.5 | 1.38 | 0.1 | 333 | 1.9 | 3.2 |
| CHOMR | None | 6 Tri | 101 | 6.8 | 29.0 | 1.75 | 0.1 | 354 | 2.3 | 4.5 |
| CHOMR | None | Control | 74 | 7.6 | 26.9 | 1.30 | 0.1 | 159 | 1.6 | 3.2 |
| CHOMR | CytD | 1 Mono | 63 | 3.8 | 9.8 | 0.95 | 0.1 | 98 | 1.6 | 3.2 |
| CHOMR | CytD | 2 Mono | 85 | 5.7 | 15.5 | 1.02 | 0.1 | 241 | 2.2 | 4.5 |
| CHOMR | CytD | 6 Mono | 115 | 10.7 | 32.8 | 1.14 | 0.1 | 525 | 2.6 | 4.5 |
| CHOMR | CytD | 1 Tri | 144 | 15.3 | 52.4 | 1.30 | 0.1 | 448 | 2.6 | 4.5 |
| CHOMR | CytD | 2 Tri | 108 | 7.2 | 31.9 | 1.40 | 0.1 | 384 | 2.3 | 4.5 |
| CHOMR | CytD | 6 Tri | 85 | 5.2 | 12.6 | 0.90 | 0.1 | 296 | 2.4 | 4.5 |
| CHOMR | CytD | Control | 55 | 3.7 | 9.9 | 1.12 | 0.1 | 188 | 2.3 | 3.9 |
| BMDC | None | 1 Mono | 87 | 1.8 | 10.9 | 1.12 | 0.1 | 317 | 2.2 | 4.5 |
| BMDC | None | 2 Mono | 96 | 2.2 | 16.5 | 1.70 | 0.1 | 317 | 2.6 | 4.5 |
| BMDC | None | 6 Mono | 76 | 2.3 | 19.9 | 1.62 | 0.1 | 266 | 2.1 | 4.5 |
| BMDC | None | 1 Tri | 88 | 1.6 | 13.6 | 1.25 | 0.1 | 224 | 1.6 | 3.2 |
| BMDC | None | 2 Tri | 69 | 1.5 | 12.1 | 1.65 | 0.1 | 211 | 1.9 | 3.2 |
| BMDC | None | 6 Tri | 68 | 2.1 | 15.8 | 1.55 | 0.1 | 239 | 2.1 | 4.5 |
| BMDC | None | Control | 90 | 2.0 | 13.7 | 1.15 | 0.1 | 349 | 2.4 | 4.5 |
| BMDC | CytD | 1 Mono | 98 | 2.3 | 11.8 | 1.00 | 0.1 | 336 | 2.1 | 3.3 |
| BMDC | CytD | 2 Mono | 81 | 2.4 | 15.4 | 1.23 | 0.1 | 417 | 2.7 | 4.5 |
| BMDC | CytD | 6 Mono | 116 | 1.6 | 9.6 | 1.00 | 0.1 | 387 | 2.4 | 4.5 |
| BMDC | CytD | 1 Tri | 124 | 2.2 | 16.4 | 1.12 | 0.1 | 454 | 2.3 | 4.5 |
| BMDC | CytD | 2 Tri | 112 | 1.7 | 12.4 | 1.10 | 0.1 | 512 | 2.7 | 4.5 |
| BMDC | CytD | 6 Tri | 74 | 2.0 | 12.8 | 1.23 | 0.1 | 230 | 2.1 | 4.5 |
| BMDC | CytD | Control | 105 | 2.0 | 12.0 | 1.05 | 0.1 | 311 | 2.1 | 3.8 |

**Table S2.** Correlation analysis of uptake and cross-presentation with Glyco-PAINT-App derived kinetic parameters. (Absolute) Pearson’s R and two-tailed t test P values are listed for each combination. Correlations presented in **Figure 4G-H** are highlighted in yellow and blue respectively.

| Variable | Correlation_with_mean_cycle | Correlation_with_PI | Abs_Correlation_with_mean_cycle | Abs_Correlation_with_PI |
| --- | --- | --- | --- | --- |
| Tau_E_BMDC_None | 0.896 | -0.611 | 0.896 | 0.611 |
| AvgDC_E_BMDC_None | 0.82 | -0.946 | 0.82 | 0.946 |
| MaxSpeed_E_BMDC_CytD | -0.699 | 0.731 | 0.699 | 0.731 |
| AvgLongTrack_E_BMDC_None | -0.668 | 0.515 | 0.668 | 0.515 |
| AvgLongTrack_E_BMDC_CytD | -0.538 | 0.3 | 0.538 | 0.3 |
| AvgSpeed_E_BMDC_None | 0.402 | -0.785 | 0.402 | 0.785 |
| KdSPR | 0.399 | -0.459 | 0.399 | 0.459 |
| Density_E_BMDC_CytD | 0.389 | -0.383 | 0.389 | 0.383 |
| MaxSpeed_E_BMDC_None | 0.289 | -0.855 | 0.289 | 0.855 |
| Tau_E_BMDC_CytD | 0.255 | 0.36 | 0.255 | 0.36 |
| TotalTrack_E_BMDC_None | -0.247 | -0.067 | 0.247 | 0.067 |
| AvgSpeed_E_BMDC_CytD | -0.216 | 0.62 | 0.216 | 0.62 |
| AvgShortTrack_E_BMDC_None | 0.213 | 0.343 | 0.213 | 0.343 |
| AvgShortTrack_E_BMDC_CytD | 0.097 | 0.402 | 0.097 | 0.402 |
| Density_E_BMDC_None | 0.089 | -0.627 | 0.089 | 0.627 |
| AvgDC_E_BMDC_CytD | 0.021 | 0.692 | 0.021 | 0.692 |
| TotalTrack_E_BMDC_CytD | -0.02 | 0.298 | 0.02 | 0.298 |

### RESOURCE AVAILABILITY

#### Lead contact

Further information and requests for resources and reagents should be directed to and will be fulfilled by the lead contact, Sander van Kasteren

#### Materials availability

A library of glycosylated synthetic long peptide antigens was synthesized in this study. All are available from the lead contact upon reasonable request until stocks run out.

#### Data and code availability

All newly generated code related to processing of Glyco-PAINT recordings is available on GitHub: github.com/jjabakker/GlycoPaint-Pipeline

Any additional information required to reanalyze the data reported in this paper is available from the lead contact upon request.

### EXPERIMENTAL MODEL DETAILS

#### Mice

Male C57Bl/6J and OT-I (C57BL/6-Tg(TcraTcrb)1100Mjb/J) mice were purchased from the Jackson Laboratory (CA, USA). The animals were bred and housed under specific pathogen-free conditions and provided with water and food ad libitum under a 12:12 day/night cycle. Mice ranging from 8 to 15 weeks old were euthanized by cervical dislocation before harvest of lymphoid organs and/or thigh bones, femur, and tibia. All animal experiments received approval from the Dutch Central Authority for Scientific Procedures on Animals (CCD) on license number AVD1060020198832 and were conducted in accordance with the European Union Directive 2010/63/EU, recommendation 2007/526/EC.

#### Cell culture

##### Bone Marrow-Derived Dendritic Cells (BMDC)

Bone marrow (BM) was isolated from femurs, tibias, and thigh bones of C57Bl/6J mice via centrifugation (1900 *rcf*, 4.5 min) of scissor-cut bones that were placed in a 1.5 mL tube. The resulting pellet was subjected to red blood cell (RBC) lysis by resuspending in 0.5 mL of ammonium chloride-potassium (ACK) lysis buffer (Gibco, A1049201). After 3 min incubation at rt, the suspension was filtered over a 70 µm filter (Falcon, 352350), rinsed with 5 mL PBS and washed once with PBS by 5 min centrifugation at 300 *rcf* at rt. Thus obtained BM was resuspended at 1E6 cells / mL in 15 cm uncoated culture dishes (Sarstedt, 82.1184.500) in complete RPMI-1640 (Capricorn, RPMI-A) supplemented with 10% heat-inactivated fetal calf serum (FCS, Gibco, A5670701), penicillin (100 I.U./mL) & streptomycin (50 µg/mL) (Gibco, 15140148), 2 mM GlutaMAX (Gibco, 35050061), 50 µM 2-mercaptoethanol (Gibco, 31350010) and 20 ng/mL mGM-CSF (Peprotech, 315-03) and cultured in a humidified incubator at 37 °C and 5% CO2. On day two, 5mL of the above-described fresh medium was added and on day four cells were reseeded in fresh medium at 1E6/mL. Cells were used for microscopy and T cell activation experiments on day 7 or 8.

##### Bone Marrow-Derived Macrophages (BMDM)

Bone marrow (BM) was isolated from femurs, tibias, and thigh bones of C57Bl/6J mice via centrifugation (1900 *rcf*, 4.5 min) of scissor-cut bones that were placed in a 1.5 mL tube. The resulting pellet was subjected to red blood cell (RBC) lysis by resuspending in 0.5 mL of ACK lysing buffer (Gibco, A1049201). After 3 min incubation at rt, the suspension was filtered over a 70 µm filter (Falcon, 352350), rinsed with 5 mL PBS and washed once with PBS by 5 min centrifugation at 300 *rcf* at rt. Thus obtained BM was resuspended at 0.8E6 cells/mL in 15 cm uncoated culture dishes (Sarstedt, 82.1184.500) in complete RPMI-1640 (Capricorn, RPMI-A) supplemented with 10% heat-inactivated fetal calf serum (FCS, Gibco, A5670701), penicillin (100 I.U./mL) & streptomycin (50 µg/mL) (Gibco, 15140148), 2 mM GlutaMAX (Gibco, 35050061), 50 µM 2-mercaptoethanol (Gibco, 31350010) and 20 20 ng/mL M-CSF (Biolegend, 576404) and cultured in a humidified incubator at 37°C and 5% CO_2_. On day 2, 5 mL fresh medium was added and on day 4 medium was aspirated and replenished with 15 mL fresh medium. Cells were used for experiments on day 7 or 8.

##### CHO-MR

The CHO-MR cell line was kindly provided by Luisa Martinez-Pomares^3^ and cultured in DMEM/F12 without phenol red (Gibco, 21041025), supplemented with 10% FCS, penicillin (100 I.U./mL), streptomycin (50 µg/mL) and selection antibiotic G418 (0.6 mg/mL). A layer of adherent cells was washed with PBS and cells were harvested by 10 min incubation with 2 mM EDTA in PBS and subcultured approximately twice per week at a 1:5 split when cells reached 70-80% confluency.

### EXPERIMENTAL METHOD DETAILS

#### Statistical analysis and sample size

Statistical analyses were conducted to compare glycan ligand binding kinetics across probes using the Glyco-PAINT square-based subsampling technology. Subsampling subcellular regions (squares) within fields of view increased the number of independent data points, enhancing statistical power compared to treating entire fields of view or cells as single units. For all Glyco-PAINT experiments at least 5 biological replicates (independent experiments with fresh mouse material or new passage number for cell lines) with 3 technical replicates (fields of view per condition) were recorded. For flow cytometry assays at least 3 biological replicates with 2 technical replicates per condition were conducted. A two-way ANOVA was used to assess differences among probes with respect to kinetic parameters derived from Glyco-PAINT experiments and flow cytometric assays, followed by Tukey’s Honest Significant Difference (HSD) test for post-hoc comparisons to control Type I error. Significance is reported as: ns (not significant), *p* ≥ 0.05; *p* < 0.05 (*); *p < 0.01* (**); *p < 0.001* (***); and *p* < 0.0001 (****).

#### Fluorescent glycan probes

Glycan and glycan SLP probes were stored as lyophilized powders at −20°C. Upon thawing, vials were reconstituted in DMSO and concentration was determined by measurement of absorbance using Nanodrop apparatus with extinction coefficients e_sCy5_ = 250.000 M^-1^cm^-1^ at 641 nm and e_ATTO655_ = 125.000 M^-1^cm^-1^ at 663 nm. Small aliquots were stored at −20 °C until use.

#### OT-I T cell isolation

A 70 µm filter was placed on a 50 mL tube and pre-wetted with PBS supplemented with 2 mM EDTA and 2% FCS (single-cell suspension buffer, SCSB). Freshly harvested spleens from OT-I transgenic mice were placed on the filter and disrupted with the back end of a syringe. After thorough washing, the suspension was centrifuged (10 min., 300 *rcf*, rt). Next, the pellet was gently resuspended in 2 mL of ACK lysing buffer (Gibco, A1049201). After 3 min incubation the suspension was diluted with 10 mL PBS, filtered once more, and centrifuged again (300 *rcf*, 10 min, rt). The pellet was resuspended and subjected to magnetically-activated depletion of non-target cells using the “Naïve CD8a+ T cell isolation Kit” (Miltenyi, 130-096-543) according to manufacturer’s protocol. The obtained T cells were counted and resuspended at a density of 5-15E6 / mL in 1 mL PBS with 5 µM CFSE (Biolegend) and incubated for 15 min. at 37 °C. After incubation, cells were spun down (10 min, 300 rcf, rt), washed once more with complete RPMI-1640 and were ready for downstream use.

#### Splenic DC isolation

Spleens isolated from C57Bl/6J mice were placed on a 10 cm petri dish (Sarstedt, 83.3902) containing 5 mL HBSS (Gibco, 14025092) supplemented with 1 mg/mL collagenase IV (Sigma, NC2115693) and 20 U/mL DNAse (Thermo Scientific, EN0525) and minced into small pieces. After incubating for 30 min at 37 °C, tissue digestion was stopped by addition of 2 mM EDTA. The remaining homogenate was disrupted and RBC-lysed as described above and subjected to magnetically-activated depletion of non-target cells using the “Pan Dendritic Cell isolation Kit” (Miltenyi, 130-100-875) according to manufacturer’s protocol.

#### Glycan uptake experiments

2.5E5 BMDC or CHO-MR were seeded in a 96 well v-bottom plate (Sarstedt, 82.1583001). The next day, Glycan SLP probes were added to the cells at 250 nM and incubated for 1h at 37 °C. Active uptake was halted by addition of ice-cold PBS and washed twice (300 *rcf*, 5 min., rt). Cells were stained with Zombie Yellow (Biolegend, 423103, 1:500), TruStain FcX (Biolegend, 101319, 1:100), CD11c - eFluor450 (clone: N418, eBioscience, 48-0114-82, 1:200), CD206-AF488 (clone: MR5D3, Biorad, MCA2235A488T, 1:20), acquired on Guava EasyCyte 12HT and analyzed using FlowJo v8. Phagocytic index (PI) was calculated according to equation 1. Statistical analysis and plotting were performed using GraphPad Prism V10.

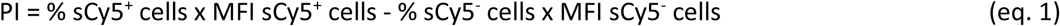

#### T cell proliferation assay

5E4 BMDC or 1E5 splenic DC were seeded in 96 well Nunc U-bottom plates (Thermo Scientific, 168136) and pulsed with Glycan SLP at 40 nM for indicated time. In indicated experiments, DC were pretreated with 1 µg/mL MPLA (Avanti, 699800P) for 2h before antigen pulse. After antigen pulse was finished, the plate was spun down for 3 min at 600 *rcf*, washed once with complete RPMI-1640 and 1-1.5E5 freshly isolated, CFSE-stained OT-I T cells were added to the pulsed DCs in 200 µL complete RPMI-1640 and incubated at 37 °C. After 3d of coculture, the plate was spun down (3 min, 600 rcf, 4 °C) and supernatant was removed and stored at −80°C. Cells were washed with FACS buffer (PBS with 2 mM EDTA, 2% FCS and 7.4 mM NaN3) and stained with Aqua Live/DEAD (Invitrogen, L34957, 1:500), TruStain FcX (Biolegend, 101319, 1:100), CD8a - APC (clone: 53-7.6, Biolegend, 100711, 1:200), and TCR V beta 5.1/5.2 - eFluor450 (clone: MR9-4, eBioscience, 48-5796-82) for 30 min on ice, washed two times and acquired on a BD Fortessa I flow cytometer and analyzed using FlowJo v8. To calculate mean cycle, the CFSE dilution factor was obtained by dividing the MFI of the antigen pulsed condition by the DMSO pulsed condition expressed as Log2.^4^ Statistical analysis and plotting was performed using GraphPad Prism V10.

#### BMDM polarization

At day 6 adherent macrophages were harvested by aspiration of medium, washing once with PBS and incubation with 10 mL of 2 mM EDTA in PBS for 10 min at 37°C. Cells were reseeded in complete medium with addition of polarizing stimulus, 20 ng/mL IFN-y (Peprotech, 315-05-100UG) + 100 ng/mL LPS-EB (Invivogen, tlrl-eblps) for M1 or 20 ng/mL IL-4 (Peprotech, 214-14-20UG) for M2 for 16h.

#### Optical setup

Single molecule imaging was performed on a Nikon Ti2 N-STORM system equipped with TIRF module, Z piezo element, perfect focus system for axial drift correction and an OkoLab incubator with temperature and CO2 controller (37 °C and 5% CO2) for live-cell imaging. Recordings were acquired using the 647 nm excitation laser (160 mW, 1.9 kW/cm2). Upon laser excitation, fluorescence was collected by a 100x 1.49 NA oil-immersion objective, passed through a quad-band dichroic mirror (97335 Nikon), and detected by a Hamamatsu ORCA Flash 4.0 CMOS camera with 160 nm pixel size. The signal was collected using the following settings: 512×512 pixel region, no binning, pixel depth 16-bit, exposure time 50 ms, live-cell observation and 2D-STORM (lens out), zoom 1x, lens x0.4 and for live-cell observation at 37°C the correction collar was set to position 8160.

#### Acquisition of Glyco-PAINT recordings

5E4 CHO-MR or 1E5 BMDC were seeded in 8 well glass-bottomed microscopy slides (Ibidi, 80827) in complete medium. After equilibration in the microscope incubator, fluorescent glycan was added at 5 nM for CHO-MR and 10 nM for BMDC experiments. Then, cells were brought into focus using brightfield illumination and 2,000 frames (at 50 ms intervals) were recorded within a single field-of-view at 40-60% of maximum 647 nm laser power using TIRF illumination. For indicated experiments, Cytochalasin D (Focus Biomolecules) was added to a final concentration of 10 µM for 30 min prior to acquisition.

#### ROI-based analysis of Glyco-PAINT recordings

TrackMate was run manually using the Fiji plugin.^5^ The LoG spot detection algorithm was applied with an object radius of 0.5 µm, with pre-processing with median filter and sub-pixel localization unchecked. Threshold values were set to 5, 10 or 15 such that no more than 800.000 spots were detected and kept identical per experimental condition. Next, single-particle tracking was performed using the Simple LAP tracker algorithm with a maximum frame gap of 3, a max linking distance of 0.6 μm and a gap closing max distance of 1.2 μm. Tracks with only two spots were discarded. Then, manual Regions of Interest (ROIs) were drawn in individual recordings using Fiji based on brightfield-defined cell outlines. The number of tracks residing within the ROI were compared with the number of tracks outside the ROI to establish a ratio of cell / glass density. Only recordings for which that ratio exceeded 2 were considered for further analysis. For all remaining recordings, the k_on_, k_off_ and MSD were calculated for tracks within the ROI according to the procedure as described by Riera et al.^1^ Statistical analysis and plotting was performed using GraphPad Prism v10.

#### Glyco-PAINT-APP analysis of recordings

Processing of recordings using Glyco-PAINT-APP was performed as described in a step-by-step procedure in the **Supplementary Manual** and accompanying **Supplementary Videos 1 and 2**. For the TrackMate processing, a batch file (Experiment Info.csv) containing the experiment metadata and tracking parameters was created. Threshold values were set to 5, 10 or 15 such that no more than 800.000 spots were detected and kept identical per experimental condition. Recordings were then processed in TrackMate using the ‘Run TrackMate Batch’ plugin provided by the Glyco-PAINT-APP. Through this plugin, spot detection and tracking by TrackMate^5^ was performed as indicated in the batch file using the Simple LAP tracker algorithm with a maximum frame gap of 3, a max linking distance of 0.6 μm and a gap closing max distance of 1.2 μm. Tracks with only two spots were discarded.

With the Glyco-PAINT-APP utility ‘Generate Squares’, a grid of squares was overlaid and kinetic properties for each square were calculated. Default parameters for grid processing are (deviations are mentioned in figure captions): Nr of Squares in row 20, Minimum Tracks to Calculate Tau 20, Min allowable R Squared 0.1, Min Required Density Ratio 2 and Maximum Allowable Variability 10.

For every recording, a background track count was calculated by averaging the track count of the 40 (10% of the total number of squares) least dense squares. Only squares for which the track count exceeded the Min Required Density Ratio of 2 were considered (and the remaining ignored as background). For each square also the variability was calculated and only squares for which the variability was less than the Maximum Allowable Variability of 10 were considered. For squares meeting both the Minimum Required Density Ratio and Maximum Allowable Variability criteria, and containing at least the Minimum Tracks to Calculate Tau kinetic parameters including k_on_, k_off_ and MSD were calculated as in Riera et al.^1^, or copied from the TrackMate Tracks table output (for velocity, displacement, and track duration).

Summary files were created using the ‘Compile Project’ utility, creating an ‘All Recordings.csv’ file, containing information for all 973 recordings, an ‘All Squares.csv’ file containing information on 389,200 (973 x 400) squares and an ‘All Tracks.csv’ file containing information on 13.4 mln tracks (binding events) in the project. Statistical analysis and plotting using these merged files was performed using the ggplot2 package in R.^6,7^

#### General procedure for automated SPPS

Peptides were synthesized using automated Fmoc-SPPS on a Liberty Blue^tm^ automated microwave peptide synthesizer (CEM corporation). Synthesis was performed 100 μmol scale on Tentagel S RAM resin (loading 0.20-0.25 mmol/g, Rapp Polymere GmbH, Germany). Resin was first swollen for 5 minutes in DMF prior to amino acid coupling. Activation was achieved using DIC/Oxyma coupling as recommended by the manufacturer. The following amino acids were used: Fmoc-Ala-OH, Fmoc-Arg(Pbf)-OH, Fmoc-Asn(Trt)-OH, Fmoc-Asp(OtBu)-OH, Fmoc-Gln(tBu)-OH, Fmoc-Glu(OtBu), Fmoc-Gly-OH, Fmoc-Ile-OH, Fmoc-Leu-OH, Fmoc-Lys(Boc)-OH, Fmoc-Lys(Mmt)-OH, Fmoc-Phe-OH, Fmoc-Ser(tBu)-OH, Fmoc-Val-OH. All amino acids were obtained from Novabiochem except Fmoc-Lys(Mmt)-OH which was obtained from CEM corporation. Standard coupling was achieved using 5 equivalents amino acid as a 0.2 M amino acid/DMF solution, 5 equivalents DIC as a 0.5 M of DIC/DMF solution and 5 equivalents Oxyma as a 1 M Oxyma/DMF solution (also containing 0.2 M DiPEA), at 90°C for 2 minutes. Standard Fmoc deprotection was achieved by 20% (v/v) piperidine in DMF at 90°C for 90 seconds, repeated once. To analyze the quality of the peptide, a small amount of resin (∼1mg) was treated with 200 µL of a TFA cocktail (95:2.5:2.5, TFA/H_2_O/TIS) for 2 hours, after which the TFA was filtered into 800 µL of ice cold Et_2_O. After five minutes the formed precipitate was collected by centrifugation and the supernatant discarded. The pellet was dissolved in 200 µL 1:1:1 H_2_O/MeCN/tBuOH and subjected to LC-MS analysis. Peptides were characterized using electrospray ionization mass spectrometry (ESI-MS) on a Thermo Finnigan LCQ Advantage Max LC-MS instrument with a Surveyor PDA plus UV detector on an analytical C18 column (Phenomenex, 3 μm, 110 Å, 50 mm × 4.6 mm) in combination with buffers A (H_2_O), B (MeCN), and C (1% aq TFA). Quality of crude was evaluated with a linear gradient of 10-90% B with a constant 10% C over 10 minutes.

#### General procedures for manual SPPS

Manual elongation of peptides was carried out in a fritted syringe at either 25 or 5 µmol scale. Fmoc deprotection was achieved using 20 % (v/v) piperidine in DMF in two steps, reacting 3 and 7 minutes, respectively. Fmoc-Gly-OH was coupled using 5 equivalents of amino acid together with 5 equivalents of HCTU (as a 0.5 M solution) and 10 equivalents DiPEA for 45 minutes. Fmoc-Lys(N_3_)-OH was coupled using 2 equivalents together with 2 equivalents HCTU (as a 0.2 M in DMF solution) and 4 equivalents of DiPEA for 90 minutes. Fmoc-TEG-OH was coupled using 4 equivalents, together with 4 equivalents HCTU and 8 equivalents DiPEA in 1 mL of DMF for 45 minutes. Analysis of the quality of the resin-bound peptide was carried out as above.

#### General procedure for *N*-terminal acetylation

The *N*-terminal Fmoc was cleaved by treating the resin twice with a 20% (v/v) solution of piperidine in DMF, for 3 and 7 minutes. This was followed by the addition of a solution containing 10 % (v/v) Ac_2_O and 5 % (v/v) DiPEA in DMF (1 mL for 25 µmol resin bound peptide). This resin was acetylated for 15 minutes under gentle agitation, followed by draining the acetylation solution and thorough washing of the resin with DMF.

#### General procedure for fluorophore labeling

Fluorophore labeling of peptides was carried out on a 5 µmol scale. The resin was first swelled in DCM, followed by selective Mmt deprotection. The Mmt group protecting the C-terminal lysine residue, the resin was treated with a mildly acidic mixture consisting of 10 % (v/v) AcOH and 20 % (v/v) TFE in DCM (1 mL) for one hour^8^. After this time has elapsed, the resin was washed three times with DCM and three times with DMF. To remove residual acetic acid, the resin was then treated with a 10 % TEA in DMF mixture (2 x 10 min) followed by an additional three washes with DMF. One equivalent of fluorophore was dissolved in a 50 mM solution of HCTU together with 2 equivalents of DiPEA. This mixture was added to the drained resin and allowed to react under gentle agitation overnight protected from light. After overnight coupling the solution was drained and the resin was thoroughly washed with DMF.

#### General procedure for peptide global deprotection and purification

Peptide cleavage was carried out using a standard TFA cleavage cocktail (TFA:H_2_O:TIS 95:2.5:2.5) using 1 mL per 25 µmol of resin bound peptide. Before initiating cleavage, the resin was thoroughly washed with DCM and drained. The TFA cocktail was added and mixed with the resin under gentle agitation for one hour, after which it was drained into a centrifuge tube containing ice cold diethyl ether (10:1 ratio Et_2_O:TFA). When deprotecting fluorophore modified peptides, the cleavage reaction was protected from light as much as possible and a 20:1 ratio of Et_2_O to TFA was used. The diethylether/TFA mixture was chilled for a minimum of 10 minutes to increase peptide recovery, and the precipitated peptide was recovered by centrifugation. The supernatant was discarded and the pellet washed with a small amount of Et_2_O, followed again by centrifugation. This pellet was dissolved in a mixture of H_2_O:MeCN:tBuOH and subjected to RP-HPLC purification on a Gilson GX281 semipreparative HPLC. This machine was equipped with a Gemini-NX C18 column (5 µm, 110 A, 250 x 10.0 mm) using a flow of 5 mL/min and buffers A = 0.1% TFA in H_2_O and B = MeCN. Peak detection was done using a UV-Vis detector set to 225 nm or 610 nm for fluorescently labeled peptides. Quality of purified peptides was determined using an electrospray ionization mass spectrometry (ESI-MS) on a Thermo Finnigan LCQ Advantage Max LC-MS instrument with a Surveyor PDA plus UV detector on an analytical C18 column (Phenomenex, 3 μm, 110 Å, 50 mm × 4.6 mm) in combination with buffers A (H_2_O), B (MeCN), and C (1% aq TFA). Quality of the peptides was evaluated with a linear gradient of 10-50% B with a constant 10% C over 9 minutes or a linear gradient of 10-90% B with a constant 10% C over 9 minutes.

**Fmoc-Lys(N_3_)-Lys(N_3_)-Lys(N_3_)-Lys(N_3_)-Lys(N_3_)-Lys(N_3_)-TEG-Asp(OtBu)-Glu(OtBu)-Val-Ser(tBu)-Gly-Leu-Glu(OtBu)-Gln(Trt)-Leu-Glu(OtBu)-Ser(tBu)-Ile-Ile-Asn(Trt)-Phe-Glu(OtBu)-Lys(Boc)-Leu-Ala-Ala-Ala-Ala-Ala-Lys(Mmt)-RAM-Tentagel S (18)**

Resin bound peptide **16** (25 µmol) was elongated manually with Fmoc-TEG-OH and Fmoc-Lys(N_3_)-OH. **LC-MS** RT = 7.6 min (C18, 10-90% B over 9 minutes) **LRMS** calcd [M+2H]^2+^ = 1941.52, [M+3H]^3+^ = 1294.68 observed M/z = 1941.53, 1294.80

**Ac-Lys(N_3_)-Lys(N_3_)-Lys(N_3_)-Lys(N_3_)-Lys(N_3_)-Lys(N_3_)-TEG-Asp-Glu-Val-Ser-Gly-Leu-Glu-Gln-Leu-Glu-Ser-Ile-Ile-Asn-Phe-Glu-Lys-Leu-Ala-Ala-Ala-Ala-Ala-Lys(sCy5)-NH_2_ (19)**

Resin bound peptide **18** (5 µmol) was N-terminally acetylated according to the standard conditions. This was followed by chemoselective deprotection of the Mmt group and coupling of sCy5-OH (1.0 eq, 3.2 mg, 5 µmol) as described in the general methods. Global deprotection followed by RP-HPLC purification yielded compound **19** as a blue solid (1.60 mg, 0.37 µmol, 7.4%). **LC-MS** RT = 6.0 min (C18, 10-90% B over 9 minutes) **LRMS** calcd [M+3H]^3+^ = 1443.06 observed M/z = 1443.00

**Fmoc-Lys(N_3_)-Gly-Lys(N_3_)-TEG-Asp(OtBu)-Glu(OtBu)-Val-Ser(tBu)-Gly-Leu-Glu(OtBu)-Gln(Trt)-Leu-Glu(OtBu)-Ser(tBu)-Ile-Ile-Asn(Trt)-Phe-Glu(OtBu)-Lys(Boc)-Leu-Ala-Ala-Ala-Ala-Ala-Lys(Mmt)-RAM-Tentagel S (20)**

Resin bound peptide **22** (25 µmol) was elongated manually with Fmoc-TEG-OH, Fmoc-Lys(N_3_)-OH and Fmoc-Gly-OH. **LC-MS** RT = 6.5 min (C18, 10-90% B over 9 minutes) **LRMS** calcd [M+2H]^2+^ = 1661.86, [M+3H]^3+^ = 1108.24 observed M/z = 1662.00, 1108.67

**Ac-Lys(N_3_)-Gly-Lys(N_3_)-TEG-Asp-Glu-Val-Ser-Gly-Leu-Glu-Gln-Leu-Glu-Ser-Ile-Ile-Asn-Phe-Glu-Lys-Leu-Ala-Ala-Ala-Ala-Ala-Lys(sCy5)-NH_2_ (21)**

Resin bound peptide **20** (5 µmol) was N-terminally acetylated according to the standard conditions. This was followed by chemoselective deprotection of the Mmt group and coupling of sCy5-OH (1.0 eq, 3.2 mg, 5 µmol) as described in the general methods. Global deprotection followed by RP-HPLC purification yielded compound **21** as a blue solid (1.24 mg, 0.33 µmol, 6.6%). **LC-MS** RT = 5.0 min (C18, 10-90% B over 9 minutes) **LRMS** calcd [M+2H]^2+^ = 1883.93, [M+3H]^3+^ = 1256.28 observed M/z = 1884.42, 1256.67

**Ac-Lys(N_3_)-TEG-Asp(OtBu)-Glu(OtBu)-Val-Ser(tBu)-Gly-Leu-Glu(OtBu)-Gln(Trt)-Leu-Glu(OtBu)-Ser(tBu)-Ile-Ile-Asn(Trt)-Phe-Glu(OtBu)-Lys(Boc)-Leu-Ala-Ala-Ala-Ala-Ala-Lys(Mmt)-RAM-Tentagel S (22)**

Resin bound peptide **16** (25 µmol) was elongated manually with Fmoc-TEG-OH, Fmoc-Lys(N_3_)-OH and Fmoc-Gly-OH, followed by acetylation under standard conditions. **LC-MS** RT = 4.9 min (C18, 10-90% B over 9 minutes) **LRMS** calcd [M+2H]^2+^ = 1465.77, [M+3H]^3+^ = 977.51 observed M/z = 1466.33, 977.83

**Ac-Lys(N_3_)-TEG-Asp-Glu-Val-Ser-Gly-Leu-Glu-Gln-Leu-Glu-Ser-Ile-Ile-Asn-Phe-Glu-Lys-Leu-Ala-Ala-Ala-Ala-Ala-Lys(sCy5)-NH_2_ (23)**

Resin bound peptide **22** (5 µmol) was chemoselectively deprotected to remove the Mmt group, followed by coupling of sCy5-OH (1.0 eq, 3.2 mg, 5 µmol) as described in the general methods. Global deprotection followed by RP-HPLC purification yielded compound **23** as a blue solid (0.88 mg, 0.25 µmol, 4.9%). **LC-MS** RT = 4.7 min (C18, 10-90% B over 9 minutes) **LRMS** calcd [M+2H]^2+^ = 1777.87, [M+3H]^3+^ = 1185.58 observed M/z = 1778.67, 1186.08

**Ac-TEG-Asp-Glu-Val-Ser-Gly-Leu-Glu-Gln-Leu-Glu-Ser-Ile-Ile-Asn-Phe-Glu-Lys-Leu-Ala-Ala-Ala-Ala-Ala-Lys(sCy5)-NH_2_ (7)**

Resin bound peptide **16** (25 µmol) was elongated manually with Fmoc-TEG-OH followed by N-terminal acetylation according to the standard conditions. Next, chemoselective deprotection of the Mmt group and coupling of sCy5-OH (1.0 eq, 3.2 mg, 5 µmol) as described in the general methods yielded the fully protected, resin bound peptide. Global deprotection followed by RP-HPLC purification yielded compound **7** as a blue solid (1.44 mg, 0.42 µmol, 8.5%). **LC-MS** RT = 4.6 min (C18, 10-90% B over 9 minutes) **LRMS** calcd [M+2H]^2+^ = 1701.32, [M+3H]^3+^ = 1134.54 observed M/z = 1701.58, 1134.75

#### CuAAC glycopeptide conjugation

##### General procedures for CuAAC modification of peptides

Peptides were dissolved in degassed DMSO and glycans in either degassed DMSO or degassed MilliQ water. These were mixed in a 1.5 mL Eppendorf tube, followed by addition of a copper click mix. This click mix typically consistent of 1 part 0.1 M CuSO_4_ (in MQ), 2 parts 0.1 M sodium ascorbate (in MQ) and 3 parts 0.1 M THPTA (in DMSO), giving a solution with a total Cu(I) concentration of around 15 mM. Of this solution, enough was added to be 0.15 equivalents compared to the peptide. The reaction was heated in a 40°C shaker block, and every 24 hours 1 equivalent of 0.1 M sodium ascorbate was added. Periodically, a 0.5 µL sample was diluted into 39.5 µL of 1:1:1 H_2_O:MeCN:tBuOH and analyzed by LC-MS to evaluate reaction progression. After full conversion of the peptide towards the desired conjugate was observed, the reaction mixture was subjected to size exclusion chromatography over Toyopearl HW-40 size exclusion resin using 150 mM NH_4_OAc or 150 mM NH_4_HCO_3_ (containing 20% MeCN) as the buffer. Fractions showing absorbance at 610 nm were combined and lyophilized.

**Ac-Lys(Man)-Lys(Man)-Lys(Man)-Lys(Man)-Lys(Man)-Lys(Man)-TEG-Asp-Glu-Val-Ser-Gly-Leu-Glu-Gln-Leu-Glu-Ser-Ile-Ile-Asn-Phe-Glu-Lys-Leu-Ala-Ala-Ala-Ala-Ala-Lys(sCy5)-NH_2_ (3)**

Compound **19** (100 nmol, 50 mM in DMSO) was mixed with propargyl mannoside **14** (12 eq., 6 µL of 200 mM solution in DMSO). 1 µL of the standard click mix was added and the reaction carried out at 40°C. After SEC purification and lyophilization compound **3** was obtained as a blue powder (0.48 mg, 85 nmol, 85%). **LC-MS** RT = 4.3 min (C18, 10-50% B over 9 minutes) **LRMS** calcd [M+3H]^3+^ = 1879.21, [M+4H]^4+^ = 1409.66 observed M/z = 1879.33, 1409.83

**Ac-Lys(triMan)-Lys(triMan)-Lys(triMan)-Lys(triMan)-Lys(triMan)-Lys(triMan)-TEG-Asp-Glu-Val-Ser-Gly-Leu-Glu-Gln-Leu-Glu-Ser-Ile-Ile-Asn-Phe-Glu-Lys-Leu-Ala-Ala-Ala-Ala-Ala-Lys(sCy5)-NH_2_ (6)**

Compound **19** (100 nmol, 50 mM in DMSO) was mixed with propargyl mannoside **15** (12 eq., 6 µL of 200 mM solution in DMSO). 1 µL of the standard click mix was added and the reaction carried out at 40°C. After SEC purification and lyophilization compound **6** was obtained as a blue powder (0.41 mg, 54 nmol, 54%). **LC-MS** RT = 4.2 min (C18, 10-50% B over 9 minutes) **LRMS** calcd [M+4H]^4+^ = 1896.07, [M+5H]^5+^ = 1517.06 observed M/z = 1896.08, 1517.17

**Ac-Lys(Man)-Gly-Lys(Man)-TEG-Asp-Glu-Val-Ser-Gly-Leu-Glu-Gln-Leu-Glu-Ser-Ile-Ile-Asn-Phe-Glu-Lys-Leu-Ala-Ala-Ala-Ala-Ala-Lys(sCy5)-NH_2_ (2)**

Compound **21** (100 nmol, 25 mM in DMSO) was mixed with propargyl mannoside **14** (4 eq., 2 µL of 200 mM solution in DMSO). 1 µL of the standard click mix was added and the reaction carried out at 40°C. After SEC purification and lyophilization compound **2** was obtained as a blue powder (0.25 mg, 60 nmol, 60%). **LC-MS** RT = 6.8 min (C18, 10-50% B over 9 minutes) **LRMS** calcd [M+3H]^3+^ = 1402.00 observed M/z = 1402.08

**Ac-Lys(triMan)-Gly-Lys(triMan)-TEG-Asp-Glu-Val-Ser-Gly-Leu-Glu-Gln-Leu-Glu-Ser-Ile-Ile-Asn-Phe-Glu-Lys-Leu-Ala-Ala-Ala-Ala-Ala-Lys(sCy5)-NH_2_ (5)**

Compound **21** (100 nmol, 25 mM in DMSO) was mixed with propargyl mannoside **15** (4 eq., 2 µL of 200 mM solution in DMSO). 1 µL of the standard click mix was added and the reaction carried out at 40°C. After SEC purification and lyophilization compound **5** was obtained as a blue powder (0.29 mg, 60 nmol, 60%). **LC-MS** RT = 6.6 min (C18, 10-50% B over 9 minutes) **LRMS** calcd [M+3H]^3+^ = 1618.07 observed M/z = 1618.08

**Ac-Lys(Man)-TEG-Asp-Glu-Val-Ser-Gly-Leu-Glu-Gln-Leu-Glu-Ser-Ile-Ile-Asn-Phe-Glu-Lys-Leu-Ala-Ala-Ala-Ala-Ala-Lys(sCy5)-NH2 (1)**

Compound **23** (100 nmol, 25 mM in DMSO) was mixed with propargyl mannoside **14** (2 eq., 1 µL of 200 mM solution in DMSO). 1 µL of the standard click mix was added and the reaction carried out at 40°C. After SEC purification and lyophilization compound **1** was obtained as a blue powder (0.30 mg, 79 nmol, 79%). **LC-MS** RT = 4.4 min (C18, 10-90% B over 9 minutes) **LRMS** calcd [M+2H]^2+^ = 1886.91, [M+3H]^3+^ = 1258.27 observed M/z = 1887.33, 1258.75

**Ac-Lys(triMan)-TEG-Asp-Glu-Val-Ser-Gly-Leu-Glu-Gln-Leu-Glu-Ser-Ile-Ile-Asn-Phe-Glu-Lys-Leu-Ala-Ala-Ala-Ala-Ala-Lys(sCy5)-NH_2_ (4)**

Compound **23** (100 nmol, 25 mM in DMSO) was mixed with propargyl mannoside **15** (2 eq., 1 µL of 200 mM solution in DMSO). 1 µL of the standard click mix was added and the reaction carried out at 40°C. After SEC purification and lyophilization compound **4** was obtained as a blue powder (0.26 mg, 63 nmol, 63%). **LC-MS** RT = 4.4 min (C18, 10-90% B over 9 minutes) **LRMS** calcd [M+3H]^3+^ = 1366.31 observed M/z = 1366.75

## LC-MS analysis

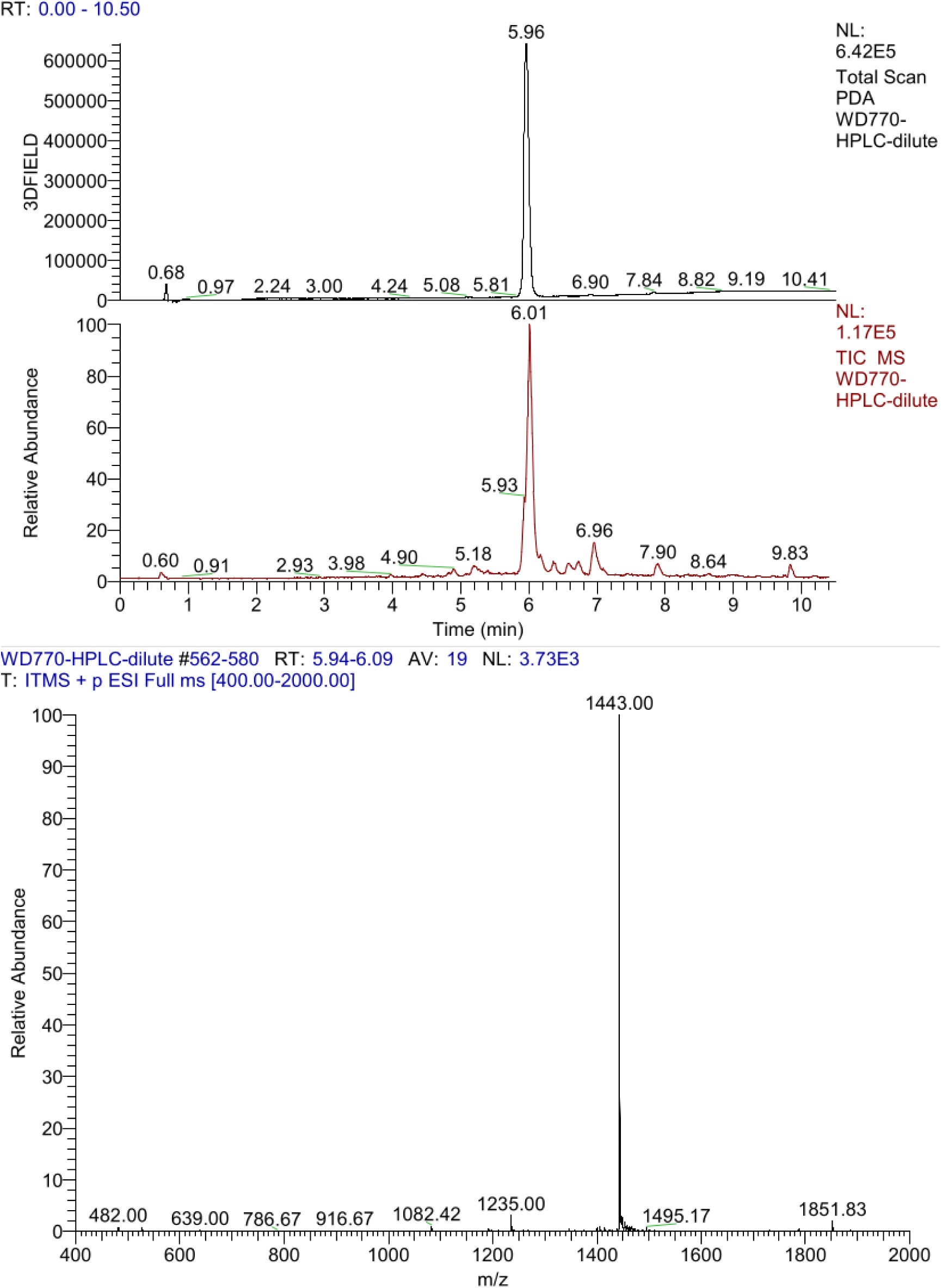

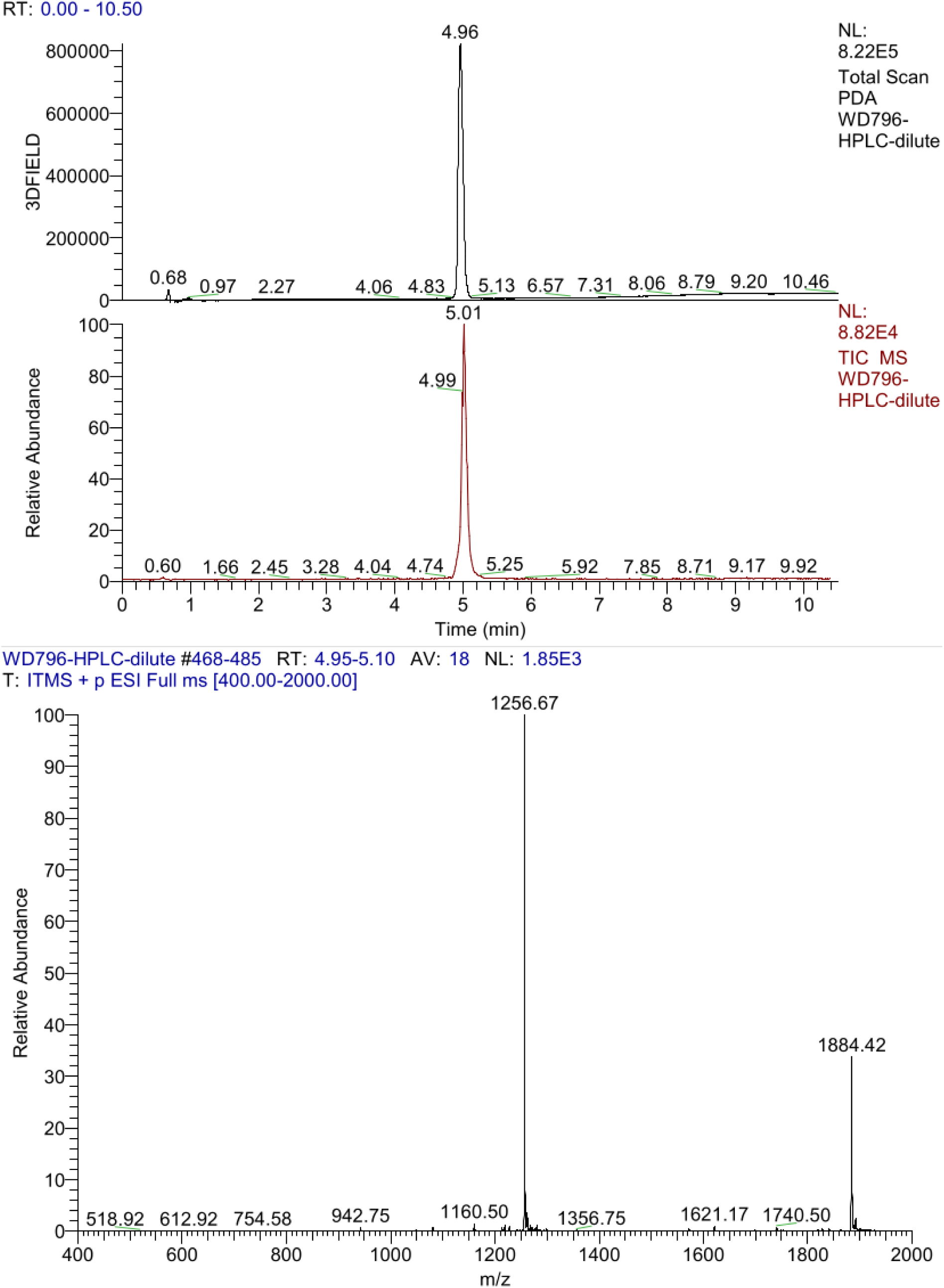

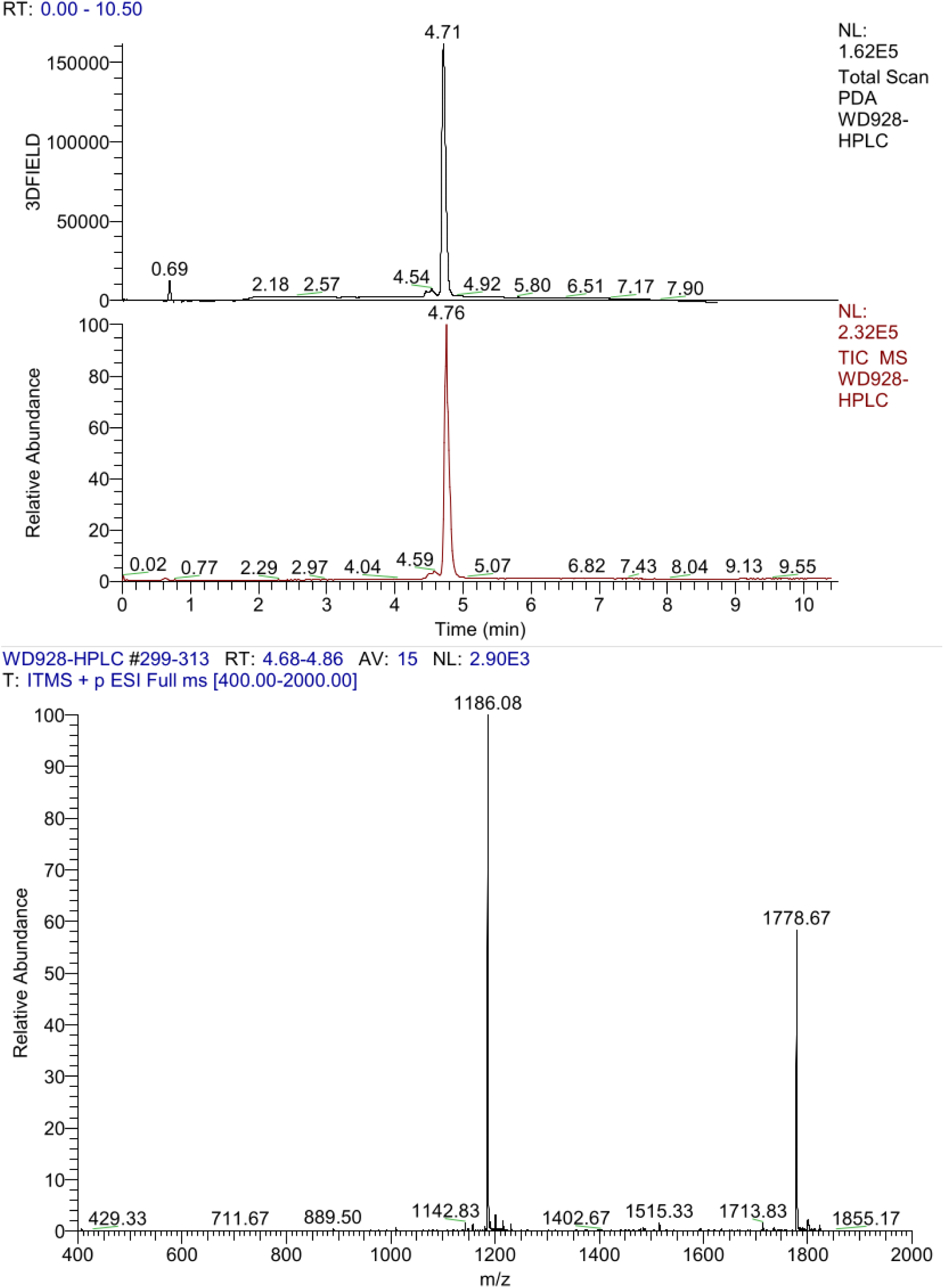

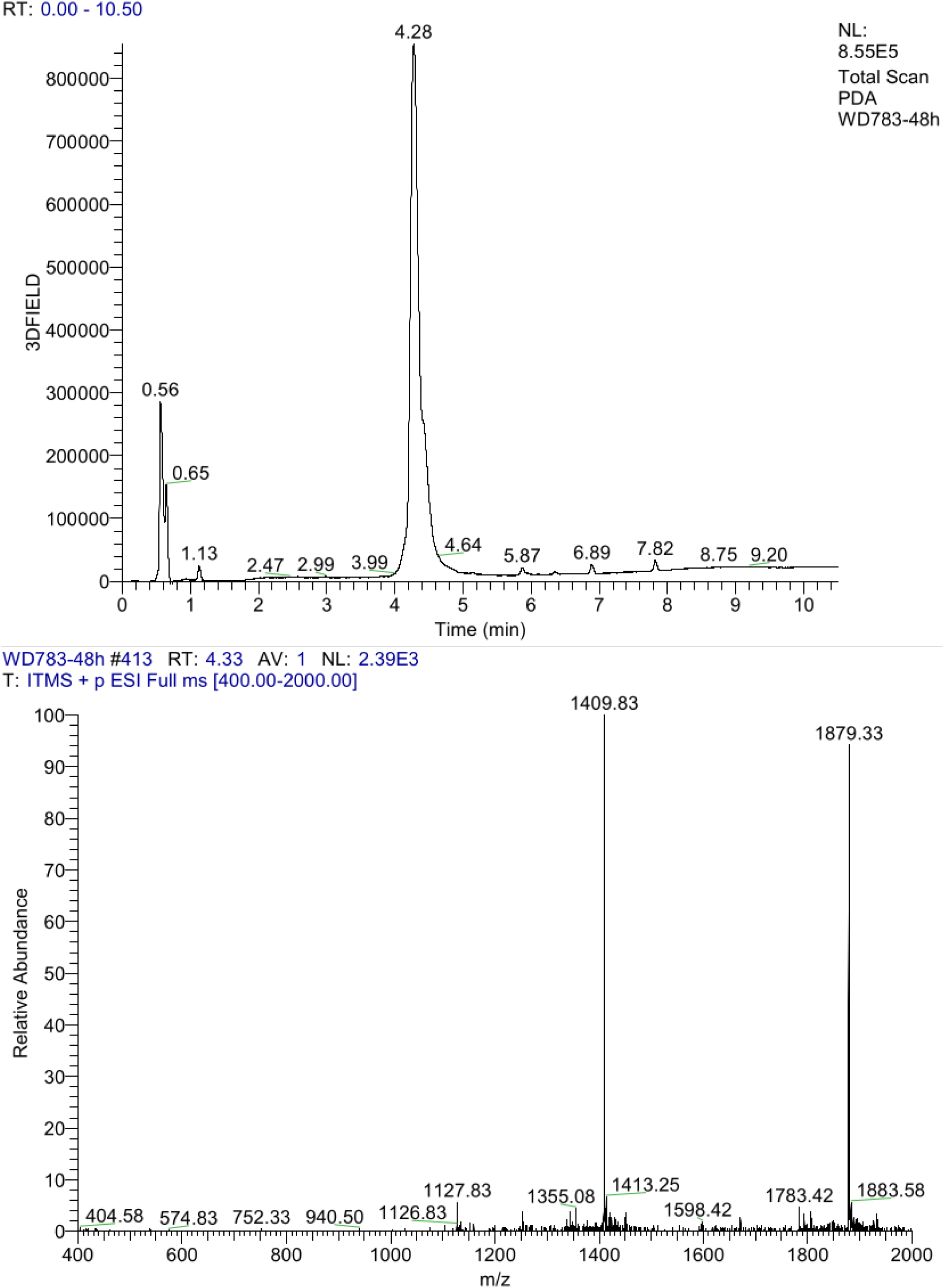

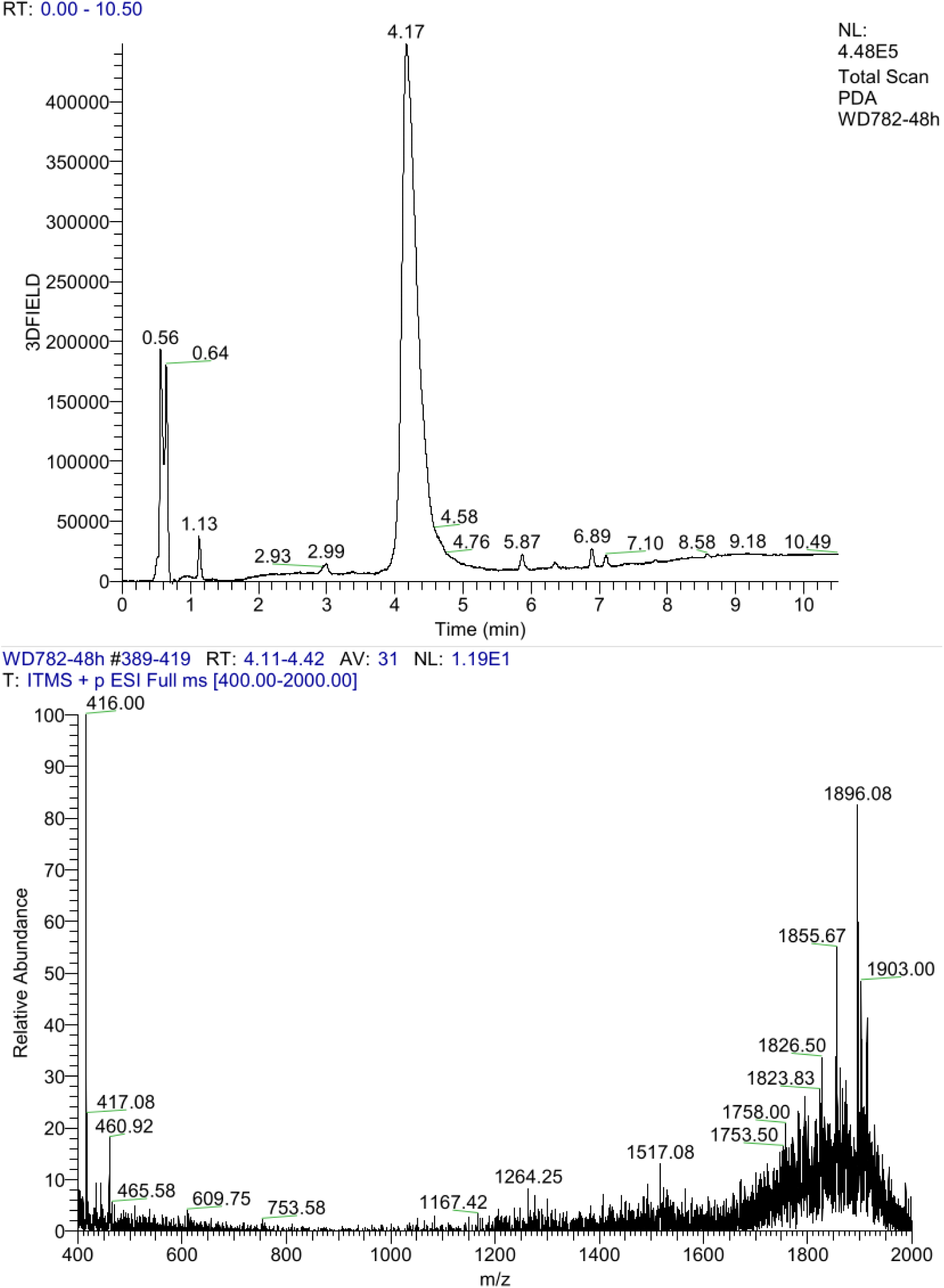

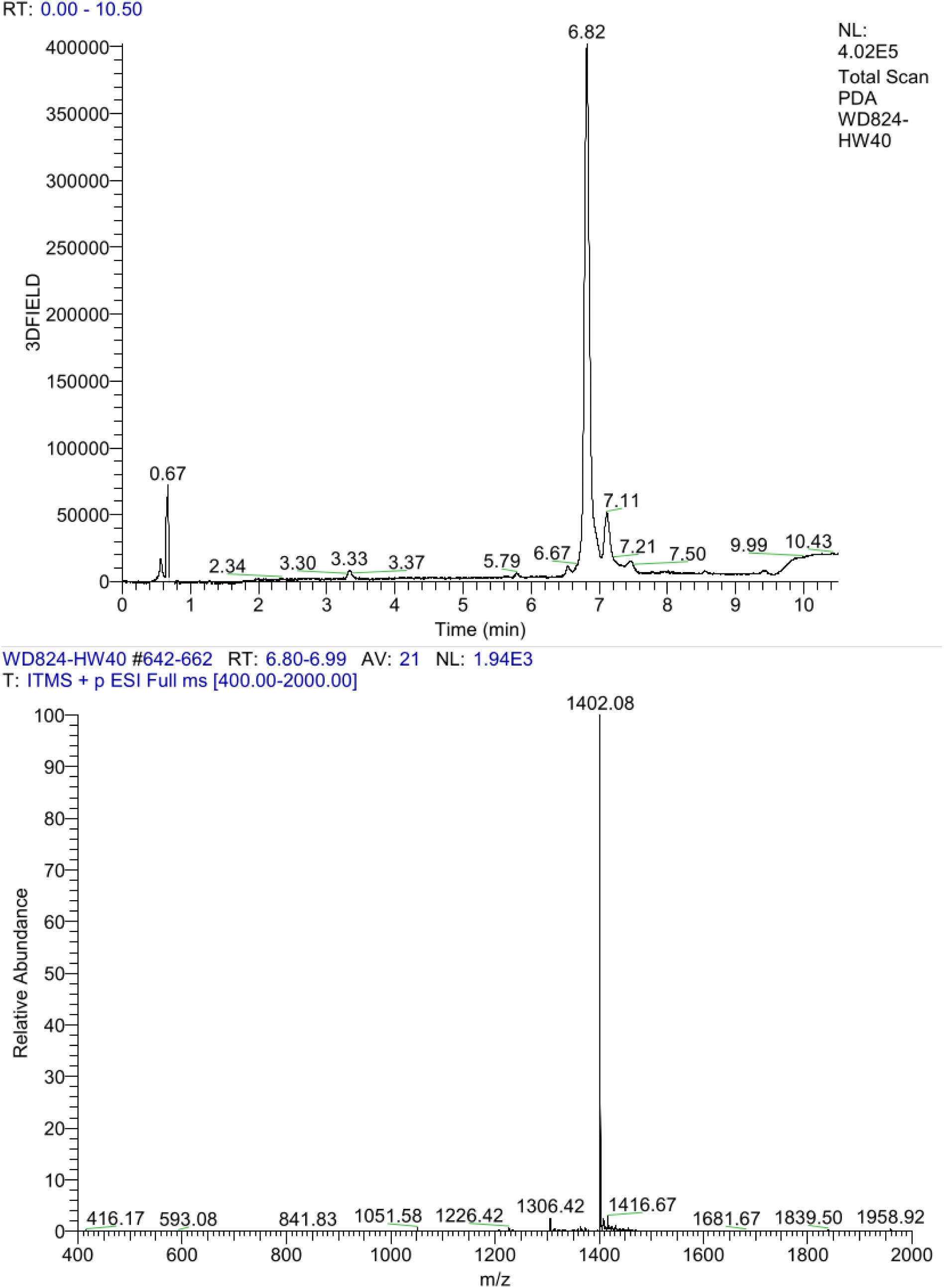

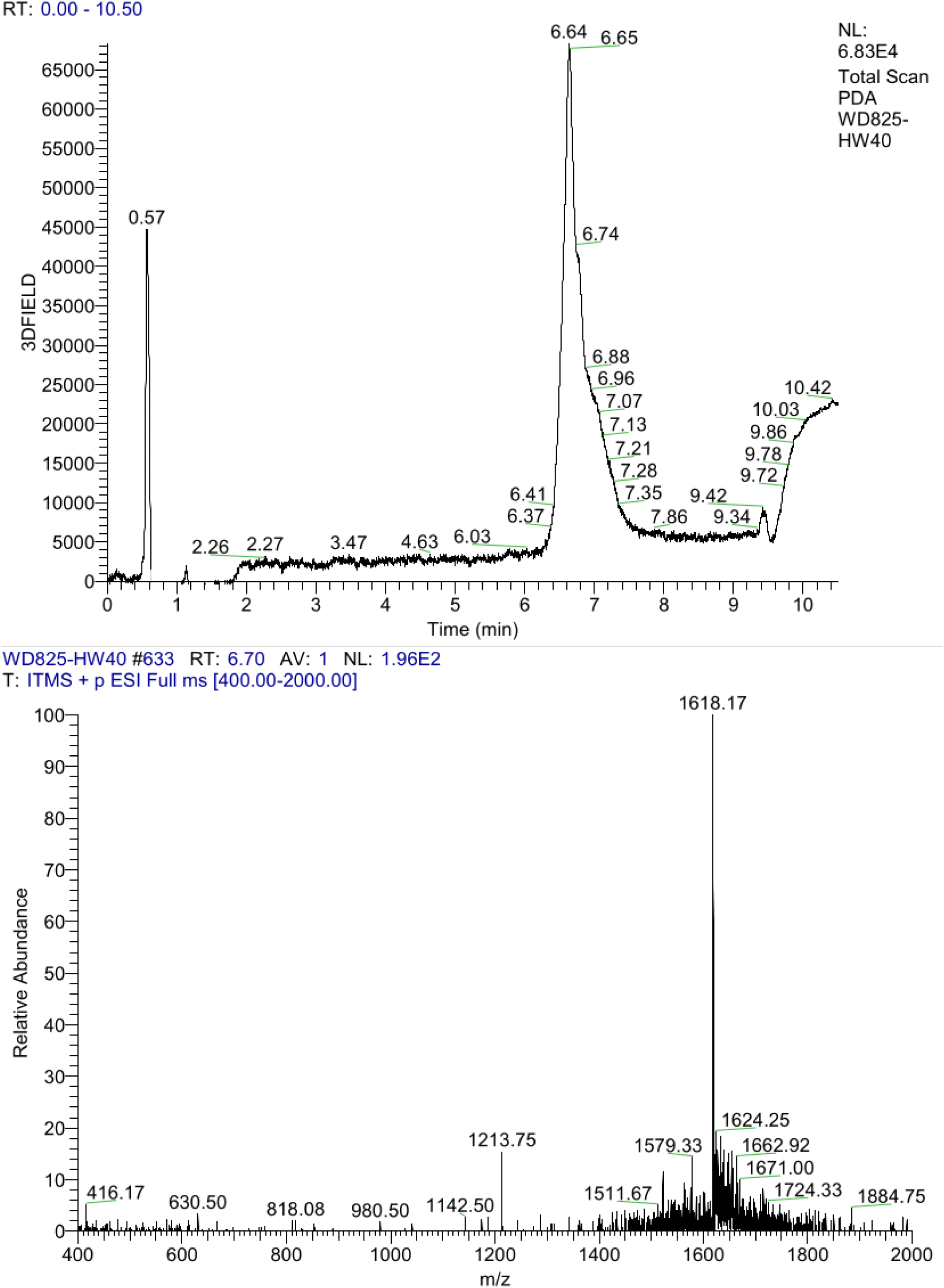

**Ac-Lys(Man)-TEG-Asp-Glu-Val-Ser-Gly-Leu-Glu-Gln-Leu-Glu-Ser-Ile-Ile-Asn-Phe-Glu-Lys-Leu-Ala-Ala-Ala-Ala-Ala-Lys(sCy5)-NH_2_ (1)**

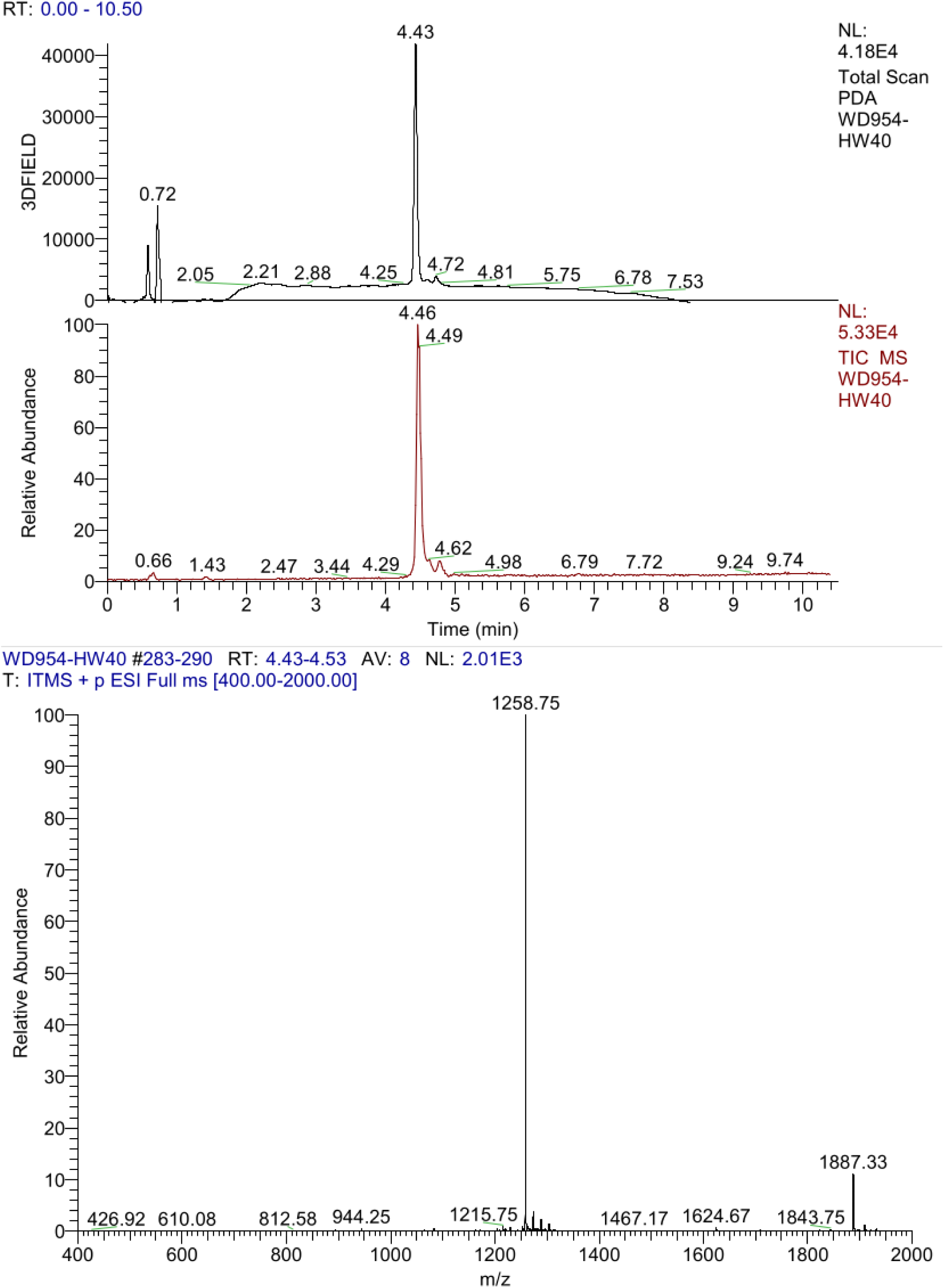

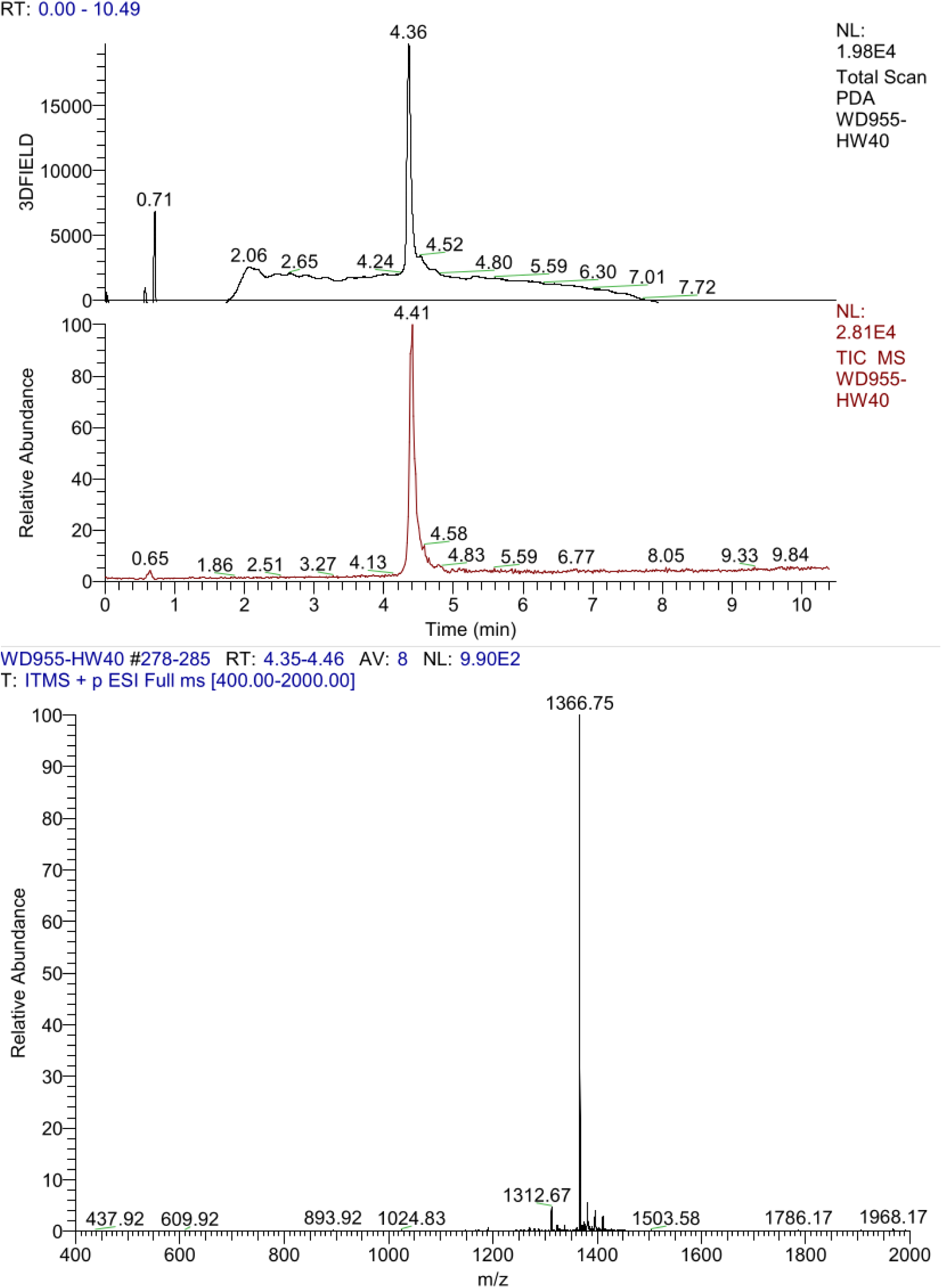

