## Supplementary material for "Quantification of single molecule glycan-mannose receptor binding kinetics on myeloid cells reveals high subcellular binding heterogeneity": supplemnetary manual

GlycoPaint Pipeline Functional description

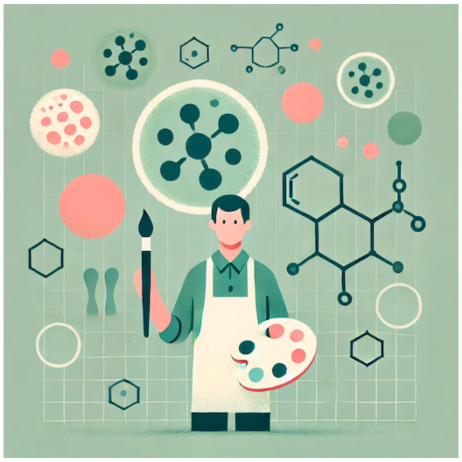

November 2024

### Overview - The Dataflow in the GlycoPaint Pipeline

Cells are observed under a microscope for 100 seconds capturing 2000 images during this interval. Bright spots, which represent fluorophores, are visible in these images. These spots may move and often disappear after a short period. Spots appear because a fluorophore has remained stationary long enough to be detected by the camera. In the GlycoPaint method, spots are interpreted as glycans (with fluorophores attached) binding to lectins on the cell surface. A key challenge is distinguishing these biologically relevant events from non-specific fluorophore sticking.

With the Fiji plugin TrackMate, spots are detected in each frame, and their positions are registered. Tracks are then created by connecting spots across consecutive frames. Each track is characterised by features such as its duration, average x and y positions, and distance travelled. If a lectin-glycan pair were completely stationary, the resulting track would consist of a series of spots in the same location, with no movement. However, lectin-glycan pairs typically move slightly within the cell membrane, resulting in tracks that resemble small 'wriggles'. These tracks are not evenly distributed across the image but are typically concentrated on the cell surface and show heterogeneous patterns within a cell.

A grid is overlaid on the recording, commonly a 20x20 or 30x30 grid, dividing the image into 400 or 900 squares. Dividing the image into squares allows for the investigation of differences between cells or between regions within a cell.

In the GlycoPaint Pipeline, the primary unit of analysis is not an individual recording but a set of recordings grouped into an Experiment. Data processing generates two main datasets:

- **All Squares**: A table containing summarised information on all tracks across all recordings in the experiment.
- **All Tracks**: A table containing detailed information on every track in the experiment.

The two datasets provide the foundation for further analysis.

A lot of data is generated. For an order of magnitude, an average recording may contain around 800,000 spots, from which approximately 50,000 tracks are constructed. A 20x20 or 30x30 grid divides the image into 400 or 900 squares. One of the demo experiments, with 14 recordings and a 20x20 grid, has an All Squares table containing 5600 rows (or 12,600 rows for a 30x30 grid) and the All Tracks table contains nearly 180,000 tracks.

The scope of the GlycoPaint pipeline is to extract information from the Recordings and to extract the maximum amount of meaningful summary information from the images for subsequent processing.

Important features of the pipeline are that the results are fully reproducible, require minimal user intervention and are efficient in terms of processing time and

#### Pipeline Concepts

The Paint Pipeline was developed to analyse (bulk) data from Glyco-PAINT experiments. The interaction of weak-binding ligands to receptors on live cells is studied on varying cell types, ligands and adjuvants. Binding events show up as bright spots under a fluorescent microscope. Analysis of the location and duration of spots reveal information about single binding events.

##### Recording

The central concept in PAINT is the recording of binding events. A recording consists of a brightfield image to identify the position of cells and a multi-frame image in which the spots, indicative of binding events, are recorded. In our experiments, we observe cells for 100 seconds, with 2,000 frames at 50 millisecond intervals.

##### Experiment

A set of recordings is referred to as an experiment. Typically, an experiment is conducted in one day, with up to 100 individual recordings. Normally, several conditions are studied in an experiment and several replicates are recorded for each condition.

##### Project

A project is a set of experiments that together are analysed. Experiments in a project may be replicates of earlier conducted experiments or introduce new combinations of cell types, probes or adjuvants.

**Metadata**

For recordings metadata needs to be provided for the Paint pipeline to work. On top of that, a simple convention for naming the recordings is used. For example, from the name '240116-Ex1-1-A1-1', the following can be derived:

- 240116-Exp-1-A1-1: Part of experiment 240116
- 240116-Exp-1-A1-1: Operating condition 1
- 240116-Exp-1-A1-1: Microscopic plate well A1
- 240116-Exp-1-A1-1: Replicate 1

### Process Flow

#### Overview

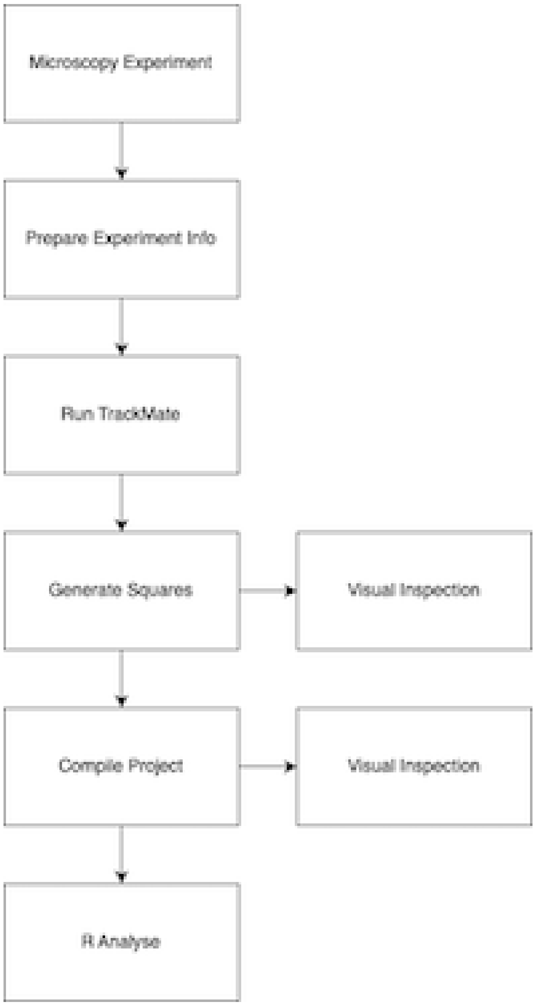


The GlycoPaint process starts with a **microscopy experiment**, in which binding events are observed under a fluorescent microscope. Such an experiment leads to a set of recordings, each consisting of a bright field image and a multi-frame image.

The user subsequently prepares the **Experiment Info** by providing metadata for the recordings. The metadata specifies properties such as Cell Type, Probe Type, Concentration and Adjuvant and provides some necessary processing parameters.

The experiment data is analysed in **TrackMate** which detects spots in each frame of the multi-frame set and connects these spots into tracks where appropriate. A track represents a binding event with the length of the track indicating the duration of the binding event.

With the tracks available, the **Generate Squares** step overlays a fine grid of squares over each recording, establishing the properties of each of the squares, for example, whether the square covers an area of interest or likely background.

**Compile Project** combines all available data into large data tables containing all information of interest.

These tables are used in the **R Analysis** phase where the data is selected, processed and presented in line with research objectives.

**Visual Inspection** of the results is possible at any time after the squares have been generated, both either for recordings in an experiment or for all recordings in a project.

The steps will be described in detail in coming sections, using demo data that can be downloaded from: DOI 10.5281/zenodo.14196381

#### Prepare Experiment Info

The metadata of the experiment is information about the conditions under which each recording is made and recorded in an 'Experiment Info' file. The Paint processing pipeline contains a 'Prepare Experiments Info' utility to prepare this file. The user provides two parameters: the directory that contains the recordings and the location of the Paint Experiment directory where the Experiment Info file (and all subsequent data) will be written.

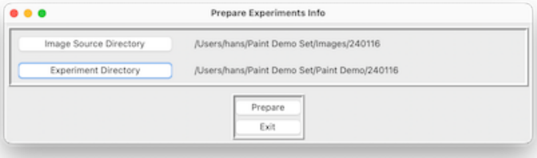


After this has been done for both demo experiments, two directories have been created each with an 'Experiment Info' file.

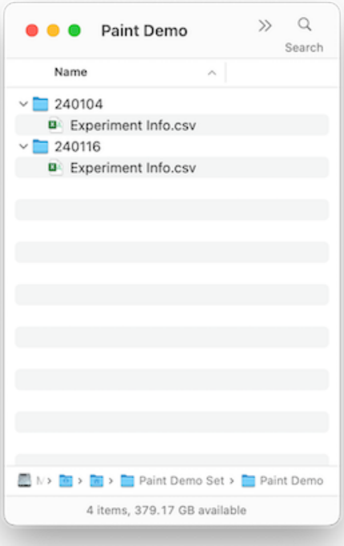


If the Paint file naming convention is used, columns such as Experiment Date, Experiment Name, Condition Nr and Replicate Nr will be filled in automatically, if not the user has to provide this information manually.

| Rec. Name | Exp. Date | Exp. Name | Cond. Nr | Rep. Nr | Probe | Probe Type | Cell Type | Adj | Conc. | Thesh. | Process |
|---|---|---|---|---|---|---|---|---|---|---|---|
| 240116-Exp-1-A1-2 | 240116 | 240116 | 1 | 2 | | | | | | | Yes |
| 240116-Exp-1-A1-3 | 240116 | 240116 | 1 | 3 | | | | | | | Yes |
| 240116-Exp-1-A1-5 | 240116 | 240116 | 1 | 5 | | | | | | | Yes |
| 240116-Exp-2-A2-2 | 240116 | 240116 | 2 | 2 | | | | | | | Yes |
| 240116-Exp-3-A3-2 | 240116 | 240116 | 3 | 2 | | | | | | | Yes |
| 240116-Exp-7-A1-2 | 240116 | 240116 | 7 | 2 | | | | | | | Yes |
| 240116-Exp-7-A1-4 | 240116 | 240116 | 7 | 4 | | | | | | | Yes |
| 240116-Exp-8-A2-2 | 240116 | 240116 | 8 | 2 | | | | | | | Yes |
| 240116-Exp-10-B1-1 | 240116 | 240116 | 10 | 1 | | | | | | | Yes |
| 240116-Exp-10-B1-2 | 240116 | 240116 | 10 | 2 | | | | | | | Yes |
| 240116-Exp-10-B1-3 | 240116 | 240116 | 10 | 3 | | | | | | | Yes |
| 240116-Exp-11-B2-2 | 240116 | 240116 | 11 | 2 | | | | | | | Yes |
| 240116-Exp-12-B3-2 | 240116 | 240116 | 12 | 2 | | | | | | | Yes |
| 240116-Exp-13-A4-2 | 240116 | 240116 | 13 | 2 | | | | | | | Yes |

An example of a fully specified Experiment Info file, with values for Probe, Probe Type, Cell Type, Adjuvant and Concentration provided is shown below. The Threshold parameter is necessary for TrackMate processing (refer to next section). The 'Process' flag indicates whether a recording is processed or ignored.

| Seq. Nr | Rec. Name | Exp. Date | Exp. Name | Cond. Nr | Rep. Nr | Probe | Probe Type | Cell Type | Adj | Conc. | Thesh. | Process |
|---|---|---|---|---|---|---|---|---|---|---|---|---|
| 1 | 240116-Exp-1-A1-2 | 240116 | 240116 | 1 | 2 | 6 Mono | Simple | BMDC | No | 10 | 5 | Yes |
| 2 | 240116-Exp-1-A1-3 | 240116 | 240116 | 1 | 3 | 6 Mono | Simple | BMDC | No | 10 | 10 | Yes |
| 3 | 240116-Exp-1-A1-5 | 240116 | 240116 | 1 | 5 | 6 Mono | Simple | BMDC | No | 10 | 10 | Yes |
| 4 | 240116-Exp-2-A2-2 | 240116 | 240116 | 2 | 2 | 1 Tri | Simple | BMDC | No | 10 | 10 | Yes |
| 5 | 240116-Exp-3-A3-2 | 240116 | 240116 | 3 | 2 | 6 Tri | Simple | BMDC | No | 10 | 10 | Yes |
| 6 | 240116-Exp-7-A1-2 | 240116 | 240116 | 7 | 2 | 6 Mono | Simple | BMDC | MPLA | 10 | 20 | Yes |
| 7 | 240116-Exp-7-A1-4 | 240116 | 240116 | 7 | 4 | 6 Mono | Simple | BMDC | MPLA | 10 | 10 | Yes |
| 8 | 240116-Exp-8-A2-2 | 240116 | 240116 | 8 | 2 | 1 Tri | Simple | BMDC | MPLA | 10 | 10 | Yes |
| 9 | 240116-Exp-10-B1-1 | 240116 | 240116 | 10 | 1 | 6 Mono | Simple | iCD103 | MPLA | 10 | 20 | Yes |
| 10 | 240116-Exp-10-B1-2 | 240116 | 240116 | 10 | 2 | 6 Mono | Simple | iCD103 | MPLA | 10 | 10 | Yes |
| 11 | 240116-Exp-10-B1-3 | 240116 | 240116 | 10 | 3 | 6 Mono | Simple | iCD103 | MPLA | 10 | 10 | Yes |
| 12 | 240116-Exp-11-B2-2 | 240116 | 240116 | 11 | 2 | 1 Tri | Simple | iCD103 | MPLA | 10 | 10 | Yes |
| 13 | 240116-Exp-12-B3-2 | 240116 | 240116 | 12 | 2 | 6 Tri | Simple | iCD103 | MPLA | 10 | 20 | Yes |
| 14 | 240116-Exp-13-A4-2 | 240116 | 240116 | 13 | 2 | 6 Tri | Simple | BMDC | LPS | 10 | 20 | Yes |

#### Run TrackMate

The TrackMate plugin in the Fiji environment is used to analyse the recordings, detect spots and connect spots to tracks where possible.

The Experiment Info ‘Threshold’ parameter determines the spot detection sensitivity. With a low threshold value, even not very well-defined spots are detected. With a high threshold value, poorly defined spots are ignored. Experience indicates that with 1,000,000 plus spots, processing takes very long and does not lead to usable results. The user chooses for each recording a threshold value in an iterative process. The threshold should be set so that the number of spots preferably is in the 300,000 to 800,000 range. A good starting value for the Threshold is 20.

The ‘Run TrackMate’ procedure is started from Fiji by selecting the GlycoPaint section from thr top level menu and from there ‘Run TrackMate’. A dialog box to select the Recordings Directory and Experiment Directory (previously created) is displayed.

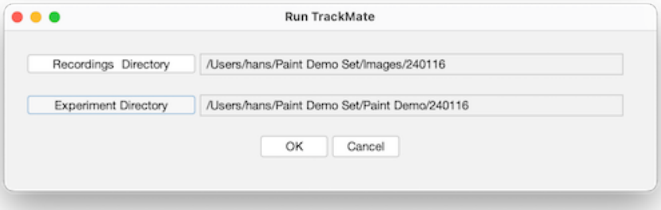


Upon successful processing, the system generates for each recording an image. A representative example of an image is shown below.

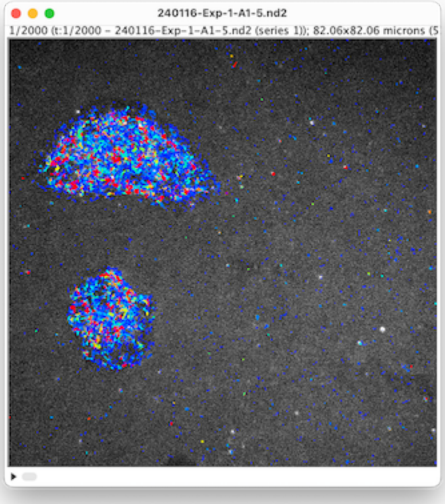


For the set of recordings in the experiment, the system generates one 'All Tracks' and one 'All Recordings' file with attributes such as Nr Spots, Nr Tracks, Run Time and a Time Stamp. The 'All Tracks' file contains, for every track, the average x and y position, and the number of spots and track duration. In addition, the diffusion coefficient (a measure of the displacement from the origin) has been calculated. An example of aan All Tracks file is shown below.

| Unique Key | Ext Recording Name | Track Label | Nr Spots | Track Duration | Track X Location | Track Y Location | Diffusion Coefficien | Square Nr | Label Nr |
|---|---|---|---|---|---|---|---|---|---|
| 240116-Exp-1-A1-2-threshold-5 - 0 | 240116-Exp-1-A1-2-threshold-5 | Track_0 | 233 | 11.75 | 31.36 | 25.48 | 909 | 127 | 9 |
| 240116-Exp-1-A1-2-threshold-5 - 1 | 240116-Exp-1-A1-2-threshold-5 | Track_1 | 816 | 41.11 | 29.21 | 25.76 | 508 | 127 | 9 |
| 240116-Exp-1-A1-2-threshold-5 - 2 | 240116-Exp-1-A1-2-threshold-5 | Track_2 | 7 | 0.3 | 78.7 | 25 | 0 | 139 | |
| 240116-Exp-1-A1-2-threshold-5 - 3 | 240116-Exp-1-A1-2-threshold-5 | Track_3 | 3 | 0.1 | 58.18 | 25.11 | 64 | 134 | |
| 240116-Exp-1-A1-2-threshold-5 - 4 | 240116-Exp-1-A1-2-threshold-5 | Track_4 | 2000 | 99.96 | 26.72 | 26.54 | 849 | 126 | |
| 240116-Exp-1-A1-2-threshold-5 - 5 | 240116-Exp-1-A1-2-threshold-5 | Track_5 | 5 | 0.2 | 72.16 | 25.52 | 353 | 137 | |
| 240116-Exp-1-A1-2-threshold-5 - 6 | 240116-Exp-1-A1-2-threshold-5 | Track_6 | 86 | 4.25 | 26.4 | 23.95 | 541 | 106 | |
| 240116-Exp-1-A1-2-threshold-5 - 7 | 240116-Exp-1-A1-2-threshold-5 | Track_7 | 950 | 47.61 | 24.67 | 23.55 | 35 | 106 | |
| 240116-Exp-1-A1-2-threshold-5 - 8 | 240116-Exp-1-A1-2-threshold-5 | Track_8 | 13 | 0.65 | 32.06 | 24.89 | 64 | 127 | 9 |
| 240116-Exp-1-A1-2-threshold-5 - 9 | 240116-Exp-1-A1-2-threshold-5 | Track_9 | 11 | 0.5 | 55.97 | 24.55 | 77 | 113 | |
| 240116-Exp-1-A1-2-threshold-5 - 10 | 240116-Exp-1-A1-2-threshold-5 | Track_10 | 120 | 6.15 | 24.06 | 25.01 | 2338 | 125 | |
| 240116-Exp-1-A1-2-threshold-5 - 11 | 240116-Exp-1-A1-2-threshold-5 | Track_11 | 24 | 1.15 | 22.63 | 24.24 | 50 | 105 | |
| 240116-Exp-1-A1-2-threshold-5 - 12 | 240116-Exp-1-A1-2-threshold-5 | Track_12 | 29 | 1.5 | 22.12 | 22.56 | 1564 | 105 | |
| 240116-Exp-1-A1-2-threshold-5 - 13 | 240116-Exp-1-A1-2-threshold-5 | Track_13 | 145 | 7.35 | 25.67 | 22.44 | 66 | 106 | |
| 240116-Exp-1-A1-2-threshold-5 - 14 | 240116-Exp-1-A1-2-threshold-5 | Track_14 | 10 | 0.45 | 52.03 | 23.11 | 86 | 112 | |
| 240116-Exp-1-A1-2-threshold-5 - 15 | 240116-Exp-1-A1-2-threshold-5 | Track_15 | 1053 | 52.76 | 27.26 | 22.37 | 264 | 106 | |
| 240116-Exp-1-A1-2-threshold-5 - 16 | 240116-Exp-1-A1-2-threshold-5 | Track_16 | 228 | 11.55 | 29.88 | 22.88 | 593 | 107 | 14 |
| 240116-Exp-1-A1-2-threshold-5 - 17 | 240116-Exp-1-A1-2-threshold-5 | Track_17 | 22 | 1.05 | 27.25 | 22.44 | 6 | 106 | |
| 240116-Exp-1-A1-2-threshold-5 - 18 | 240116-Exp-1-A1-2-threshold-5 | Track_18 | 716 | 36.01 | 21.71 | 20.81 | 174 | 105 | |
| 240116-Exp-1-A1-2-threshold-5 - 19 | 240116-Exp-1-A1-2-threshold-5 | Track_19 | 21 | 1.05 | 42.32 | 20.52 | 13 | 90 | 40 |
| 240116-Exp-1-A1-2-threshold-5 - 20 | 240116-Exp-1-A1-2-threshold-5 | Track_20 | 85 | 4.55 | 27.81 | 20.49 | 729 | 86 | 23 |
| 240116-Exp-1-A1-2-threshold-5 - 21 | 240116-Exp-1-A1-2-threshold-5 | Track_21 | 22 | 1.15 | 23.49 | 21.87 | 526 | 105 | |
| 240116-Exp-1-A1-2-threshold-5 - 22 | 240116-Exp-1-A1-2-threshold-5 | Track_22 | 21 | 1.1 | 11.03 | 21.16 | 122 | 102 | |
| 240116-Exp-1-A1-2-threshold-5 - 23 | 240116-Exp-1-A1-2-threshold-5 | Track_23 | 1361 | 68.16 | 21.78 | 19.4 | 2018 | 85 | 7 |

The Project directory contents just after TrackMate has been run for both the 240104 and 241116 experiment is shown below.

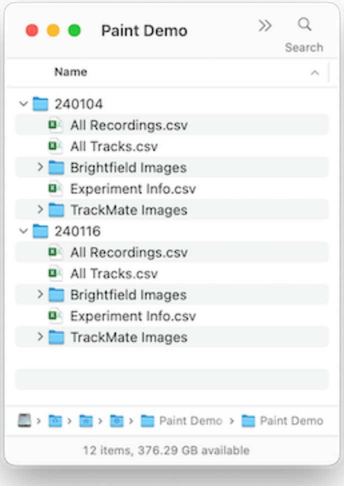


In the GlycoPaint menu a second option is listed: ‘Run TrackMate Batch’. This option allows a number of projects to be processed without user intervention. In this option, a batch file needs to be provided, which is a csv file with 4 columns, specifing the Project directories, the Image Source, the Experiment and a Process flag indicating if the experiment needs to be processed or not. An example is shown below.

| Project | Image Source | Experiment | Process |
|---|---|---|---|
| /Users/hans/Paint Demo Set/Paint Demo | /Users/hans/Paint Demo Set/Images | 240104 | Yes |
| /Users/hans/Paint Demo Set/Paint Demo | /Users/hans/Paint Demo Set/Images | 240116 | Yes |

#### Generate Squares

With the tracks for the experiment available in the ‘All Tracks’ file, the recordings in the Experiment folder (or for all experiments in a Project folder) are analysed by superimposing a grid over the image and determining the characteristics of each square. The user interface to invoke this function is shown below.

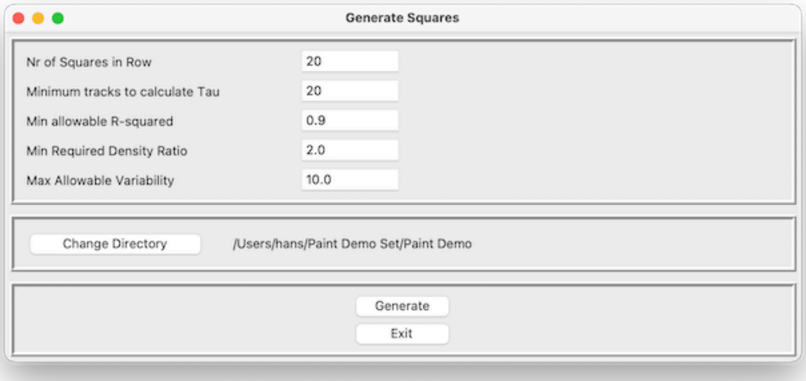


The Number of Squares in a Row (and Column) determines the size of the squares in the grid (typical values are 20 or 30). The number of tracks is then determined for each square. The 10% least dense squares determine a 'background' density. All square densities are compared to this background density. If the density ratio exceeds the 'Minimum Required Density Ratio', squares are interpreted to be sufficiently distinct from the background to be of interest, otherwise they are ignored. The homogeneity of squares is also determined and, if requested, the lack of homogeneity, i.e., the variability, can be limited, excluding squares with too much variability.

A histogram of track durations is generated for each square that meets the specified density and variability criteria. Tau is then calculated through curve fitting. To ensure accurate curve fitting, a minimum number of data points (tracks) is required. The user can define this threshold with the 'Minimum Tracks to Calculate Tau' parameter.

The quality of curve fitting is expressed in an R2 parameter. An R2 value of 1 indicates a perfect fit, values lower than 0.5 that a fit was of low quality. The user-specified 'Min allowable R-squared' parameter sets a limit to acceptable quality of fit.

For all squares in each recording, several attributes are calculated:

- The number of tracks
- The duration of the longest track
- The average duration of the longest 10 tracks
- The total track duration
- The Tau (provided that the squares meet the selection criteria)
- The R2 indicating the quality of the Tau curve fitting

With the 'Generate Squares' function run, the directory structure is shown below (with now additional the All Squares files.).

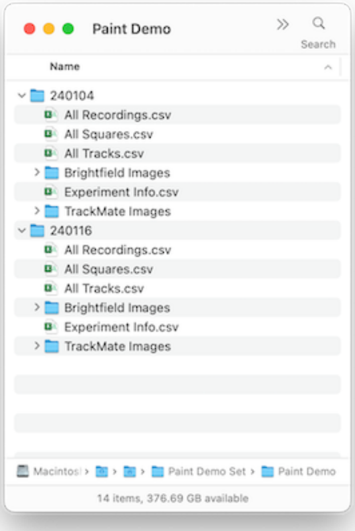


#### Compile Project

Typically, data to be analysed comes from more than one experiment. With the Compile project option, the data from the experiments in the project are compiled and an ‘All Recordings’, ‘All Squares’ and ‘All Tracks’ file on project level is generated.

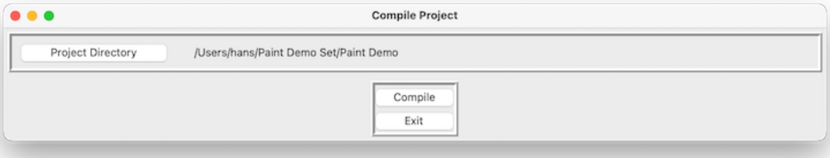

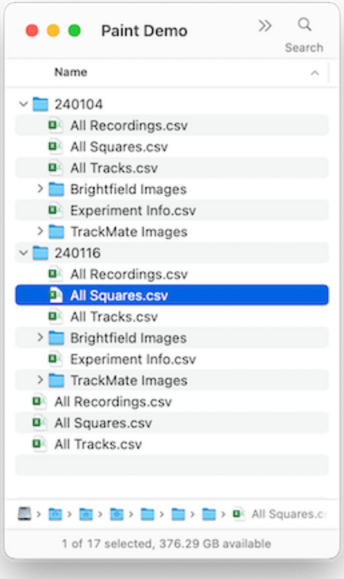


#### Visual Inspection

Once the squares are generated, the results can be reviewed in the Image Viewer. A straightforward dialogue enables the selection of the Project or Experiment directory.

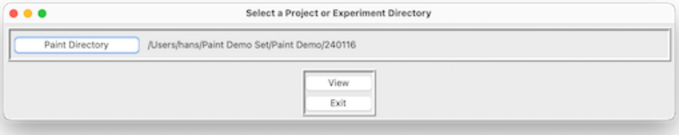


The Viewer dialogue appears, showing on the left the image with tracks and squares, and on the right the corresponding bright field image. Through scroll buttons at the bottom (or the combo box immediately under the ‘Squares’ window), different recordings can be selected.

Metadata of the recording in view is displayed below the Squares image. To better view the underlying cells and tracks, keyboard options allow toggling between showing squares or not (key ‘ts’) or showing numbers or not (key ‘n’).

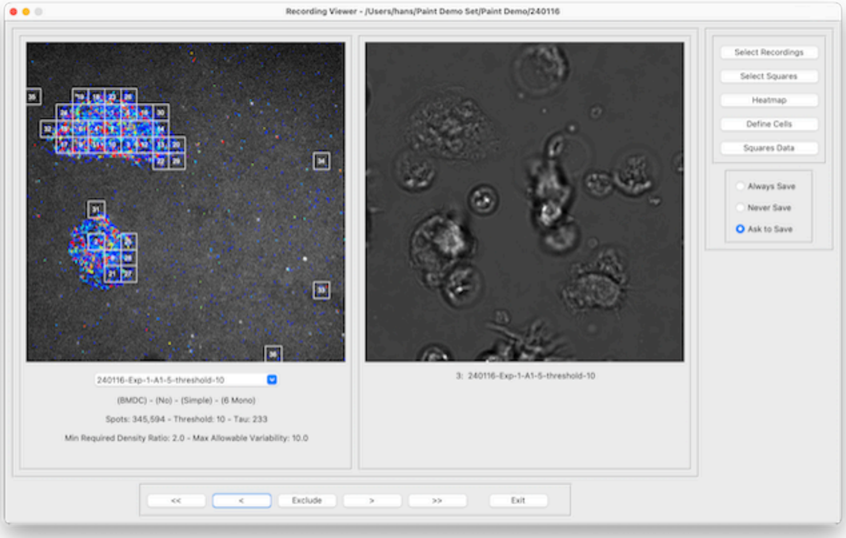


#### Select Recordings

The ‘Select Recordings’ dialogue allows users to narrow down the recordings being viewed. In the example provided, recordings are available for two Cell Types and three Probes. If the user is only interested in BMDC recordings, they can select ‘BMDC,’ then click ‘Filter’ followed by ‘Apply All Filters’ to display only the relevant recordings. The’Reset’ buttons undo a previoulsy made selection. ‘Reset All’ undoes all selections previously made.

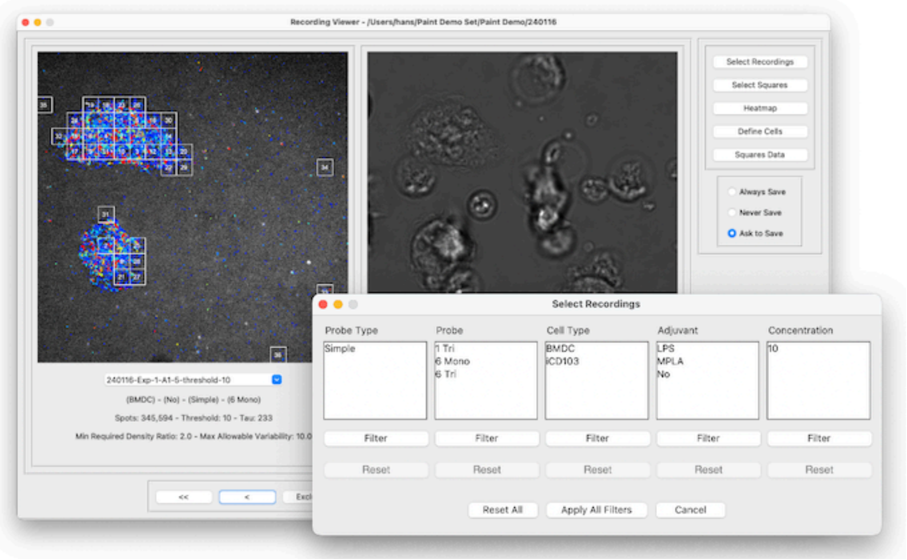

##### Select Squares

The selection of squares of interest for each recording can be adjusted using the Select Squares dialogue. Sliders enable you to set the values for Max Allowable Variability, Min Required Density, and Min and Max Longest Track Duration. Squares that no longer meet the selection criteria will disappear, while those that meet them will be visible.

The ‘Neighbour’ mode defines limitations in the spatial distribution of squares. There are no restrictions in ‘Free’ mode; in ‘Relaxed’ mode, squares must touch at least at the corners; and in ‘Restricted’ mode, they must be adjacent along the edges. The ‘Set All’ button applies the current selection criteria to all recordings in the set.

The Recording Viewer dialogue has three options ‘Always Save’, ‘Never Save’ and ‘Ask to Save’ that determine if the changes are saved or not when the user closes the Recording Viewer.

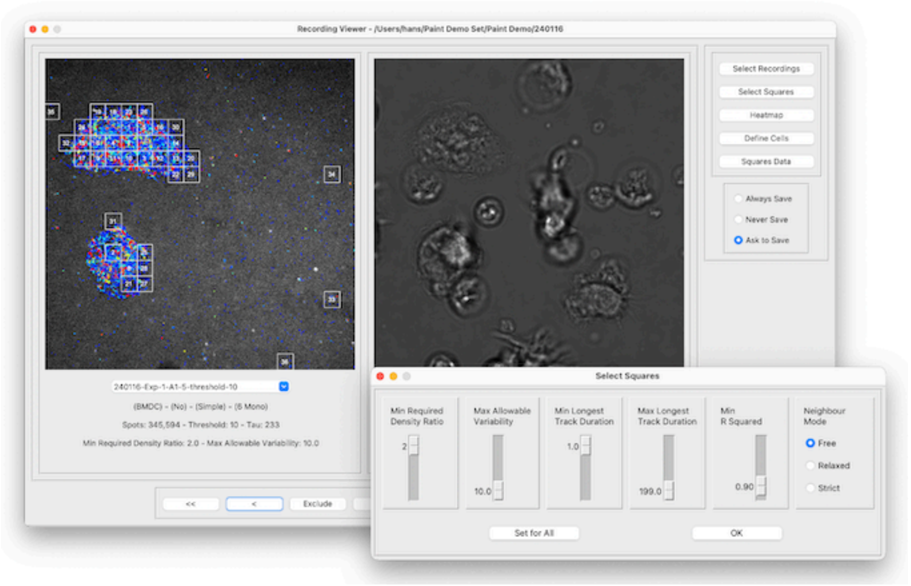


##### Define Cells

In the Define Cells dialogue it is possible to assign squares to cells. The user selects squares by drawing a rectangle with the mouse, selecting a cell (colour) and pressing ‘Assign’.

Any changes made will be saved when the user exits the Recording Viewer, depending on the selected save options.

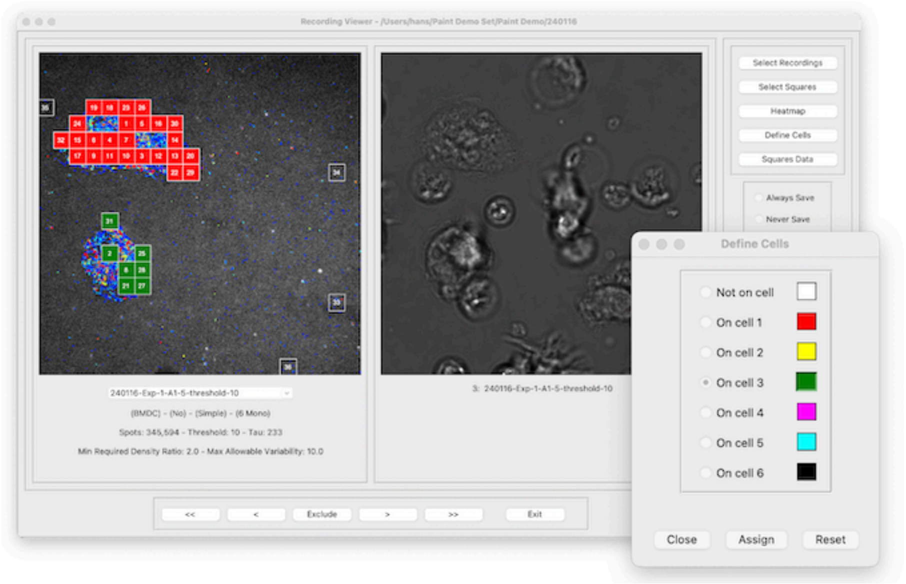


**Show Heatmap**

The Heatmap dialogue allows properties of squares to be viewed relative to each other. Five properties are available: Tau, Density, Average Diffusion Coefficient, Longest Track Duration and Cumulative Track Duration. A colour range of 20 steps is used to display values. The minimum and maximum value is limited to the current recording unless the 'global' option is checked and the minimum and maximum are determined for the current set of recordings. The 'Toggle' function, which can also be invoked by pressing 't', switches between the heatmap and regular display.

Whilst the Heatmap dialogue is displayed, the user can scroll through images.

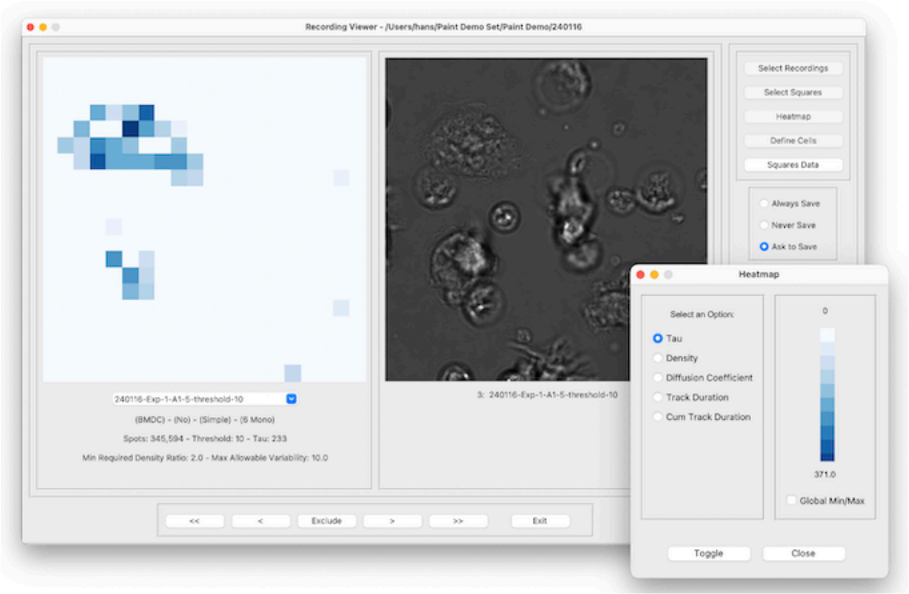


##### Special keybindings

In addition to the interactions that are presented through user interface elements, there are a number of interactions that are bound to keys.

- Key ‘s’ toggles the display of squares.
- Key ‘n’ toggles the display of label numbers.
- Key ‘v’ toggles the display of squares that meet the filtering criteria, but for which no Tau was calculated because the number of tracks is insufficient.
- Key ‘o’ writes the currently displayed images to disc (with and without squares). Images are placed in the Output/Squares directory under the Experiment.
- Key ‘p’ writes all images of the current experiment to disc (with and without squares). Images are placed in the Output/Squares directory under the Experiment.

### Data formats

#### All Recordings Format

Holding information on an experiment. Two versions of All Recordings are shown. Version 1 shows the contents of All Recordings directly after TrackMate had been run and has some extra fields in additiion to the original Experiment Info file. Version 2 shows the contents after Generate Squares has been run. The Tau, Density and R Squared values relate to those calculated for the

Recording. The Tau, Density and R Squared values for individual squares are in the All Squares file.

| | Experiment Info | All Recordings-1 | All Recordings-2 | Comment |
|---|---|---|---|---|
| 1 | Recording Seq. Nr | Recording Seq. Nr | Recording Sequence Nr | Factor for selection |
| 2 | Recording Name | Recording Name | Recording Name | Factor for selection |
| 3 | Experiment Date | Experiment Date | Experiment Date | Factor for selection |
| 4 | Experiment Name | Experiment Name | Experiment Name | Factor for selection |
| 5 | Condition Nr | Condition Nr | Condition Nr | Factor for selection |
| 6 | Replicate Nr | Replicate Nr | Replicate Nr | Factor for selection |
| 7 | Probe | Probe | Probe | Meta Data |
| 8 | Probe Type | Probe Type | Probe Type | Meta Data |
| 9 | Cell Type | Cell Type | Cell Type | Meta Data |
| 10 | Adjuvant | Adjuvant | Adjuvant | Meta Data |
| 11 | Concentration | Concentration | Concentration | Meta Data |
| 12 | Threshold | Threshold | Threshold | TrackMate |
| 13 | Process | Process | Process | TrackMate |
| 14 | | Nr Spots | Nr Spots | TrackMate |
| 15 | | Nr Tracks | Nr Tracks | TrackMate |
| 16 | | Run Time | Run Time | TrackMate |
| 17 | | Ext Rec. Name | Ext Recording Name | Factor for selection |
| 18 | | Recording Size | Recording Size | TrackMate |
| 19 | | Time Stamp | Time Stamp | TrackMate |
| 20 | | | Max Frame Gap | TrackMate |
| 21 | | | Linking Max Distance | TrackMate |
| 22 | | | Gap Closing Max Distance | TrackMate |
| 23 | | | Nr Spots in All Tracks | TrackMate |
| 24 | | | Min Tracks for Tau | User specified |
| 25 | | | Min Required R Squared | User specified |
| 26 | | | Nr of Squares in Row | User specified |

| | Experiment Info | All Recordings-1 | All Recordings-2 | Comment |
|---|---|---|---|---|
| 27 | | | Max Allowable Variability | User specified |
| 28 | | | Min Req. Density Ratio | User specified |
| 29 | | | Exclude | User specified |
| 30 | | | Neighbour Mode | User specified |
| 31 | | | Tau | Calculated |
| 32 | | | Density | Calculated |
| 33 | | | R Squared | Calculated |

#### All Squares format

| | All Squares |
|---|---|
| 1 | Unique Key |
| 2 | Recording Sequence Nr |
| 3 | Ext Recording Name |
| 4 | Experiment Name |
| 5 | Experiment Date |
| 6 | Condition Nr |
| 7 | Replicate Nr |
| 8 | Square Nr |
| 9 | Probe |
| 10 | Probe Type |
| 11 | Cell Type |
| 12 | Adjuvant |
| 13 | Concentration |
| 14 | Threshold |
| 15 | Row Nr |
| 16 | Col Nr |
| 17 | Label Nr |
| 18 | Cell Id |
| 19 | Nr Spots |
| 20 | Nr Tracks |
| 21 | X0 |
| 22 | Y0 |
| 23 | X1 |
| 24 | Y1 |
| 25 | Visible |
| 26 | Variability |
| 27 | Density |
| 28 | Density Ratio |

| | All Squares |
|---|---|
| 29 | Tau |
| 30 | R2 |
| 31 | Diffusion Coefficient |
| 32 | Average Long Track Duration |
| 33 | Max Track Duration |
| 34 | Total Track Duration |

##### All Tracks

Holding information on all tracks in a recording

| | All Tracks |
|---|---|
| 1 | Recording Name |
| 2 | Track Label |
| 3 | Nr Spots |
| 4 | Track Duration |
| 5 | Track X Location |
| 6 | Track Y Location |
| 7 | Diffusion Coefficient |

#### Algorithms

##### Tau

Tau is a measure used to characterise the distribution of track durations. To calculate Tau, a frequency distribution is created from the track durations. The track durations are ordered and fitted with an exponential curve to determine the decay and obtain the Tau value.

All tracks in that square are included in the Tau calculation, but the calculation is performed only if a minimum number of tracks is present. Furthermore, the Tau calculation is only valid if the R2 value meets or exceeds a specified threshold.

To calculate one Tau for the whole Recording, all tracks within squares that meet the specified selection criteria - such as the minimum required density ratio, maximum allowable variability, minimum and maximum track durations, and neighbour state - are considered.

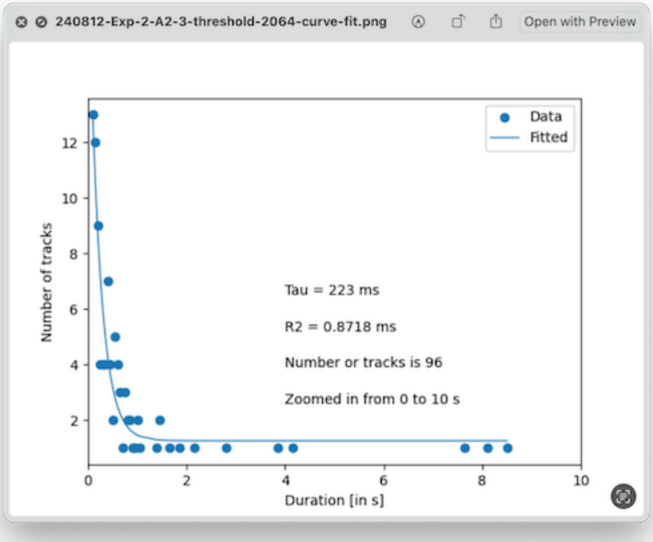


#### Variability

The variability of a square calculation starts with overlaying a grid over the square (i.e., a finer grid over the already exisitng grid) and determining the number of tracks in each grid element.  The variability is then calculated as the quotient of the standard deviation and the mean of the grid track numbers. In the figure below the variability for four fictional squares is shown.

| | | | | | | | | | |
|---|---|---|---|---|---|---|---|---|---|
| 1 | 1 | 1 | 1 | 1 | 10 | 10 | 10 | 10 | 10 |
| 1 | 1 | 1 | 1 | 1 | 10 | 10 | 10 | 10 | 10 |
| 1 | 1 | 1 | 1 | 1 | 10 | 10 | 10 | 10 | 10 |
| 1 | 1 | 1 | 1 | 1 | 10 | 10 | 10 | 10 | 10 |
| 1 | 1 | 1 | 1 | 1 | 10 | 10 | 10 | 10 | 10 |
| 1 | 1 | 1 | 1 | 1 | 10 | 10 | 10 | 10 | 10 |
| 1 | 1 | 1 | 1 | 1 | 10 | 10 | 10 | 10 | 10 |
| 1 | 1 | 1 | 1 | 1 | 10 | 10 | 10 | 10 | 10 |
| 1 | 1 | 1 | 1 | 1 | 10 | 10 | 10 | 10 | 10 |
| 1 | 1 | 1 | 1 | 1 | 10 | 10 | 10 | 10 | 10 |

| | |
|---|---|
| Average | 5.500 |
| Standard Deviation | 4.523 |
| Variability | 0.822 |

| | | | | | | | | | |
|---|---|---|---|---|---|---|---|---|---|
| 10 | 10 | 10 | 10 | 10 | 10 | 10 | 10 | 10 | 10 |
| 10 | 10 | 10 | 10 | 10 | 10 | 10 | 10 | 10 | 10 |
| 10 | 10 | 10 | 10 | 10 | 10 | 10 | 10 | 10 | 10 |
| 10 | 10 | 10 | 10 | 10 | 10 | 10 | 10 | 10 | 10 |
| 10 | 10 | 10 | 10 | 10 | 10 | 10 | 10 | 10 | 10 |
| 1 | 1 | 1 | 1 | 1 | 10 | 10 | 10 | 10 | 10 |
| 1 | 1 | 1 | 1 | 1 | 10 | 10 | 10 | 10 | 10 |
| 1 | 1 | 1 | 1 | 1 | 10 | 10 | 10 | 10 | 10 |
| 1 | 1 | 1 | 1 | 1 | 10 | 10 | 10 | 10 | 10 |
| 1 | 1 | 1 | 1 | 1 | 10 | 10 | 10 | 10 | 10 |

| | |
|---|---|
| Average | 8.000 |
| Standard Deviation | 3.763 |
| Variability | 0.470 |

| | | | | | | | | | |
|---|---|---|---|---|---|---|---|---|---|
| 10 | 10 | 10 | 10 | 10 | 10 | 10 | 10 | 10 | 10 |
| 10 | 10 | 10 | 10 | 10 | 10 | 10 | 10 | 10 | 10 |
| 10 | 10 | 10 | 10 | 10 | 10 | 10 | 10 | 10 | 10 |
| 10 | 10 | 10 | 10 | 10 | 10 | 10 | 10 | 10 | 10 |
| 10 | 10 | 10 | 10 | 10 | 10 | 10 | 10 | 10 | 10 |
| 10 | 10 | 10 | 10 | 10 | 10 | 10 | 10 | 10 | 10 |
| 10 | 10 | 10 | 10 | 10 | 10 | 10 | 10 | 10 | 10 |
| 10 | 10 | 10 | 10 | 10 | 10 | 10 | 10 | 10 | 10 |
| 1 | 1 | 1 | 1 | 1 | 10 | 10 | 10 | 10 | 10 |
| 1 | 1 | 1 | 1 | 1 | 10 | 10 | 10 | 10 | 10 |

| | |
|---|---|
| Average | 9.400 |
| Standard Deviation | 2.258 |
| Variability | 0.240 |

| | | | | | | | | | |
|---|---|---|---|---|---|---|---|---|---|
| 10 | 10 | 10 | 10 | 10 | 10 | 10 | 10 | 10 | 10 |
| 10 | 10 | 10 | 10 | 10 | 10 | 10 | 10 | 10 | 10 |
| 10 | 10 | 10 | 10 | 10 | 10 | 10 | 10 | 10 | 10 |
| 10 | 10 | 10 | 10 | 10 | 10 | 10 | 10 | 10 | 10 |
| 10 | 10 | 10 | 10 | 10 | 10 | 10 | 10 | 10 | 10 |
| 10 | 10 | 10 | 10 | 10 | 10 | 10 | 10 | 10 | 10 |
| 10 | 10 | 10 | 10 | 10 | 10 | 10 | 10 | 10 | 10 |
| 10 | 10 | 10 | 10 | 10 | 10 | 10 | 10 | 10 | 10 |
| 10 | 10 | 10 | 10 | 10 | 10 | 10 | 10 | 10 | 10 |
| 10 | 10 | 10 | 10 | 10 | 10 | 10 | 10 | 10 | 10 |

| | |
|---|---|
| Average | 10.000 |
| Standard Deviation | 0.000 |
| Variability | 0.000 |

#### Square Density

The Square Density is normalised to duration and concentration and is calculated with the following formula:

$$Square\ Density = \frac{Track\ Count}{Area * Duration * Concentration}$$

The area of a square is calculated with the following data: 0.1602804 micrometer per pixel, 512 pixels in a row (or column).

$$Square\ Area = \left(\frac{0.1602804 * 512}{Nr\ of\ Squares\ in\ Row}\right)^2$$

##### Background Density and Density Ratio

The background 'density' represents the average track count of the lowest 10% of squares with a track count greater than zero. This measure helps distinguish squares with high track counts, likely on the cell surface, from those on the glass. The 'Min Required Density Ratio' - calculated by dividing the track count in a square by the background track count per square - determines which squares are considered of interest.

##### Diffusion Coefficient

The diffusion coefficient is calculated for each track in the recording that contains three or more spots, using the following formula. Here, *n* represents the dimensionality (2), and *t* is the time interval over which displacement is measured (0.05 s).

$$MSD = \sum_{i=1}^{nr\ spots} (x_i - x_0)^2 + (y_i - y_0)^2$$

$$Diffusion\ Coefficient = \frac{MSD}{(2\ * n * t)}$$

#### Parameters

The operation of the Paint Pipeline can be tuned with parameters that are kept in 'Paint.json' file.

```
{
    "Paint": {
        "Version": "1.0",
        "Image File Extension": ".nd2",
        "Fiji Path": "/Applications/Fiji.app"
    },
    "User Directories": {
        "Project Directory": "~",
        "Experiment Directory": "~",
        "Images Directory": "~",
        "Level": "Experiment"
    },
    "Generate Squares": {
        "Plot to File": false,
        "Plot Max": 5,
        "Fraction of Squares to Determine Background": 0.1,
        "Exclude zero DC tracks from Tau Calculation": false,
        "Neighbour Mode": "Free",
        "Min Track Duration": 0,
        "Max Track Duration": 1000000,
        "Nr of Squares in Row": 20,
        "Min Tracks to Calculate Tau": 20,
        "Min Allowable R Squared": 0.9,
        "Min Required Density Ratio": 2.0,
        "Max Allowable Variability": 10.0,
        "logging": {
            "level": "INFO",
            "file": "Generate Squares.log"
        }
    },
    "TrackMate": {
        "logging": {
            "level": "INFO",
            "file": "Run Trackmate Batch.log"
        },
        "MAX_FRAME_GAP": 3,
        "LINKING_MAX_DISTANCE": 0.6,
        "GAP_CLOSING_MAX_DISTANCE": 1.2,
        "ALTERNATIVE_LINKING_COST_FACTOR": 1.05,
        "SPLITTING_MAX_DISTANCE": 15.0,
        "ALLOW_GAP_CLOSING": true,
        "ALLOW_TRACK_SPLITTING": false,
        "ALLOW_TRACK_MERGING": false,
        "MERGING_MAX_DISTANCE": 15.0,
        "CUTOFF_PERCENTILE": 0.9,
        "MIN_NR_SPOTS_IN_TRACK": 3,
        "DO_SUBPIXEL_LOCALIZATION": false,
        "RADIUS": 0.5,
        "TARGET_CHANNEL": 1,
        "DO_MEDIAN_FILTERING": false,
        "TRACK_COLOURING": "TRACK_DURATION"
    }
}
```

#### Paint

Two parameters are of interest: - Image File Extension: Specifies the extension of the images generated by the microscope. For example, for Nikon it is '.nb2'. Generally speaking, any tiff-compatible format is suitable.

- Fiji Path: Under normal circumstances, this does not have to be specified, as the software will detect the location of Fiji itself.

#### User Directories

In this section,directories previously specified by the user are stored. so they can be offered as defaults in a nect compinent of the pipeline.

#### Generate Squares

In this section, parameters are stored that are used by the Generate Squares app.

- Five parameters, Nr of Squares in Row, Min Tracks to Calculate Tau, Min Required R Squared, Min Required Density Ratio and Max Allowable Variability, have been entered previously by the user in the Generate Squares dialog and stored so they can be presented in the next invocation of Generate Squares.
- Three other parameters, Neighbour Mode, Min Track Duration and Max Track Duration, provide values that cannot be specified in the user interfaces. The defaukts are chosen so that no squares are eliminated.
- Fraction of Squares to Determine Background: Specifies the fraction of squares that are used to avarege the background.

#### TrackMate

In this section, parameters are stored that are used by TrackMate. The TrackMate documentation pages contain much useful information on TrackMate.

In this section, TrackMate parameters that are important in GlycoPaint are explained. Not parameterised, but important choices for TrackMate are the 'LoG detector' model for spot detection and the 'Simple LAP tracker' model for tracking.

##### Spot Detection

The RADIUS parameter is used during the spot detection phase to specify the size of the spots being tracked. It determines the radius of the detection filter applied to identify objects in the dataset.

The value is dependent on multiple factors, including the microscope, recoding parameters and fluorophore used. For the GlycoPaint experiments we conducted, a RADIUS of 0.5 micrometerm was found to give good results.

If positional accuracy is important, the DO_SUBPIXEL_LOCALIZATION parameter enables or disables the refinement of spot positions to subpixel accuracy. This feature is particularly useful for improving the precision of object tracking when working with high-resolution data or small objects that do not align perfectly with pixel boundaries. For GlycoPaint, positional accuracy is not important and the value of the parameter is set to False to avoid unncessarily increasing processing time.

The DO_MEDIAN_FILTERING parameter determines whether a median filter should be applied to the image during the spot detection phase. This preprocessing step helps reduce noise and enhance the detection of objects, especially in noisy datasets. The proposed value is True.

The TARGET_CHANNEL parameter specifies which image channel is used for spot detection in multi-channel datasets. This is critical when working with datasets containing multiple channels (e.g., fluorescence microscopy) and tracking is needed in only one specific channel. For GlycoPaint the parameter defaults to 1.

**Building Tracks**

TrackMate constructs tracks from spots in subsequent frames. The LINKING_MAX_DISTANCE parameter plays a role in the linking phase of tracking. It sets the maximum spatial distance within which two spots in consecutive frames can be linked as part of the same trajectory. If the distance is exceeded, then the two different tracks are assumed.

If we expect that a lectin cano move a considerable distance in the cellmembrane in subsequent frames (50 ms) apart, a high value is required. The proposed value for LINKING_MAX_DISTANCE: 0.5 micrometerm (which is equal to the expected radius of the spots).

In the case that there are multiple spot candidates for linking to a spot. The ALTERNATIVE_LINKING_COST_FACTOR parameter modifies how TrackMate penalises alternative connections that are less ideal (e.g., farther away). For this parameter the recommendedc value of 1.05 is used.

**Gap Closing**

The MAX_FRAME_GAP parameter is part of the gap-closing step in the tracking process. It determines the maximum number of consecutive frames an object can disappear for and still be linked into the same trajectory if it reappears.

Imagine the following situation. There is a clearly defined track, with spots on proximate positions defined in subsequent frames. However, in some frames a spot is missing. Do we interpret these as distinct tracks (binding events) or do assume that the camera just missed a spot and really this is one long track?

If ALLOW_GAP_CLOSING parameter is set to True, TrackMate will consider if gaps in what is possibly one track should be closed. If the number of consecutive frames where a spot is missing is smaller than MAX_FRAME_GAP and if the distance between the last known and new spot location is smaller than GAP_CLOSING_MAX_DISTANCE, the two tracks are interpreted to be one track with some spots missing, otherwise as separarate tracks.

If we expect that a lectin can release a glycan and immeditialy bind another, a low value of MAX_FRAME_GAP is required.

The suggested value for MAX_FRAME_GAP is 1 frame and for GAP_CLOSING_MAX_DISTANCE 0.5 micrometerm (which is equal to the expected radius of the spots).

##### Track Splitting

Gap splitting concerns the ability for a track to split into two or more tracks during tracking. This commonly used in scenarios like Cell division: when a single cell divides into two daughter cells and branching motion (when a particle splits into multiple trajectories). For the GlycoPaint application this is not relevant and ALLOW_TRACK_SPLITTING is set to False.

##### Track Merging

Track mergings describes the ability for two or more tracks to merge into one during tracking. This occurs in scenarios like cell aggregation (two cells move together and become indistinguishable) or particle collision (two particles collide and continue as one). For the GlycoPaint application this is not relevant and ALLOW_TRACK_MERGING is set to False.

##### GlycoPaint

The MIN_NR_SPOTS_IN_TRACK parameter is specific for GlycoPaint, and with its default setting of 3, ignores the shortest possible tracks, consisting of only 2 spots, as they are considered imaging noise.

The TRACK_COLOURING parameter determines what track characteristic is used for colouring the tracks. Valid choices are specified in the TrackMate documentation. Options currently supported are TRACK_ID and TRACK_DURATION.

#### Paint directories

The Paint pipeline creates a 'Paint' directory in the user's home directory, with two subdirectories: 'Defaults' and 'Logger'.

- In the ‘Defaults’ directory, a ‘Paint.json’ holds the parameters that are used by the various components of the Paint pipeline. For regular use, parameters do not need to be changed, but the option to change is provided (but requires detailed insight into the pipeline’s operation). The parameters are explained in the next section.
- In the ‘Logger’ directory, the system writes log files, that provide information about the progress of operations in the pipeline and if any difficulties encountered. The log files can be statically viewed with a regular text editor or dynamically with the MacOS Console application.

#### Structure of Generate Squares

The core of the data processing takes place in Generate Squares. To facilitate stuyding the cose, some high level pseudo-code is presented hers.

**Generate Squares manin structure**

```
For all the Experiments in the Project (Process Project)
    For all the Recordings in the Experiment (Process Experiment)
        For all the Squares in the Recording (Process Recording)
            Calculate Square properties (Process Square)
```

**Process Project**

```
1. Start project processing and log the project path.
2. List and sort all experiment directories.
3. For each directory:
  - Skip if it is not an experiment directory (e.g. if it is an 'Output' directory).
  - Skip if already processed, unless forced by a paint_force flag.
  - Call process_experiment for each unprocessed experiments.
4. Return the number of experiments processed.
```

**Process Experiment**

```
1. Initialize variables for tracking and logging.
2. Load tracks and recordings data into DataFrames.
3. Validate consistency between tracks and recordings data
4. Log the number of recordings to process.
5. For each recording
   - Retrieve recording details.
   - Call process_recording to process data.
   - Update experiment-level metrics with results from recording.
6. Save updated tracks, recordings, and squares data to files.
7. Log total processing time for the experiment.
```

**Process Recording**

```
1. Initialize processing variables.
```

2. Loop through all squares in the grid:
   - Calculate square coordinates.
   - Filter tracks within square boundaries.
   - Compute metrics (Tau, Density, Variability, etc.).
   - Append square data to the main DataFrame.
3. Compute the density ratio for the squares.
4. Label selected squares and propagate labels to tracks.
5. Calculate the recording level Tau and Density
6. Return processed squares, tracks, and recording-level metrics.

##### Process Square

1. Calculate square boundaries based on grid coordinates.
2. Filter tracks within the square.
3. If no tracks:
   - Assign default values for metrics (e.g., Tau, Density = 0).
4. If tracks exist:
   - Compute metrics (Tau, R-squared, Density, Varia| bility, etc.).
5. Return a dictionary containing square-level data.
